# Traceback of Core Transcription Factors for Soybean Root Growth Maintenance under Water Deficit

**DOI:** 10.1101/2020.03.19.999482

**Authors:** Li Lin, Jan Van de Velde, Na Nguyen, Rick Meyer, Yong-qiang Charles An, Li Song, Babu Valliyodan, Silvas Prince, Jinrong Wan, Mackensie C Murphy, Eiru Kim, Insuk Lee, Genevieve Pentecost, Chengsong Zhu, Garima Kushwaha, Trupti Joshi, Wei Chen, Gunvant Patil, Raymond Mutava, Dong Xu, Klaas Vandepoele, Henry T. Nguyen

## Abstract

Some crops inhibit shoot growth but maintain root growth under water-deficit conditions. Unraveling the molecular mechanisms of root plasticity under water deficit conditions in plants remains a major challenge. We developed an efficient platform for identifying core transcription factors (TFs) that collectively regulate each other and/or themselves in response to water stress, and exploring their interconnected regulatory circuitry involved in root growth maintenance under water deficit in soybean. We performed multi-species phylogenetic footprinting combined with spatial-temporal transcriptome analysis of soybean (Glycine max) roots under water deficit to identify conserved motifs that function in the water-stress response. Using these functional conserved cis-motifs, we applied a new approach to trace back motifs-associated core TFs ingroup as signal mediators, which mediate signaling between abiotic and endogenous stimuli. We integrated a co-functional TF–TF network and conserved motif-centered TF–DNA networks to construct a core TF network defined by mutual cross-regulation among core TFs. We found that core TF ARG (Abscisic acid response element binding factor-like Root Growth regulator) represses BRG (Brassinosteroid enhanced expression-like Root Growth regulator) expression through binding to its promoter at a conserved binding site. ARG and BRG antagonistically regulate Phytochrome-interacting factor-like Root Growth regulator (PRG) and combinatorially regulate some other core TFs. These core TFs form complex regulatory circuits to integrate light and multiple hormone signaling pathways and maintain root growth in response to varying degrees of water stress. Our study provides valuable information to unravel the complicated mechanisms of molecular networks involved in the regulation of root growth under water deficit.

## BACKGROUND

Drought is the primary cause of yield loss in agriculture worldwide. Understanding the development and architecture of roots, as well their plasticity, would pave the way for improving drought tolerance in crops and stabilizing productivity under drought conditions (1). Several agronomic species, such as soybean (*Glycine max*) and maize (*Zea mays*), exhibit an important adaptation to drought stress: substantial primary root elongation is maintained at low water potentials, whereas shoot growth completely stops (2). Unraveling the molecular mechanisms of root plasticity under water deficit conditions in plants remains a major challenge.

It has been demonstrated that ABA regulates root growth under osmotic stress conditions *via* an interacting hormonal network with cytokinin, ethylene and auxin (3). Since plant stress responses must be coordinated with growth and development, there is crosstalk between stress signaling pathways and both hormonal and growth/developmental signaling pathways (4). The integration of plant hormone signaling predominantly occurs at the transcriptional level (5). Transcription factors (TFs), which serve as major regulators of gene expression, play crucial roles in stress responses (6–8). TFs are often sites of signal convergence; signal-regulated TFs act in concert with other context-specific TFs and transcriptional co-regulators to establish sensory transcription regulatory networks in response to abiotic stress (5, 7, 9).

Evidence suggests that multiple drought-responsive TFs interact. For example, AREB1, AREB2 and ABF3 bind to the promoter of *DREB2A* and induce its expression in an ABRE-dependent manner (10). In addition, DREB1A, DREB2A and DREB2C physically interact with AREB/ABF proteins (11), and crosstalk occurs between ABA-dependent and ABA-independent response pathways (7, 12). Furthermore, ABRE sequences are present in the promoter regions of stress-responsive NAC genes in *Arabidopsis thaliana* and rice (13), and NAC016 directly binds to the *AREB1* promoter and represses its expression through a trifurcate feed-forward regulatory loop involving NAP during the drought stress response (14). DREB/CBF binds to a DRE/CRT motif in the *ERF1* promoter under abiotic stress, and also to the GCC box in the promoter of *ERF1* in response to biotic stress, suggesting that ERF1 might integrate ethylene, jasmonate and ABA signaling and play an important role in biotic and abiotic stress responses (15). It has been suggested that a highly interconnected TF gene regulatory network coordinates cell differentiation in the Arabidopsis root (16). A recently constructed TF network provides a roadmap of ABA-elicited transcriptional regulation by 21 ABA-related TFs in Arabidopsis seedlings (17). Despite the central roles of some TFs in water stress responses, the exact regulatory mechanisms of individual TFs and their interactions remain obscure.

Soybean has experienced a relatively recent whole-genome duplication (13 million years ago) (18). Compared to Arabidopsis, soybean has a higher total number of TFs and a greater proportion of some TF families, including some legume-specific subfamilies (18). TFs are often regulated by other TFs and/or by themselves, resulting in interconnected regulatory circuits within a large, complex regulatory network (19). Individual subcircuits within a Gene Regulatory Network (GRN) do not operate in a linear hierarchy but are instead strongly intertwined (20, 21). Some important regulatory information in GRNs is ultimately encoded in individual *cis*-regulatory elements (CREs) that modulate gene activities through their interaction with TFs (20). However, even in well-studied organisms, TF binding site information is available for relatively few TFs, and there are limited DNA binding data for TFs in the majority of eukaryotes (22).

Here, we aimed to identify a group of master TFs that mediate signaling between developmental programs and water stress. These core TFs collectively regulate each other and/or themselves, forming interconnected regulatory circuits to integrate multiple signaling pathway under water deficits in soybean primary root. We developed an efficient platform for identifying core TFs and exploring core regulatory circuitry involved in root growth maintenance under water deficit in soybean using the assembly strategy (Fig. 1 a and b). We examined the spatial-temporal expression of soybean genes under water deficit and identified conserved functional CREs using multi-species phylogenetic footprinting. Since current *cis*-motif datasets are generalized for TF families and cannot associate individual TFs with specific cognate motifs, we thus applied a new approach to trace back core TFs ingroup associated with identified CREs. The regulatory subnetworks defined by mutual cross-regulation among groups of core TFs were further investigated through integration of information about co – functional links of core TFs, TF - DNA interaction, different gene function annotations and gene expression profiles. Finally, based on this network, we prioritized and functionally characterized several key regulators involved in the crosstalk between hormonal signaling and light to help maintain root growth during water deficit.

**Figure 1.**
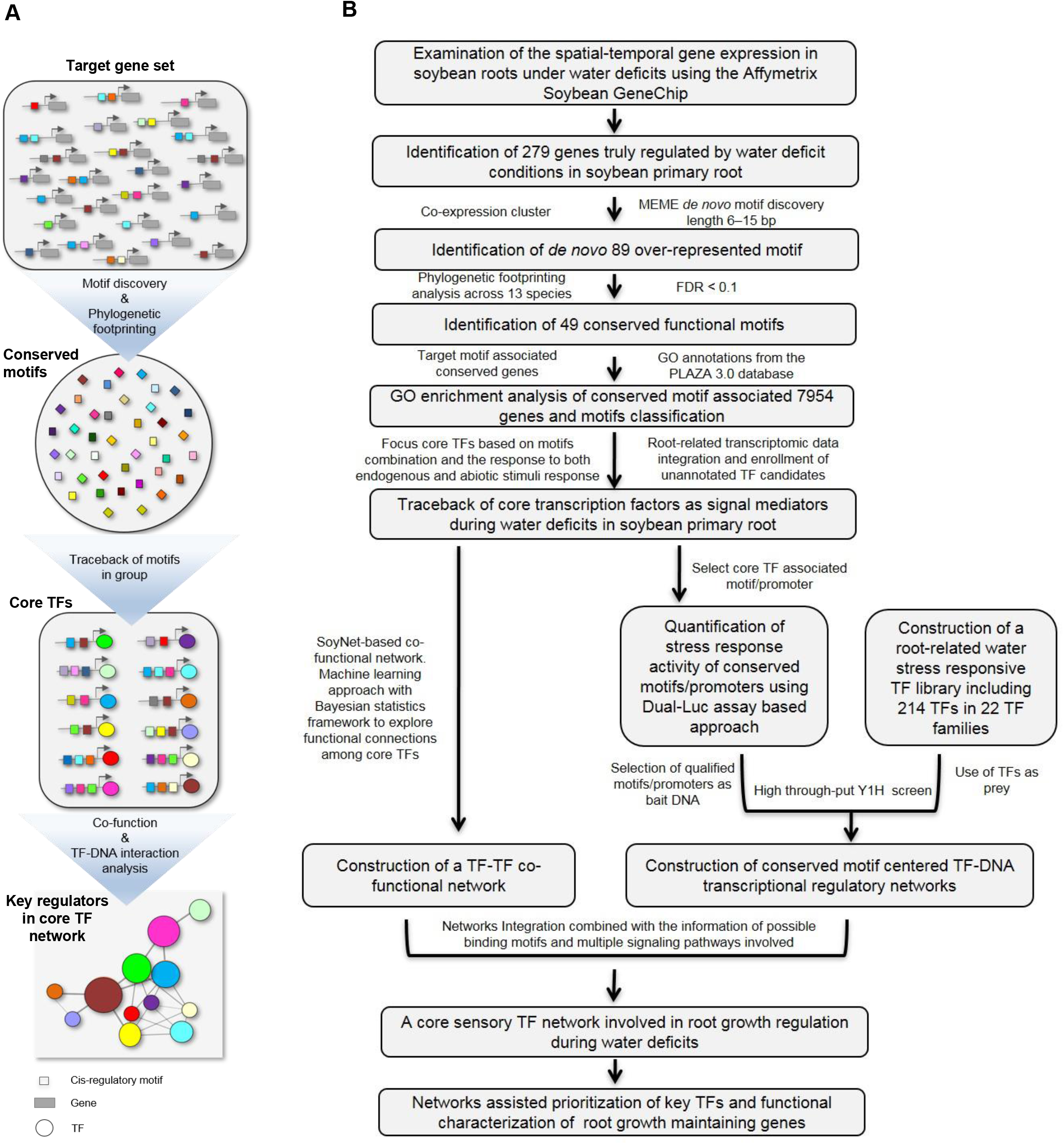
Overview of the workflow used to trace back core TFs and to prioritize key regulators using an assembly strategy. **(A)** An assembly strategy to trace back core TFs in group and prioritize key regulators. **(B)** Overview of the workflow used to trace back core TFs and to prioritize key regulators involved in maintaining root growth under water deficit. MEME (Multiple Expectation maximization for Motif Elicitation): a tool for discovering *de novo* motifs. FDR: false discovery rate. GO: gene ontology. PLAZA: an online platform for plant comparative genomics (http://bioinformatics.psb.ugent.be/plaza/). SoyNet: a database of co-functional networks of *G. max* genes (www.inetbio.org/soynet). Y1H: yeast one-hybrid.

## RESULTS

### Transcript profiling of various regions of soybean roots in response to water stress

Soybean roots continue to elongate when exposed to drought and exhibit region-specific responses to low water potentials. We previously identified three regions of soybean primary roots based on their growth under low water potential −1.6 MPa (23): region 1 (R1) exhibits the same elongation rates under well-watered and water stress conditions; region 2 (R2) exhibits maximum elongation rates in well-watered soil, but progressive deceleration in growth under water stress; and region 3 (R3) exhibits slow growth under well-watered conditions and no growth under water stress.

To explore the molecular mechanism underlying root elongation in response to water stress, we set out to examine the expression of soybean genes in the three different regions (R1, R2 and R3) of the root after 5 h and 48 h (root tip water potential had decreased to approximately −1.6 MPa by this time) of water stress treatment using an Affymetrix Soybean GeneChip containing 37,500 *G. max* probe sets. Transcript expression was profiled in different regions of water-stressed (WS) and well-watered (WW) roots at both 5h and 48h time points. Therefore, the transcripts were divided into five groups according to their expression profiles across the comparisons (Fig. 2a, Table S1): 5hR1WS vs 5hR1WW (D5hR1, D stands for differentiation); 48hR1WS vs 48hR1WW (D48hR1); 5hR2WS vs 5hR2WW (D5hR2); 48hR2WS vs 48hR2WW (D48hR2); 48hR2WS vs 48hR3WW (D48hR2vR3). The last comparison between R2 of water-stressed roots and the zone of growth deceleration in R3 well-watered roots distinguished stress-responsive genes in R2 from those involved in cell maturation. Therefore, we reasoned that the 279 genes with more than two-fold changes in expression detected in both D48hR2 and D48hR2vR3 were important (Fig. 2b; Table S2), because these genes might indeed be regulated by water deficit conditions instead of merely being associated with growth deceleration and tissue maturation closer to the apex. Moreover, a comparison of this group of genes with significantly differentially expressed genes identified in D5hR1, D48hR1 and D5hR2 revealed 49, 69 and 71 genes in common, respectively, as well as 17 genes that were shared by all groups (Fig. 2b). These genes were enriched in categories such as cell wall, hormone metabolism, phenylpropanoids and flavonoids, lignin biosynthesis, peroxidases, TFs and oil biosynthesis. (Fig. 2c and d). It has been reported that lignification increases in R2 and R3 of the elongation zone in soybean roots grown under water deficit conditions, as well as in the equivalent regions of maize primary roots (23). Therefore, these results suggest that the set of 279 genes are likely involved in root elongation in the water stress response and stress adaptation across different root regions.

**Figure 2.**
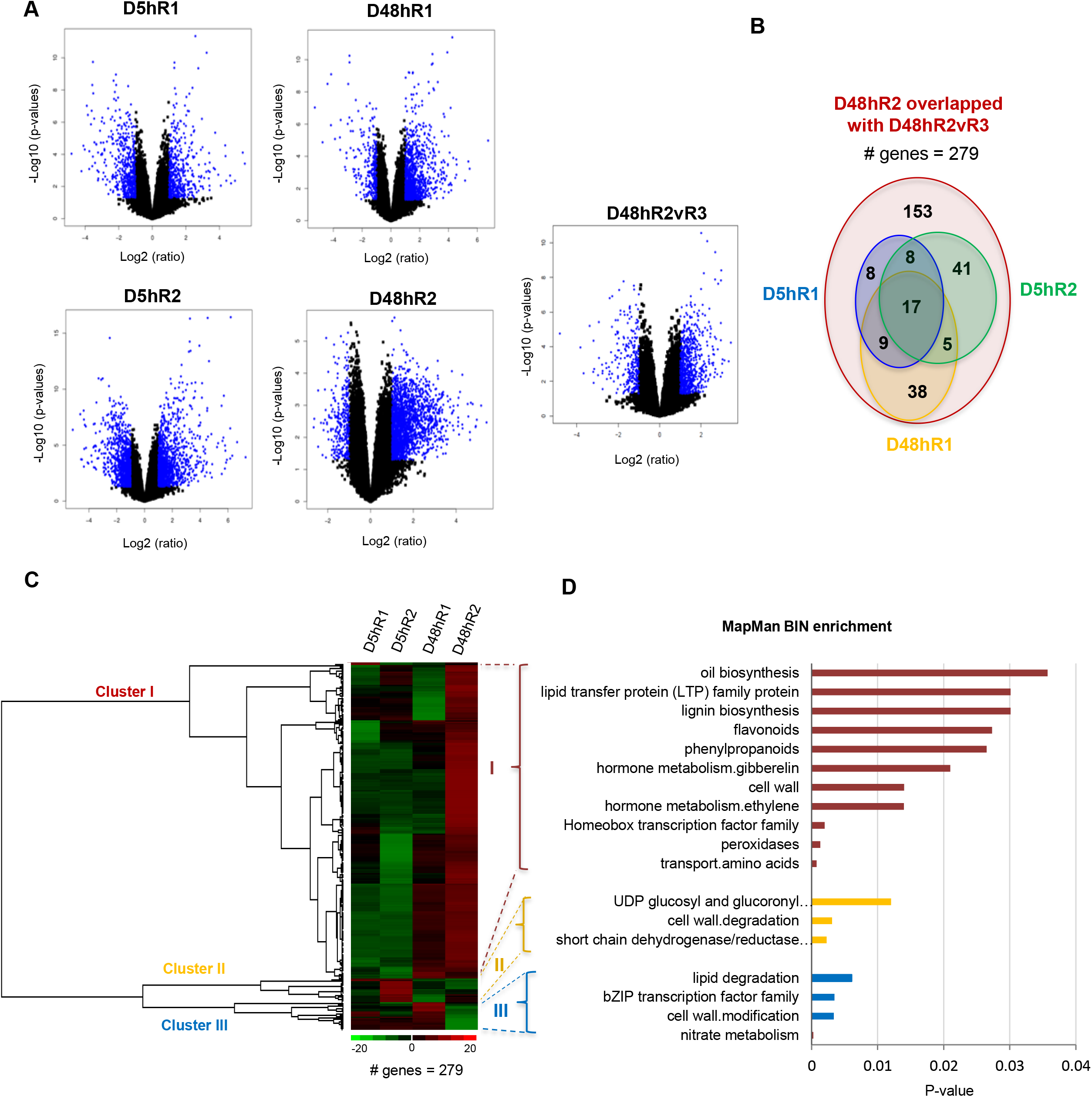
Identification of 279 genes spatially–temporally regulated by water deficit conditions in soybean primary roots. **(A)** Volcano plots showing significant changes in gene expression in soybean root regions R1, R2 and R3 in response to 5 h and 48 h of water stress treatment. The transcripts were divided into five groups according to their expression profiles across the comparisons: D5hR1, D5hR2, D48hR1, D48hR2 and D48hR2vR3. The Benjamini-Hochberg p-values were plotted against the fold change in gene expression for all genes. Log2 fold change (x-axis) versus negative log10 p-values (y-axis). Blue indicates transcripts with fold-changes of ≥ |2.0| and p-values of ≤ 0.05 with respect to the corresponding well-watered control. **(B)** Venn diagram illustrating the number of stress-responsive genes (279 in total) with significantly altered (≥ |2.0| fold, the Benjamini-Hochberg p-values ≤ 0.05) expression in both D48hR2 and 48hR2vR3 (red) and the number of genes shared with D5hR1 (blue), D5hR2 (green) and D48hR1 (yellow). **(C)** Hierarchical tree clustering analysis of 279 differentially expressed genes in different soybean root regions after the primary root was deprived of water for 5 hand 48 h. Hierarchical clustering of the 279 selected genes yielded three gene clusters: I, II and III. **(D)** Overrepresented MapMan bins assigned to the genes in each of three clusters. Enrichment of each category was tested with Fisher’s exact test (p-values ≤ 0.05)

### Identification of conserved functional motifs using multi-species phylogenetic footprinting

We analyzed the 279 water-stress responsive genes using MEME *de novo* motif discovery algorithms and a total of 89 CREs were discovered in the promoter regions of these genes. However, motif-searching tools can generate many false positive candidates, because transcription factor binding sites are often short and typically contain some level of degeneracy in the binding motif (24). We therefore performed multi-species phylogenetic footprinting (25) to identify conserved noncoding sequences in soybean *via* a comparison of soybean with 11 dicot and one monocot genome. We identified 49 of the 89 newly discovered motifs as conserved motifs, with a false discovery rate of <10% (Table S3). We used the STAMP tool (26) to compare the 49 conserved motifs with known *cis*-elements in the plant motif database PLACE (Fig. S1). Several motifs, such as Root Drought-related motif (RDmotif) 18, 12, 5, 65, 22 and 93, showed significant similarities (low E-values) to a G-box motif and ABA responsive elements (ABREs) (PyACGTGG/TC). This result is consistent with the observations that ABA regulates the expression of many genes under osmotic stress conditions and that ABRE is the major *cis*-element for ABA-responsive genes (27). Furthermore, the RDmotif-14 is similar to DREB, another drought-related motif (28).

### Traceback of core TFs ingroup using identified conserved motifs

Extensive reprogramming of temporal and spatial transcription in response to stress occurs *via* the interactions of TFs with a group of *cis*-regulatory motifs. The 49 conserved functional motifs identified in this study could be used to explore how individual network nodes are wired within a root developmental Gene Regulatory Network (GRN) in response to water stress and how subcircuits are connected within the overall network hierarchy. However, the current *cis*-motif datasets are generalized for TF families and cannot associate individual TFs with specific cognate binding motifs. Thus, based on the finding that strong interactions usually occur between individual subcircuits during development and that complex interactions occur among TFs involved in multiple signaling pathways in response to stress (7, 12, 20), we developed a strategy to traceback a set of 49 conserved motif-associated core TFs that mutually regulate each other and/or regulate themselves in response to water stress.

We reasoned that a core TF would have a cognate binding site motif, i.e., one of the 49 conserved motifs, in its own promoter and/or in the promoters of the other core TFs. Therefore, we identified 7954 soybean genes associated with the 49 conserved motifs using the multi-species phylogenetic footprinting approach described above, with each of these conserved genes having at least one of the 49 conserved motifs. Second, we performed GO enrichment analysis of the 7954 soybean genes to obtain a genome-wide overview of their functional roles (Fig. 3a). We classified the conserved motifs and associated the genes into two groups based on GO annotation. In Group I (GI), the conserved genes are primarily involved in plant growth and development. In Group II (GII), the conserved genes are enriched in abiotic stress responses. Although these conserved motifs play multifunctional roles in diverse biological processes, they might work together to process specific responses to water deficit in soybean roots. Interestingly, both GI and GII contain genes enriched in the category “response to endogenous stimulus”. With the aim of revealing important mediators of signaling between development and water stress responses, we focused on a core set of TFs with two properties: 1) encoded by genes associated with both plant growth and development-related motifs and abiotic stress-related motifs; and 2) responsive to both endogenous and abiotic stimuli. Using these two criteria, we identified the set of 35 core TFs by overlapping three groups of TFs (Fig. 3b): 1) GI-Endo: TFs annotated with the GO term “response to endogenous stimulus” in GI; 2) GII-Endo: TFs annotated with the GO term “response to endogenous stimulus” in GII; and 3) GII-Abiotic: TFs annotated with the GO term “response to abiotic stimulus” in GII.

**Figure 3.**
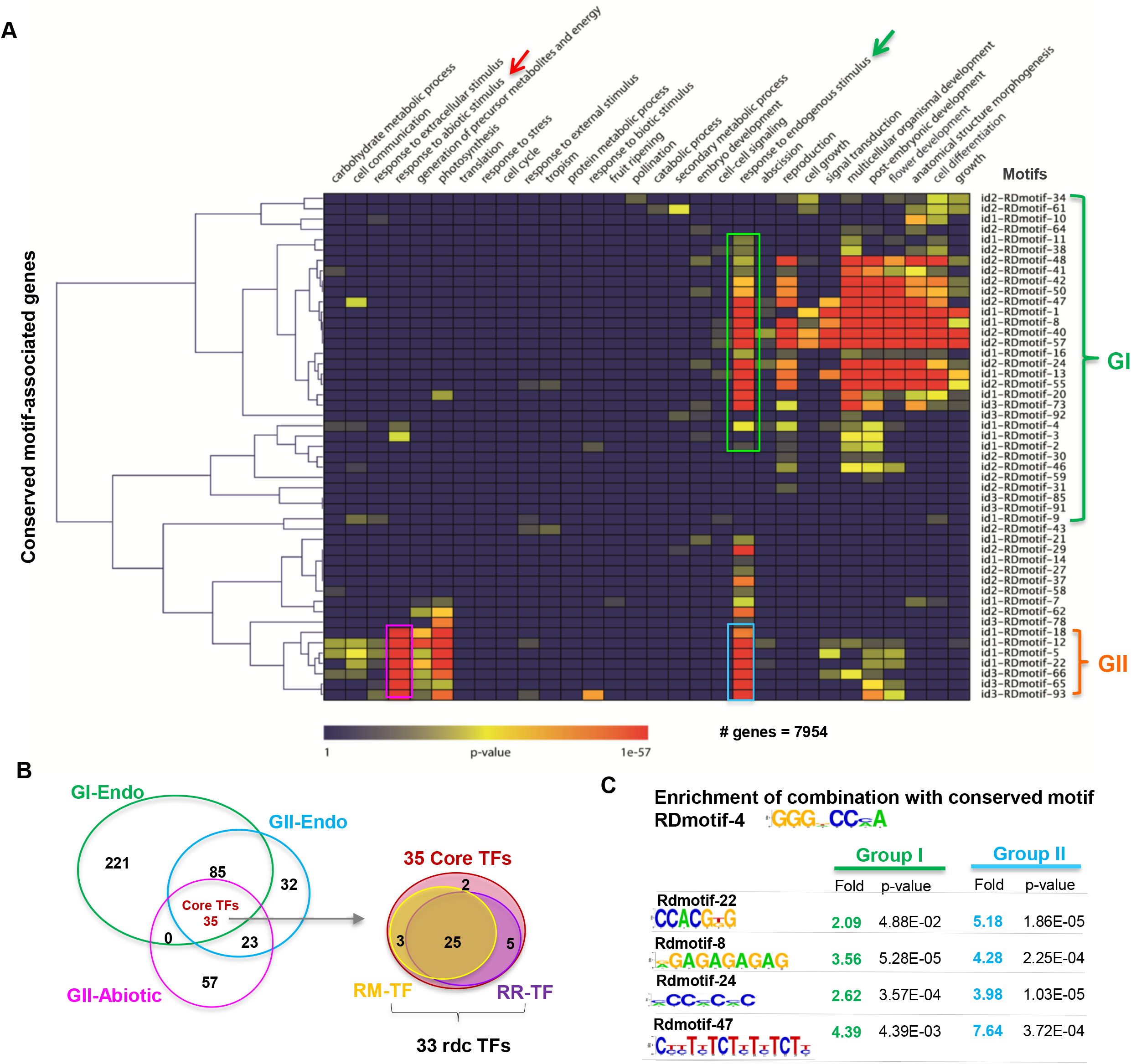
Identification of root-related core TFs that function in the response to endogenous and abiotic stimuli under water-deficit conditions. **(A)** GO enrichment analysis of the 7954 conserved target genes associated with the 49 conserved motifs genome-wide. The vertical axis shows the conserved motifs and the horizontal axis shows the enriched GO terms for the target genes. Conserved motifs and associated target genes were classified into two groups based on GO annotations. Group I (GI): Genes and motifs involved in plant growth and development. Group II (GII): Genes and motifs involved in responses to abiotic stress. Green arrow: response to endogenous stimulus. Red arrow: response to abiotic stimulus. Green box contains genes annotated with the GO term “response to endogenous stimulus” in GI. Blue box contains genes annotated with the GO term “response to endogenous stimulus” in GII. Pink box contains genes annotated with the GO term “response to abiotic stimulus” in GII. GO: gene ontology. **(B)** Venn diagram showing the 35 core TFs that overlap in all three groups of TFs. GI-Endo (green): TFs associated with GO term “response to endogenous stimulus” in Group I. GII-Endo (blue): TFs associated with GO term “response to endogenous stimulus” in Group II. GII-Abiotic (pink): TFs associated with GO term “response to abiotic stimulus”. The expression of 33 rdc TFs among the 35 core TFs is either significantly altered in the root elongation region-specific microarray gene expression profile (RM-TF, yellow) or in the RNA-seq transcriptome profiling data for soybean primary roots under various water deficit conditions (RR-TF, purple). **(C)** Enrichment of the motifs combined with conserved motif RDmotif-4 in the promoters of genes in Group I or Group II with the GO term “response to endogenous stimulus”. Group I represents the genes shown in the green box in (A). Group II represents the genes shown in the blue box in (A).

We verified whether these 35 core TFs indeed respond to water stress in soybean primary root tissue. Since only 37,500 transcripts (approximately 23,787 genes) were examined in our microarray dataset, we introduced our recent RNA-seq-based dataset obtained by transcriptional profiling of soybean primary roots under water deficit (29). This primary root-specific transcriptome in soybean changes in response to very mild stress (VMS), mild stress (MS), severe stress (SS) and recovery from severe stress after re-watering during a long period (21-days) under various water-deficit conditions (Fig. S2a). Remarkably, the expression of 33 of the 35 core TFs significantly differed in at least one of the soybean root-related transcriptome profiles, with 25 TFs in common (Fig. 3b, Fig. S2b). Therefore, we named this set of 33 TFs the root drought-related core TFs (rdcTFs) (Table S4).

The rdcTFs belong to several large multigene families, such as Homeodomain/HOMEOBOX (HB), bHLH, basic leucine zipper (bZIP), APETALA2/ethylene-responsive element binding protein (AP2/EREBP), NAC, Cys2-His2 zinc finger (C2H2), MYB and auxin response factor (ARF) (Table S4). Most rdcTFs contain at least five different conserved motifs (Fig. S2c), pointing to the combinatorial relationships of the 49 conserved motifs in the promoters of core TFs. For example, RDmotif-4 is often present in combination with other motifs, such as RDmotif-22 in GII and RDmotif-8, 24, 47 in GI (Fig. 3c), suggesting that genes containing RDmotif-4 tend to respond to both abiotic stress and endogenous signals. We used this conserved motif association principle to identify additional core TF candidates that have not yet been assigned to GO terms but belong to our set of 279 root elongation-related water stress-responsive genes. Three additional TFs associated with RDmotif-4 and abiotic stress-related RDmotif-22 were found, including two *HAT22* homologs (*Glyma01g40450* and *Glyma17g16930*) and one *BEE* (*BR Enhanced Expression*) gene *Glyma07g10311*. *HAT22* is already known to be involved in an ABA-activated signaling pathway and the response to water deprivation (17). Interestingly, the *BEE* gene *Glyma07g10311* was one of 17 common genes identified in the comparison of the 279 gene set with D5hR1, D48hR1 and D5hR2 (Fig. 2b).

### Co-functional core TF network regulating root growth in response to water stress

Based on SoyBase genome annotation (http://www.soybase.org/genomeannotation/), we found that among the 33 rdcTFs and three additional core TF candidates, 27 TFs are known to be annotated with abiotic stress (e.g., water deprivation, osmotic, salt, cold, superoxide, peroxide, oxidative and heat stress) and 22 TFs are involved in the ABA signaling pathway, indicating that these core TFs regulate abiotic-stress responses in soybean roots (Table S5). These TFs are also responsive to multiple hormones and other signaling pathways, which suggest that these core TFs might orchestrate drought-related responses in soybean roots.

While much is known about the Arabidopsis genome, for other less well-studied plant species, such as soybean, the lack of knowledge regarding gene annotation and interactions severely limits network analysis using gene prioritization (30). Functional gene networks, such as AraNet (31) and RiceNet (32), have proven to be useful for predicting gene functions and for genetically dissecting plant traits. To investigate functional links among our core TFs involved in multiple signaling pathways, we constructed a TF–TF co-functional network based on SoyNet (www.inetbio.org/soynet), a database of co-functional networks of *G. max* genes, which enabled us to obtain the most comprehensive view of soybean pathway systems to date (33). We used the SoyNet to explore functional connections among the 33 rdcTFs and three additional core TFs, finding 24 TFs that were wired to others in co-functional TF–TF network (Fig. S3). These 24 core TFs are close together in the network, with an Area Under ROC Curve score of 0.8362 and p-value 3.7e-102, indicating that these core TFs are well predicted and associated with specific biological processes (Table S6). After analyzing network statistics using Cytoscape (34), we ranked the TFs based on their betweenness centrality (BC) to identify hub genes (Fig. S3 and Table S6). The BC of a node reflects the amount of control that this node exerts over the interactions of other nodes in the network, which helped us target hub TF genes that join different signaling pathways (dense subnetworks), rather than TFs that lie inside a community (35). Network nodes at the intersections of subcircuits are often controlled by signaling interactions (20).

In this TF–TF network, several homologs of core TFs, such as *RVE7*, *RVE8*, *RAP2.4*, *BLH1*, *IAA14*, *VNI2* and *PIF4/PIF5* have been reported to function as mediators of signaling between a developmental program and an environmental cue; these TF are involved in circadian, ABA, auxin, ethylene and phytochrome signaling pathways for cell elongation, xylem formation, root elongation and drought tolerance in Arabidopsis (36–38). To explore the dynamics of these core TFs in response to water stress, we added the induction patterns for each TF using transcriptome profiling data from soybean primary roots under various water-deficit conditions (Fig. S3 and Fig. S2b). Several TF genes serving as hubs in the network, such as *Glyma07g10311, Glyma10g05560, Glyma08g47520* and *Glyma20g35270*, were induced by very mild stress, while most other TFs in the network were induced by mild and/or severe stress conditions, implying that different core TFs play different roles in regulating the response of soybean roots to water stress, depending on its severity, or that they might function in succession as drought conditions progress from very mild to severe through the integration of endogenous and external signals.

### Mapping TF-DNA interactions using a high-throughput Y1H screen and the reconstruction of a core TF network

We hypothesized that core TFs in the co-functional TF network play important roles in integrating multiple abiotic stress and internal signals by regulating each other or themselves *via* binding to conserved motifs located in their promoters. We therefore developed a quantitative motif/promoter analysis platform to confirm the stress-response activity of the conserved motifs and their associated promoters. We also performed a high-throughput Y1H assay to measure TF–DNA interactions to verify the binding of the TFs.

We selected a hub gene, *Glyma07g10311,* a core TF from the 279 genes set, one of 17 genes that were shared by all groups of significantly differentially expressed genes (Fig. 2b), and also one of three additional TFs associated with RDmotif-4 tending to respond to both abiotic stress and endogenous signals, having the most connections with the other core TFs involved in different signaling pathways within the network, for further investigation: we named this gene *BEE-like Root Growth regulator* (*BRG*) (Fig. S3). Phylogenetic analysis showed that the combination of motif RDmotif-22 and RDmotif-4 located in the *BRG* promoter is also found in most homologous genes across 13 species (Fig. 4a). RDmotif-22 is a G-box-type conserved motif that represents the conserved ABRE site gcCACGTGgc. Two phytochrome A-induced motifs, SORLIP1AT and SORLIP2AT are usually over-represented among root-specific genes (39). Another motif, CCAAT/ATTGG, was suggested binding site of NF-YB factors (40). To quantitatively characterize these motifs and their associated promoters, we constructed a single plasmid, pLL-QMP, harboring two reporter genes, firefly luciferase (*LUC*) and *Renilla* luciferase (*REN*) (Fig. 4b). A promoter region or motif repeats of interest was inserted before the minimal promoter to form a synthetic promoter to drive the expression of *LUC* (Fig. S4). In this experiment, five different types of G-box motifs from GII and six representative motifs from GI were selected for analysis based on their inducibility by 100 μM ABA, 150 mM NaCl2 or 16% PEG treatment. Considering the degeneration of motif expression, each motif was designed based on the most frequent occurrence pattern in our root-specific 279 gene set (Fig. 4c, Table S7). The *BRG* promoter, which contains a motif named Gbox-I-gc, a representative RDmotif22 motif, was strongly induced by both salt and ABA. However, other types of G-boxes, such as Gbox-I-ac, Gbox-I-tc, h-Gbox and their associated promoters PROM-*Glyma11g27480*, PROM-*Glyma06g03450* and PROM-*Glyma12g33600,* showed little or no induction by ABA or salt (Fig. 4d). In addition, selected representative motifs from Group I, such as dASE, dGA, dGT, dTG, dGAGA and dACAC (related conserved motifs listed in Table S7), exhibited different levels of induction by ABA, PEG and salt (Fig. 4e). Overall, this quantification pipeline provides a simple method to determine the activities of various motifs/promoters in response to different stimuli.

**Figure 4.**
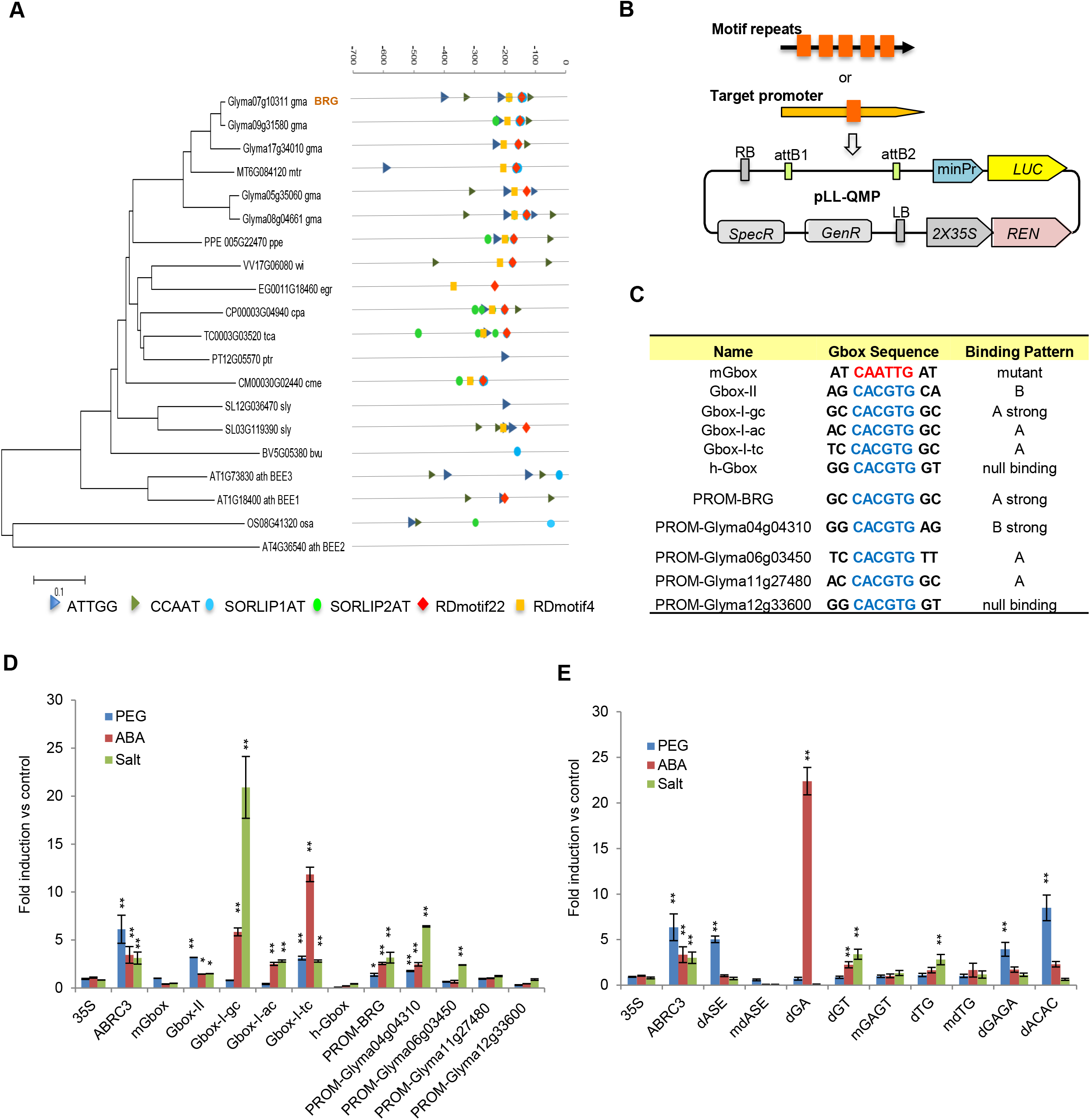
Quantification of stress response activity of the conserved motifs and promoters. **(A)** Phylogenetic tree of *BRG* genes across 13 species, and conserved motif pattern in the promoters of *BRG* and homologous genes showing that RDmotif-4, NF-YB binding sites and other binding site motifs are collocated with RDmotif-22. **(B)** The basic vector pLL-QMP used to quantify the activity of the promoter/motif. **(C)** The G-box motifs and promoter fragments examined. The *BRG* promoter contains motif Gbox-I-gc, a representative RDmotif22. **(D)** Stress response activity of the G-box motif and its associated promoter fragments. The CaMV *35S* and *ABRC3* promoter was used as a negative and positive control, respectively. **(E)** Stress response activity of selected conserved motifs from GII. (D)–(E) The comparisons are between the CaMV *35S* promoter control and the motifs/promoter fragments under the same treatment (mean ±SE; n=5). Significance relative to control was determined by two-tailed Welch’s *t*-test: single and double asterisks denote significance level at P < 0.01 and P < 0.001, respectively.

To identify TFs that potentially bind to the selected motifs and promoters, we constructed a representative drought-related TF library from soybean roots consisting of 214 TFs with more than two-fold differences in expression in response to water stress from our soybean root elongation microarray dataset, (Table S8). The final TF library contained 22 TF families, including some well-known drought-related TFs or their homologs from the MYB, bZIP, AP2, NAC, bHLH and WRKY families (Fig. 5a). We used a conserved motif-centered Y1H platform to identify TFs that interact with the selected conserved motifs and their associated promoters. (Fig. 5b). In total, 70 TF–DNA interactions involving 12 different TFs were identified for the motif Gbox-I-gc group (Fig. 5c and Table S9). Moreover, 15 TF-DNA interactions involving 12 TFs were identified for the dASE motif (a representative of RDmotif-10), an ABA-independent osmotic stress-responsive motif (Fig. 5d). Notably, the Gbox-I-gc motif is preferentially bound by bZIP (*Glyma06g04353*), bHLH (*Glyma10g28290*) and NAC (*Glyma06g21020, Glyma08g18470* and *Glyma09g37050)* TFs, whereas the dASE motif is bound by HB (*Glyma02g04550, Glyma08g20170* and *Glyma03g36070*) TFs. In the Gbox-I-gc motif-centered network, six promoter fragments were classified into three groups based on their preferred binding TFs. Compared with the TFs that bind to mutant G-box (mGbox) and other negative promoters, the bZIP gene *Glyma06g04353* (homolog of *ABF2*), encoding a binding TF shared by motifs Gbox-I-gc, PROM-*BRG* and PROM- *Glyma04g04310*, was characterized as a TF that targets the *BRG* promoter. Thus, based on the binding specificity of PROM-*BRG, Glyma06g04353* and the three other TFs were selected as the candidate regulators of *BRG* for further network analysis. We named these genes based on their homology: *ARG* (*Glyma06g04353*, **A**BRE binding factor-like **R**oot **G**rowth regulator); *PRG* (*Glyma10g28290*, **P**hytochrome-interacting factor-like **R**oot Growth regulator); *NRG* (*Glyma09g01650*, **N**F-YB-like **R**oot **G**rowth regulator); and *HRG* (*Glyma11g03850*, **H**AT-like **R**oot **G**rowth regulator).

**Figure 5.**
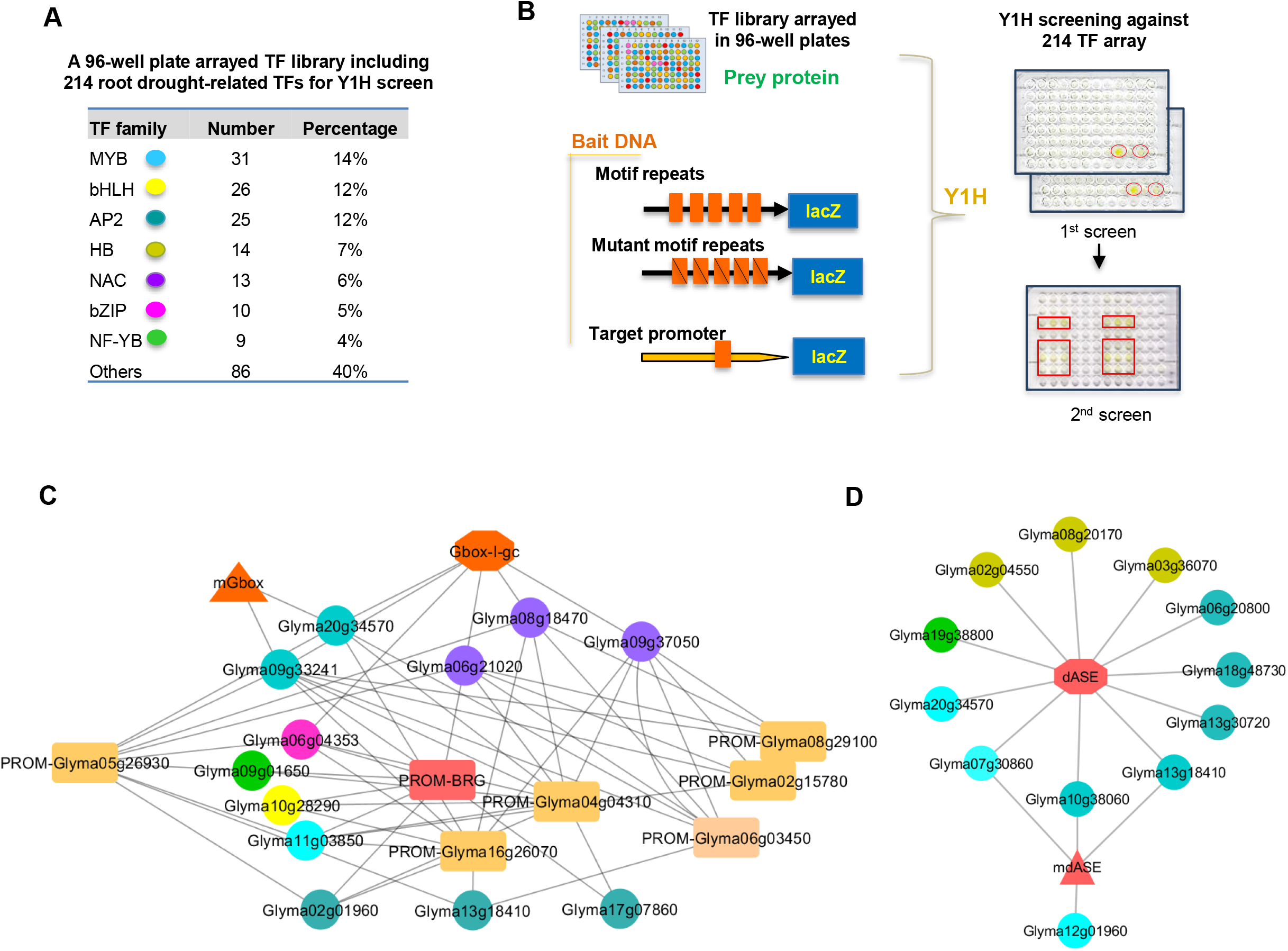
Construction of conserved motif-centered TF-DNA interaction networks. **(A)** A drought-related transcription factor library consisting of 214 TFs in 22 TF families, with a color-coded legend. **(B)** Schematic diagram of the conserved motif-centered Y1H assays. Bait DNAs included conserved motif repeats of interest, mutant conserved motif repeats and associated promoter fragments. Each TF was tested in quadruplicate. Red boxes show positive interactions. **(C)** Conserved RDmotif22-centered TF-DNA interaction network. Gbox-I-gc represents RDmotif22. mGbox represents the mutated G-box. PROM-BRG represents the RDmotif22-associated promoter fragment of *BRG*. PROM-Glyma06g03450 are negative binding promoters. In the network, six promoter fragments were classified into different groups based on their preferred binding TFs. **(D)** Conserved dASE motif-centered TF-DNA interaction network. mdASE represents the mutated dASE motif. (C)–(D) Octagon: conserved motif. Triangle: mutated conserved motif. Rectangle: promoter fragments. Circle: interacting TF. The color of each oval represents the same TF family as in (A).

After exploring the co-functional connections of the four candidate regulators with the other core TFs in SoyNet and integrating TF-DNA interactions into the co-functional TF–TF network, we re-constructed a core TF network to visualize possible mutual regulation among core TFs involved in multiple signaling pathways that function under water deficit in soybean roots (Fig. 6, Table S10). The most highly ranked gene, *BRG*, is connected with 10 TFs. We discovered that *ARG* is the paralog of *Glyma04g04170*, and both genes respond to ABA stimulus and water deprivation, while the two other genes that respond to water deficit, *Glyma04g11290* and *Glym06g1101,* homologs of *RAP2.4*, are upregulated by cytokinin and ethylene. *HRG* responds to ABA and auxin stimuli and is downregulated under salt and drought stress in soybean (41). *PRG* is a GA-responsive gene whose homolog, *Glyma02g45150* functions in gravitropism responses to light. A homolog of *RVE7* (*Glyma16g34340*) induced by ethylene stimulus and salt stress links *BRG* with three other circadian rhythm-related genes, *Glyma10g05560*, *Glyma07g05410* and *Glyma16g01980*. Moreover, *Glyma08g47520* (homolog of *VNI2*, a transcriptional repressor of the master regulator VND7 during xylem vessel development), which responds to both ABA and BR, connects *BRG* with two auxin-responsive genes and four *BLH1*-like genes. Interestingly, these four *BLH1-*like genes respond to multiple hormones and abiotic stress, such as ABA, auxin, salicylic acid, ethylene, superoxide and salt stress. In addition, *Glyma03g36070*, one of the BLH1-like genes in the network, has a dASE motif site in its own promoter, as do the 11 other core TFs. This observation suggests that *Glyma03g36070* not only binds to the dASE site in its own gene and regulates its own expression, but it may also bind to dASE sites in many other core TFs in the network and regulate their expression. It has been reported that BLH1 itself enhances *BLH1* expression (42), suggesting that the dASE motif site may participate in BLH1 self-regulation. RDmotif22 motif is associated with 10 core TFs besides *BRG*, which was already shown to be bound by ARG in our Y1H assays. The link between *ARG* and *NRG* in the network suggests that these two TFs might function together by binding the RDmotif22 and CCAAT in the *BRG* promoter. The detection of *cis*-regulatory motifs implies that the core TFs are heavily intertwined to ensure mutual regulation to coordinate the response to water deficit. *ARG*, *NRG* and *PRG* were upregulated under mild and severe water stress conditions in soybean roots, whereas the auxin signaling-related *HRG* gene was upregulated under very mild water stress conditions and then downregulated, which is similar to the expression patterns of two IAA genes (*Glyma20g35270* and *Glyma08g22190*) in the network. *BRG* was upregulated under very mild water stress conditions and during the water recovery stage, a pattern unlike that of most of its associated genes in the network, suggesting that BRG might function as a repressor and/or be repressed by some of its associated TFs through a tightly controlled network for the sequential activation of core TFs under varying water deficit conditions. Four candidate regulators of BRG might also contribute to the integration of multiple signaling pathways to help regulate root growth in response to water stress.

**Figure 6.**
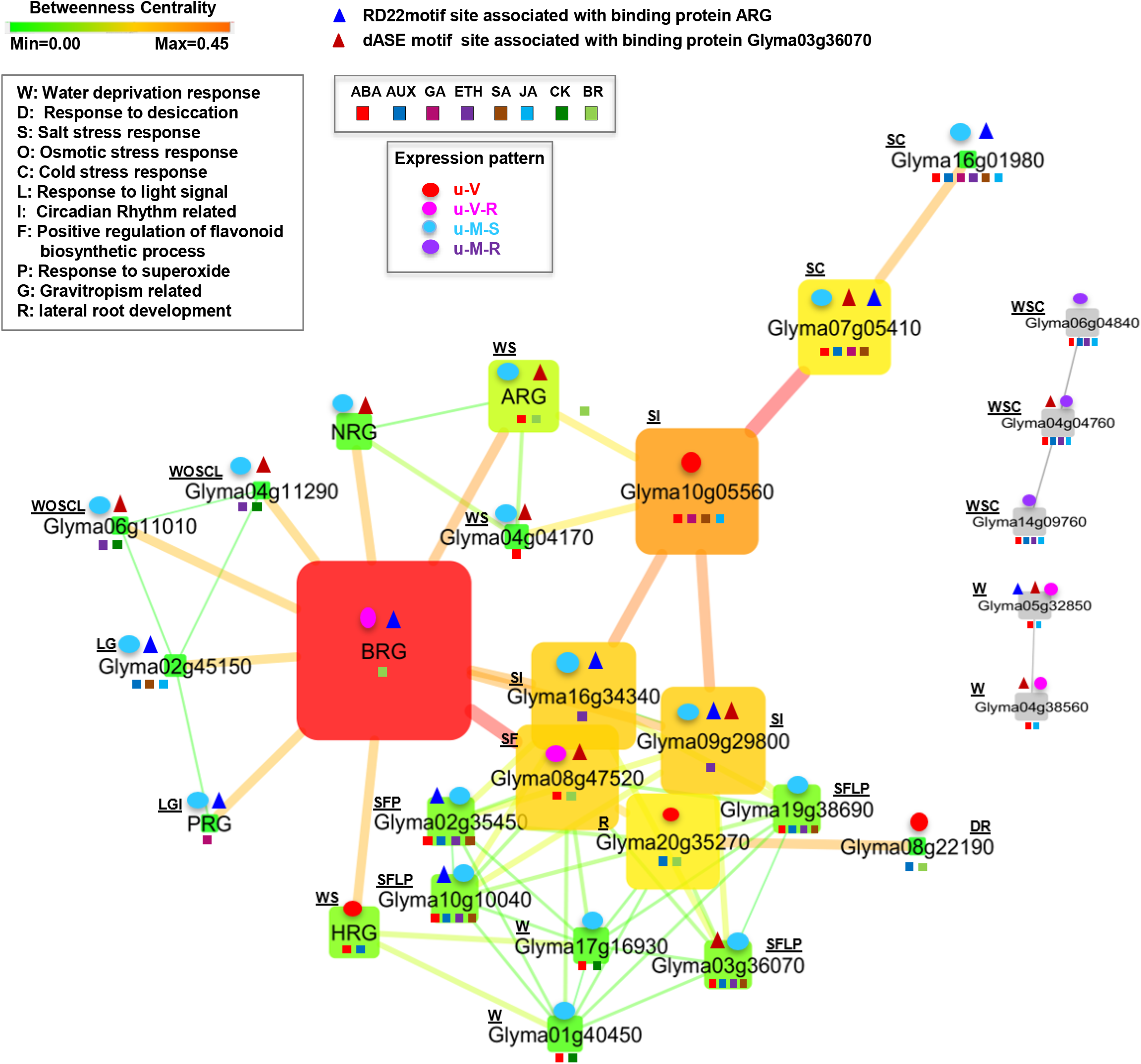
Core sensory TF network that functions in soybean primary roots under water-deficit conditions. Gene IDs are shown in boxes. The node size corresponds to the strength of the combined evidence from SoyNet and the Y1H assay. The color intensity indicates the betweenness of centrality in the network. Edge betweenness is indicated by line thickness. Different abiotic stress responses and the other pathways involved are represented by underlined capital letters associated with each core TF. Smallsquares of different colors connected with each Gene ID box represent different hormone responses. ABA, abscisic acid. AUX, auxin. GA, gibberellic acid. ETH, ethylene. SA, salicylic acid. JA,jasmonic acid. CK,cytokinin.BR, brassinosteroid. Triangles represent the motifs on the promoters of core TFs showing possible binding sites, including motifs RDmotif22 and dASE site. Ovals attached to each Gene ID represent the gene expression pattern under different water-deficit levels in soybean primary roots. A heatmap of gene expression levels is shown in Figure S3A.

### Core TFs help maintain root growth, and ARG is a repressor of *BRG*

To assess the acute ABA and osmotic stress responses of *BRG*, we generated hairy root lines harboring the *promBRG::GFP* reporter construct and measured reporter activity at 5 h and 48 h under 10 μM, 50 μM, 100 μM ABA and PEG (−1.7 MPa) treatment. The reporter gene responded to ABA, which was inversely correlated with ABA concentration and incubation time (Fig. 7a). In addition, we examined soybean root samples at 7 days after 10 μM, 50 μM and 100 μM ABA treatment. *BRG* levels significantly increased in response to 10 μM ABA, followed by a decrease compared with the untreated control, whereas *ARG* expression increased with increasing concentration of ABA, which is in agreement the possible role of ARG as a repressor of *BRG* expression (Fig. 7b). This hypothesis is also based on our finding that ARG bound to the *BRG* promoter in Y1H assays. We performed chromatin immunoprecipitation (ChIP) using transgenic hairy roots transformed with the *ARG* minigene (*35S*::*ARG*:*GFP*) after 3 days of dark treatment plus 100 μM ABA until the expression of *BRG* was repressed (Fig. 7c). We determined the enrichment of *BRG* genomic regions after immunoprecipitation with an anti-GFP antibody by qPCR using primers covering the *BRG* promoter, 5’UTR and CDS. Our results indicate that ARG associated with a region upstream of the transcription start site containing RDmotif-22 (CCACGTG) (Fig. 7d). Taken together, these results clearly demonstrate that ARG directly interacts with *BRG in vivo* with or without ABA application.

**Figure 7.**
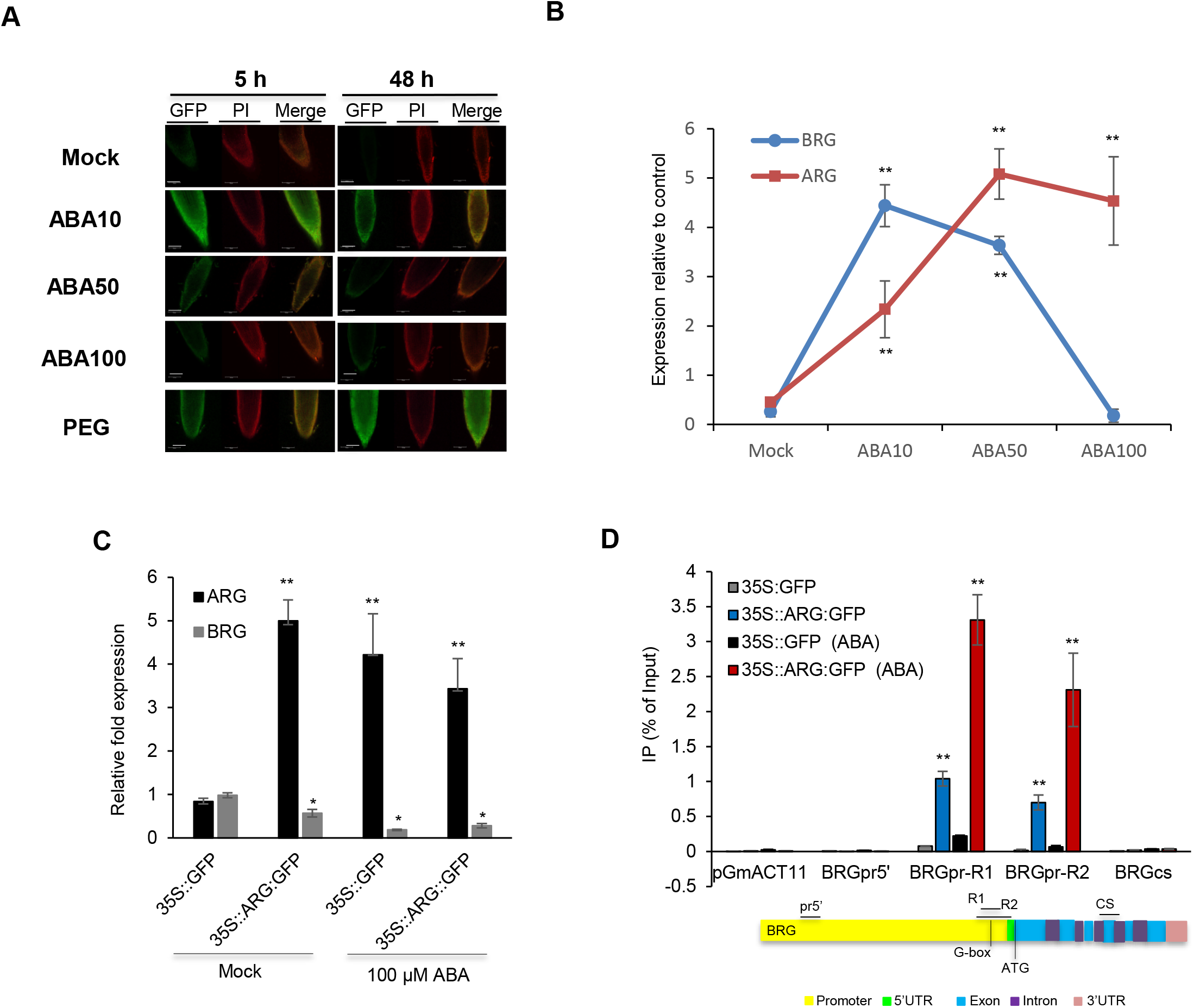
ARG directly binds to the *BRG* promoter and represses its expression. **(A)** promBRG::GFP expression after ABA or PEG treatment. Soybean transgenic hairy root tips (PROM-BRG::GFP) used for scanning were incubated on ½ MS solid medium (3% sucrose, 0.6% agar) containing 10 μM, 50 μM and 100 μM ABA or PEG-infused medium (−1.7 MPa) at 5 h and 48 h. (representative images are shown; n=4 per line, with three independent lines). Cell walls were stained with propidium iodide (PI, red). Scale bar: 100 μm. **(B)** The expression of *BRG* and *ARG* is inversely correlated with ABA levels. *ARG* and *BRG* expression levels in soybean roots relative to the *Glyma18g52780* (*GmACT11*) control were evaluated after 7 days on ½ MS solid medium supplemented with 10 μM, 50 μM and 100 μM ABA. (n=3 biological replicates with 3 technical replicates; mean ± SE). Significance relative to mock-treated root samples was determined by two-tailed Welch’s *t*-test: double asterisks denote significance level at P < 0.001. **(C)** *ARG* and *BRG* expression relative to the *GmACT11* control in transgenic 35S::ARG::GFP soybean roots relative to the empty vector control (35S::GFP). Relative *ARG* and *BRG* expression levels were evaluated after 3 days with or without 100 μM ABA treatment (n=3 biological replicates with 3 technical replicates. mean ± SE). Significance relative to the empty vector control in mock samples was determined by two-tailed Welch’s *t*-test: single and double asterisks denote significance level at P < 0.01 and P < 0.001, respectively. **(D)** ChIP-qRT-PCR of four areas of the *BRG* genomic regions. Values for 35S::ARG::GFP and 35S::GFP were calculated as the percentage of the input after 3 days with or without 100 μM ABA. The relative placement of the amplified regions is marked by black lines. (Regions: BRGpr5’ = −1967/-1882, BRGpr-R1 = −124/-65, BRGpr-R2 = − 120/−37, BRGcs = +1004/+1079, base pairs from the transcriptional start site). n=3 biological replicates with 3 technical replicates (mean ± SE). Significance relative to the control (promoter region of *GmACT11*; pGmACT11) was determined by two-tailed Welch’s *t*-test: double asterisks denote significance level at P < 0.001.

To investigate the possible roles of selected key TFs in regulating root growth under water deficit, we generated transgenic soybean hairy roots overexpressing *BRG*, *ARG*, *NRG, HRG* and *PRG,* as well as roots with loss-of-function due to RNA silencing (*arg-si* and *brg-si*) and roots expressing gene fused to the CRES repressor domain (*ARG*-*cres* and *BRG*-*cres*). We discovered that overexpressing *BRG* and its four candidate regulators, had effect on root growth maintenance under ABA (Fig. S5a, b and g), NAA (Fig. S5f and g) and osmotic stress treatment (Fig. S5 c and g). Interestingly not all of four regulators appear to activate *BRG* expression. Only overexpressing *HRG* modestly upregulated *BRG*, whereas *BRG* was downregulated in *ARG-*ox*, PRG-*ox and *NRG*-ox (Fig. S5j). *BRG* expression was reduced in *ARG-ox* and *ARG*-cres with or without ABA treatment, significantly increased in *arg-si* samples without ABA application, and slightly increased in the ARG deficient lines in response to ABA compared to the control (Fig. 8a). *ARG* expression in *brg-si* root lines was significantly higher than the control with or without ABA treatment, whereas *ARG* was downregulated in both the *BRG-ox* and *BRG-cres* lines, indicating that BRG also negatively regulates the expression of *ARG* (Fig. 8b).

**Figure 8.**
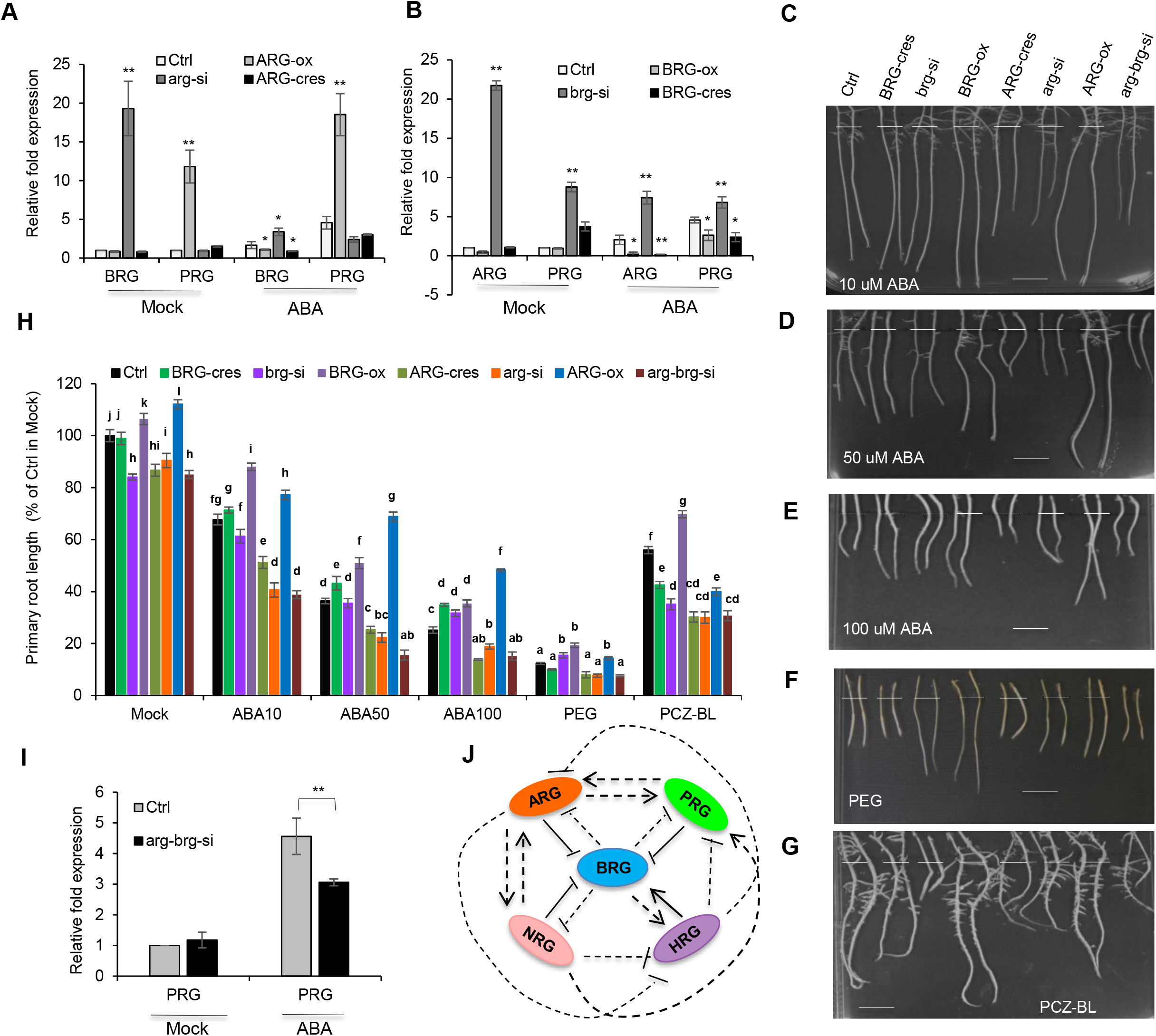
ARG and BRG antagonistically regulate *PRG*, and their effect on root growth maintenance. **(A)** RT-qPCR showing *BRG* and *PRG* transcript levels in root samples of the empty vector control (Ctrl), *ARG-ox, arg-si* and *ARG-cres m*utants treated for 3 days with 100 μM ABA relative to mock-treated roots. **(B)** RT-qPCR showing *ARG* and *PRG* transcript levels in root samples from Ctrl, *ARG-ox, arg-si* and *ARG-cres* plants treated for 3 days with 100 μM ABA relative to mock-treated roots. (A)–(B) Expression was normalized to the *GmACT11* reference gene (n=3 biological replicates with 3 technical replicates; mean ±SE). Significance relative to the empty vector control was determined by two-tailed Welch’s *t*-test: single and double asterisks denote significance level at P < 0.05 and P < 0.001, respectively. **(C) –(G)** Differential growth responses in primary roots of *BRG-ox*, *brg-si*, *BRG-cres*, *ARG-ox*, *arg-si*, *ARG-cres* and *arg-brg-si* after 7 days of treatment with 10 μM ABA, 50 μM ABA, 100 μM ABA and PEG (−1.7 Mpa). PCZ-BL treatment involved 4 days of 5 nMBL treatment after 3 days of 10 μM PCZ treatment. Scale bar: 1 cm. **(H)** Quantification of the primary root growth of *BRG-ox*, *brg-si*, *BRG-cres*, *ARG-ox*, *arg-si*, *ARG-cres*and *arg-brg-si* in response to 10 μM, 50 μM, 100 μM ABA, PEG (−1.7 Mpa) or PCZ-BL (10μM PCZ-5 nMBL). Each sample from different root tips representing independent transformation events (mean ±SE; n=15). Comparisons are made between empty vector control plants and mutants prepared using the same growth conditions and the same treatments. Bars with the same letter are not significantly different (two-way ANOVA + Tukey HSD, P < 0.05, ANOVA table in Table S12 showing a significant genotype: hormone/PEG treatment interaction term) in the response to PCZ-BL, ABA and PEG treatment. **(I)** RT-qPCR showing *PRG* transcript levels in root samples of the control (Ctrl) and the *arg-brg-si* mutant treated for 3 days with 100 μM ABA relative to mock-treated roots. Expression was normalized to the *GmACT11* reference gene (n=3 biological replicates with 3 technical replicates; mean ± SE). Significant differences relative to the empty vector control were determined using two-tailed Welch’s *t*-test; double asterisks denote a significance level at P < 0.001. **(J)** Model depicting the possible feedback and feed-forward loops among ARG, BRG, PRG, NRG and HRG. Black lines indicate transcriptional activation (arrows) or inhibition (bar ends). Solid lines represent direct interaction. Dotted lines represent unresolved or indirect relationships.

We discovered that *BRG-ox* roots were significantly longer than empty vector control and *ARG*-ox roots under 10 μM ABA and PEG (−1.7 MPa) treatment, whereas *ARG-ox* roots were much longer than the control under ABA 50 μM and 100 μM treatment (Fig. 8c, d, e, f and h). Interestingly, *BRG-cres* and *brg-si* primary roots did not lose the ability to maintain root growth, exhibiting some root growth under 10 μM, 50 μM and 100 μM ABA treatment (Fig. 8c, d, e, f and h). This might be due to the presence of the other homologs or the strength of ARG in the absence of the repressive activity of BRG, since *arg-brg-si* roots did not sustain much more growth than control roots when treated with various concentrations of ABA or PEG treatment. Following pretreatment with PCZ for three days, only *BRG-ox* roots were hypersensitive to BL (0.5 nm), whereas the other roots were not (Fig. 8g and h). Overall, these results suggest that BRG and ARG are not genetically redundant in the regulation of root elongation. Moreover, their transcriptional activity might depend on the severity of water stress, and their antagonistic interplay, which occurs in an ABA dependent manner, indicates that the actions of BRG and ARG are not equally required throughout exposure to water stress signals.

### ARG and BRG regulate a set of core TFs and have an ABA dose-dependent effect on dark-grown root growth

To investigate the roles of BRG and ARG in our core TF network, we analyzed the transcriptional activity of some selected core TFs when *ARG* or *BRG* was misregulated. *PRG* expression in the *BRG-ox*, *BRG-cres, ARG-cres*, *arg-si* and *arg-brg-si* lines was significantly downregulated in response to ABA and more drastically enhanced in *ARG-ox* and *brg-si* lines with or without 100 μM ABA treatment, indicating that BRG and ARG antagonistically regulate the expression of *PRG* (Fig. 8a, b and i). Therefore, we reasoned that ARG, BRG and PRG might jointly form a feed-forward loop to adjust the responses to ABA. We found that NRG and HRG are involved in these core TF interactions through direct binding to the *BRG* promoter and that they regulate the expression of *BRG*, *ARG* and *PRG*. Together, we proposed a model describing the mutual regulation among *ARG*, *BRG*, *PRG*, *NRG* and *HRG*, which integrate multiple hormone pathways with the phytochrome signaling pathway to sustain the growth of the underground root system (Fig. 8j). We also observed significant differences in expression among the other core TF genes in the network in *BRG-ox*, *BRG-cres, ARG-cres*, *arg-si* and *arg-brg-si* plants (Fig. S6). These observations are in agreement with the notion that some core TFs have a direct effect on the expression of the other core TFs, and they suggest that ARG and BRG together regulate these core TFs, which might coordinate water stress responses through the formation of extended cognitive TF networks.

To study the roles of ARG and BRG in plant growth/development and in response to drought conditions, we analyzed transgenic *BRG-ox* and *ARG-ox* Arabidopsis plants. The *BRG-ox* and *ARG-ox* lines had larger leaves, higher biomass, longer primary root and total root length and more root tips than wild-type (WT) Col-0 plants (Fig. S7 a and Fig. 9 a-d). Only the *BRG-ox* lines, however, had larger floral organs and siliques than comparable WT and *ARG-ox* plants (Fig. S7 b and c). In addition, ARG and NRG enhanced drought tolerance when overexpressed in Arabidopsis (Fig. 9 e). We performed a root growth response assay in which WT, transgenic *BRG-ox* and *ARG-ox* Arabidopsis plants were grown in long days for 3 days, followed by 7 days of growth on MS medium supplemented with 10 μM, 50 μM ABA or PEG (−1.7 MPa) under continuous light or in the dark. No significant difference was observed between WT, *ARG-ox* and *BRG-ox* plants under continuous light, but the roots of dark-grown *ARG-ox* and *BRG-ox* plants were more tolerant to ABA or PEG compared to dark-grown WT plants (Fig. 9 f–g). We also found that only *BEE1* expression was repressed in the dark, which corresponds to the presence of the ARG binding site RDmoitf22 locating in the promoter of *BEE1* (Fig. S7 d and e). Finally, to examine whether a functional link also exists between the expression of *BRG* and light regulation in soybean, we analyzed root growth in WT, *BRG*-ox, *BRG-CRES* and *brg-si* soybean plants treated with 10 μM, 50 μM and 100 μM ABA under continuous light and in the dark. Light-exposed *BRG-ox* roots were more sensitive to ABA than WT, whereas dark-grown *BRG-ox* roots showed less inhibition and continued growing in the presence of 10 μM and 50 μM ABA (Fig. 9 h and i). Taken together, these results suggest that regulons are conserved between *ARG* and *BRG*, which have ABA dependent effects on dark-grown roots, indicating that hormonal and light signaling pathways utilize the same junctions to help plants adapt to water stress.

**Figure 9.**
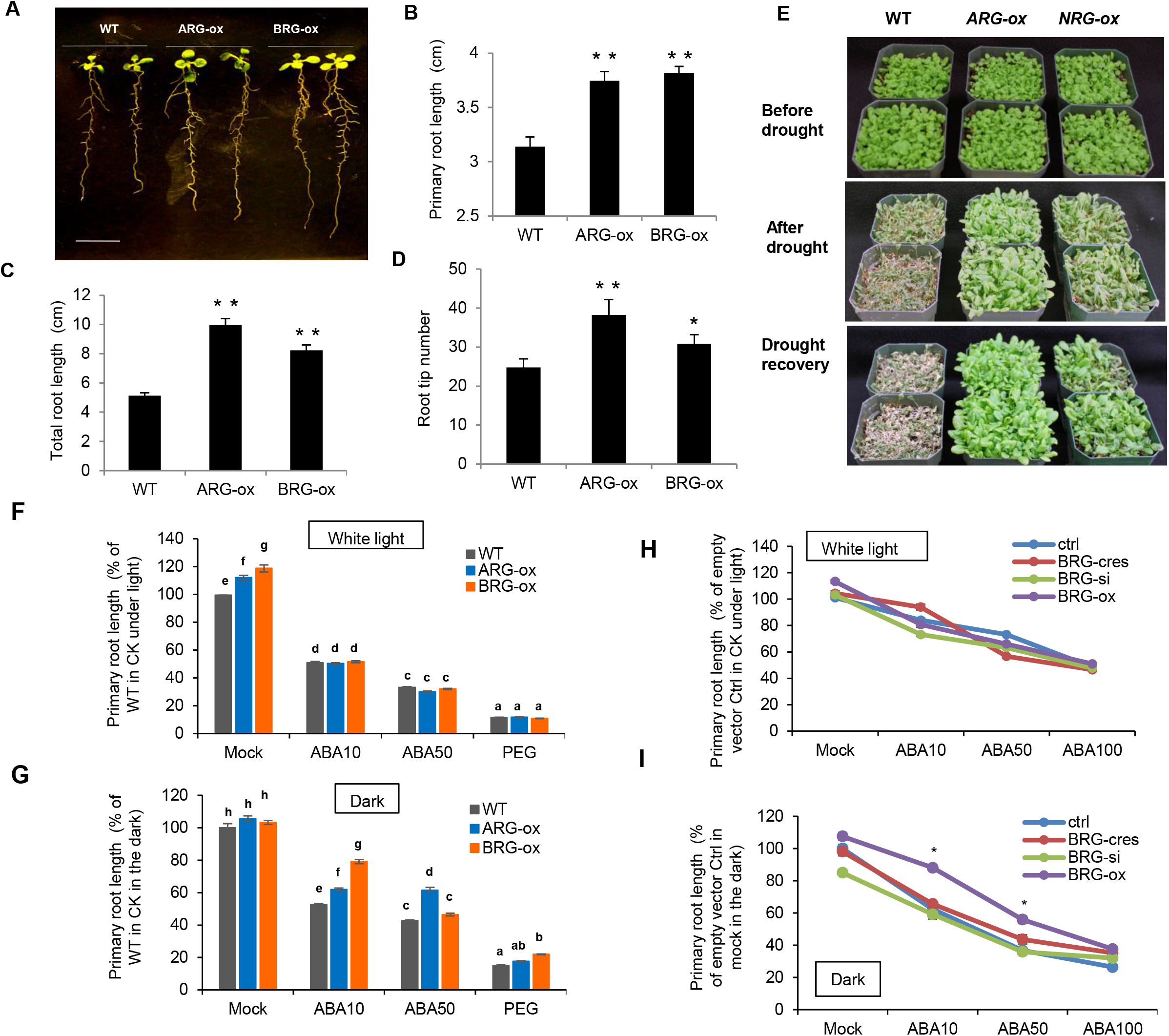
BRG and ARG have an ABA dose-dependent effect on dark-grown root growth. **(A)** Increased primary root length and root tip number of *BRG-ox* and *ARG-ox* Arabidopsis plants. Scale bar: 1 cm. **(B)-(D)** Primary root length, total root length and number of root tips in *BRG-ox* and *ARG-ox* plants compared with WT (Col-0). The germinated seedlings were transferred to medium in square Petri dishes and grown vertically in long days for 7 days. Primary root length, total root length and root tip number were measured (mean ± SE; n=18-24). Significance was determined by two-tailed Welch’s *t*-test: single and double asterisks denote significance level at P < 0.01 and P < 0.001, respectively. (E) Transgenic Arabidopsis *ARG-ox* and *NRG-ox* plants are drought resistant. Fourteen-day-old T3 transgenic Arabidopsis plants were well-watered for 12 days, after which water was withheld for 12 days, followed by re-watering for 5 days. **(F)-(G)** Quantification of Arabidopsis WT, *ARG-ox* and *BRG-ox* primary root length in response to 10 μM ABA, 50 μM ABA or PEG (−1.7 Mpa) under continuous light and in the dark. The comparisons are between WT plants and mutant plants under the same growth conditions and the same treatment (mean ± SE; n=18). Bars with the same letter are not significantly different (two-way ANOVA + Tukey HSD, P < 0.01, ANOVA table in Table S13 showing a significant genotype: ABA/PEG treatment interaction term) in the response to ABA and PEG treatment. **(H)-(I)** Response of *BRG-cres*, *brg-si* and *BRG-ox* transgenic soybean roots to 10 μM ABA, 50 μM ABA and 100 μM ABA after 7 days of culture under continuous light or in the dark. Two independent experiments were performed, each sample from different primary roots from independent transformation events (mean ± SE; n=12). Significance was determined by two-tailed Welch’s *t*-test: single asterisks denote significance level at P < 0.01.

## DISCUSSION

In this study, we developed an effective platform for identifying interconnected core TFs and prioritized key regulators involved in root growth maintenance under water deficit in soybean. We used four steps to identify and prioritize core TFs to help elucidate the regulation of root growth under water deficit: 1) target a set of genes involved in root elongation in the water stress response and stress adaptation across different root regions; 2) identify a group of conserved functional motifs related to root growth regulation in response to water stress; 3) decompose genome-wide conserved motif-associated genes to trace back interconnected core TFs ingroup that respond to both endogenous and abiotic stimuli; and 4) combine co-functional TF–TF networks with TF– DNA networks to prioritize key regulators. Using this platform, we succeeded in identifying a set of core TFs that are potentially important regulators of soybean root growth under water deficit conditions. Previous studies in Arabidopsis have shown that a number of homologs of these core TFs, e.g., ABF2, RVE7/8, RAP2.4, IAA14/IAA19, BLH1, PIF3/PIF4, HAT22, HAT2, NF-YB3, ZAT10 and VNI2, are involved in abiotic stress responses through mediating crosstalk among auxin, GA, BR, ABA, ethylene and phytochrome signaling. Therefore, we constructed a conserved core sensory TF network to explore their complex interconnected regulatory circuits by examining mutual cross-regulation among this group of TFs, which process signals to generate appropriate responses to water stress. Based on current and previous findings, we developed a model describing how root growth might be regulated under water deficit (Fig. 10).

**Figure 10.**
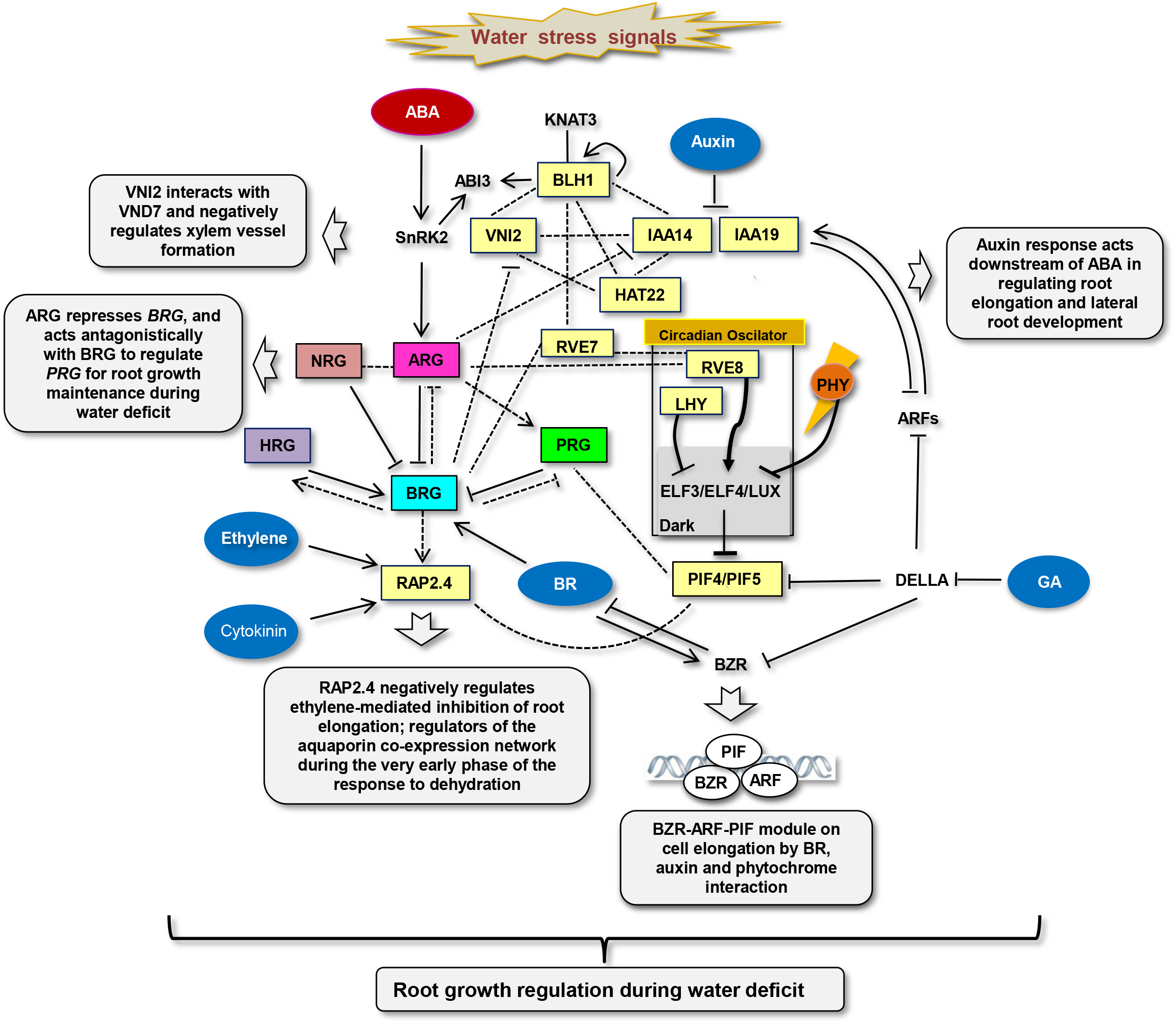
Core sensory TF network integrating hormonal and environmental signaling in roots under water deficit conditions. Core TFs mutually regulate each other, acting as convergence points in the crosstalk among ABA, auxin, GA, BR, ethylene, cytokinin and light signaling to regulate root growth in response to varying water deficit conditions. The genes in boxes are derived from our core sensory network. The genes in yellow boxes and listed outside the boxes are putative orthologs of Arabidopsis genes that are thought to function in the multi-hormonal interaction involved in root growth regulation or the cell elongation module. Hormones are shown in ovals. Black lines indicate known interactions based on published data and direct interactions confirmed by experimental evidence in this study. Black dotted lines represent unresolved relations or indirect interactions awaiting additional investigation. Lines represent transcriptional activation (arrows) or inhibition (bar ends). Functional descriptions are indicated in grey boxes.

The hormonal signaling mechanism that controls root growth in response to water availability is not yet fully elucidated. However, we know that ABA plays an important role in regulating stress-responsive genes, mainly through three bZIP TFs: AREB1/ABF2, AREB2/ABF4 and ABF3 (43). In our network, *ARG* and another *ABF2*-like gene (*Glyma04g04170*) were upregulated only in root tissue under water deficit, whereas other ABF genes including *GmbZIP1* (44) were upregulated in both root and shoot tissue (45). There are four *BLH1* homologs in our core TF network. It has been reported that BLH1 interacted with another homeobox protein, KNAT3, and synergistically increased the ABA responses by binding to the promoter of *ABI3*, which plays a role in auxin-induced lateral root formation (46). BLH1 also promotes its own expression, which is expected to accelerate this feedback loop to reinforce the ABA signaling network (42). So we suggest that *Glyma03g36070*, one of the BLH1-like genes in the network, and its binding site dASE motif may participate in this regulation. Our conserved motif-associated network analysis method can be used to associate specific groups of duplicate genes with an adaptive trait, which will help elucidate how the CREs in duplicate genes differ and why duplicate genes have different responses to stressful environments.

The auxin response acts downstream of ABA in regulating root elongation, which is primarily dependent on altered Aux/IAA gene expression or altered gene expression in combination with additional factors (47). We also discovered the homologs of *IAA14*/IAA19 in our network. GA promotes cell elongation largely by releasing the DELLA-mediated repression of PIF4, BZR1 and ARF6 (48). In particular, the existence of the BZR-ARF-PIF module explains how auxin, BR, GA, light and temperature coordinately regulate cell elongation in hypocotyls and likely other parts of the shoot in Arabidopsis (48). Based on our network, the BR-responsive gene *BRG* appears to function as a signaling intermediary in multiple pathways, likely through interactions with homologous genes of *ABF2*, *VNI2*, *RAP2.4*, *RVE7*, *PIF4/PIF5* and *PIF3*. PIF3 regulates hypocotyl length downstream of an auxin and ethylene cascade, whereas PIF4 and PIF5 regulate hypocotyl length upstream of this cascade (49). It remains to be elucidated whether the mechanisms underlying signaling integration involve a similar module in root tissue and how water stress signaling is integrated into this module.

Our present data suggest that BRG, ARG and PRG may achieve their roles in maintaining root growth through mediating crosstalk among BR, ABA and light signaling. ARG may play a major role in maintaining root growth in an ABA-dependent manner that leads to the downstream regulation of AUX/IAA factors to support auxin-related coordination between root elongation and lateral root development. BRG, on the other hand, may coordinate ABA-dependent and ABA-independent pathways through the regulation of two *RAP2.4*-like genes (*Glymao6g11010* and *Glyma04g11290*), which might negatively regulate ethylene-mediated inhibition of root elongation (50). Our data suggest that ARG activates *PRG* by repressing *BRG*, whereas PRG and BRG negatively regulate each other in soybean root tissue. We discovered that a similar mechanism might function in soybean and Arabidopsis in which overexpressing *ARG* represses *BEE1* through binding to a conserved G-box motif in its promoter.

One notable regulatory mechanism is that HY5 and PIFs co-regulate photosynthetic genes at common G-box motifs, in conjunction with the circadian oscillator, to adjust the levels of rhythmic photosynthetic gene expression (51). In our network, ARG and PRG might act at a common G-box-type motif (RDmotif22) to regulate *BRG* expression in response to water stress. Perhaps this dual control system works in conjunction with several circadian oscillator factors described in our network, including two *LHY*-like genes, two *RVE7*-like genes and one *RVE8*-like gene. CCA1 and LHY profoundly influence hypocotyl elongation and flowering time by regulating the expression of *PIF4* and *PIF5*, respectively (52, 53). *RVE7* acts primarily as a clock output gene and positively regulates the expression of an auxin biosynthetic gene (54). *RVE8* affects light inhibition of hypocotyl elongation at low light intensities (55). The plant clock is organ specific, which could be attributed to the different light inputs to the shoot versus root clocks (56). The finding that ARG and BRG help maintain dark-grown root growth in response to ABA and osmotic stress suggests that light has an additive, negative effect on root growth in response to these stresses, which is similar to the finding that the illumination of roots alters salt and mannitol stress responses (57). We therefore suggest that clock genes operate in conjunction with ARG, BRG, PRG and other core TFs to adjust the levels of root growth in response to varying water deficit conditions in the dark compared to their effect on the inhibition of shoot growth in the light.

One important feature of our platform is the identification of functionally conserved cis-regulatory motifs and the traceback of these motif-associated core TFs as mediators of signaling in response to both endogenous and abiotic stress stimuli. We suggest that water stress-responsive gene expression in soybean roots is controlled by a core TF network consisting of mutually regulated TFs that bind to the cognate motif sites in their own genes and regulate their own expression and/or bind to the motif sites in many other core TF genes and regulate their expression.

Although both ARG and BRG help maintain root growth under water deficit conditions, which core TF dominates this process might depend on the degree of water stress and/or ABA dose. BRG might play a more important role in rapid responses to water stress during early stages of the water stress response by enhancing BR pathways and regulating ethylene production. During long-term water stress, ARG might terminate this signaling under normal and very mild stress conditions by repressing *BRG* and the other factors involved, making a more “permanent decision” to switch to a new cellular state or fate in response to more severe water stress signals. Future studies should focus on determining the molecular mode of action of the core TF complexes in the control of root growth and explore how these core TFs form complex feedback and feed-forward loops. These loops might act as stress signal-sensitive delays or accelerators to regulate root growth by efficiently coordinating cellular responses to very mild, mild and severe drought stress. The strategies used in this study can be adopted to identify key genes and their interconnected networks for other complex biological processes in plants or other species. Our core sensory TF network lays the foundation for understanding the drought response in crops over long periods and should provide guidance for further investigating the cell-type-specific transcriptional regulatory circuitry that maintain root elongation but inhibit shoot growth under drought. Such studies might ultimately lead to the fine-tuning of cell-type-specific GRNs to improve crop tolerance and to avoid the negative effects of drought on plant growth and yield.

## METHODS

### Plant materials and growth conditions

Soybean (*Glycine max* cv. Magellan) seeds were surface sterilized in 1% NaClO solution for 2 min, rinsed in deionized water for 30 min and germinated between sheets of germination paper moistened with a solution of 5 mm CaCl2 and 5 mm Ca(NO3)2 for 36 h at 29°C and near-saturation humidity in the dark. Seedlings with primary roots approximately 15 mm long were transplanted against the interior surfaces of Plexiglas cylinders (14.5 cm diameter) filled with a 1:1 (v/v) mixture of vermiculite (no. 2A, Therm-ORock East Inc., New Eagle, PA, USA) and Turface (Profile Products LLC, Buffalo Grove, IL, USA) at water potentials of −0.1 MPa (well-watered treatment, moistened to the drip point) or −1.6 MPa (water-stressed treatment) according to Yamaguchi et al. (2010). Three regions of soybean primary roots were identified based on their growth under water stress treatment: region 1 (R1: 0–4 mm from the apex, including the root cap) exhibits the same elongation rates under well-watered and water stress conditions; region 2 (R2: 4–8 mm) exhibits maximum elongation rates in well-watered soil, but progressive deceleration in growth under water stress; and region 3 (R3: 8–15 mm) exhibits slow growth under well-watered conditions and no growth under water stress. At 48h time point (root tip water potential had decreased to approximately −1.6 MPa by this time), R3 samples were collected under well-watered conditions and there were no R3 samples under water stress conditions because root elongation rates had been completely inhibited in R3. The root tips were sectioned into regions 1–2 (for both water stressed and well-watered controls) at 5h, while 1–2 (water stressed) and 1-3 (well-watered controls) at 48h, giving a total of nine samples per experiment. In each of three replicate experiments, approximately 100 roots were harvested from the water-stressed treatment and from the well-watered controls. The root samples were immediately frozen in liquid nitrogen and stored at −80°C.

### RNA extraction, labeling and array hybridization

Total RNA was isolated from root tissues using TRIZOL reagent (Invitrogen, Carlsbad, CA, USA). Genomic DNA was eliminated by digesting the RNA with Turbo DNA-free DNaseI (Ambion) according to the manufacturer’s instructions. After DNaseI treatment, the RNA concentration was determined and the quality checked using an Agilent 2100 Bioanalyzer (Agilent Technologies, CA). Total RNA was used for Affymetrix Soybean Genome GeneChip hybridization and analysis following the manufacturer’s recommended protocols (Affymetrix, CA). cDNA synthesis, cRNA amplification and the synthesis of sense-strand cDNAs were carried out using an Ambion WT Expression kit. Hybridizations were conducted at the DNA Core Facility, University of Missouri (http://biotech.rnet.missouri.edu/dnacore) following the standard Affymetrix procedures (Affymetrix, Santa Clara, CA). The microarrays were scanned with a high resolution GeneChip scanner. Scans from each microarray were analyzed with Affymetrix GeneChip Microarray Suite version 5.0 software (MAS 5.0) in Expression Console version 1.1 of the Affymetrix GeneChip Command Console suite. The microarray data sets used in the present study were deposited in the Gene Expression Omnibus under accession number GSE102749 (http://www.ncbi.nlm.nih.gov/geo/query/acc.cgi?acc=GSE102749).

### Microarray experiment and data analysis

The data files containing the probe set intensities (.cel files) were used for background correction and normalization by the log2 scale GC-RMA procedure provided in GeneSpring Suite 11. Twoway ANOVA was used to identify genes that were differentially expressed at the 5 h and 48 h time points of five pairwise comparisons between WS and WW root regions R1, R2 and R3 with a False Discovery Rate (FDR) of 0.05. Benjamini and Hochberg multiple testing corrections were used in the Two-way ANOVA analysis. The differential expression levels of water-deficit-regulated genes were clustered using DNA-Chip Analyzer (dChip) (Li and Wong, 2001). The functional classification of differentially regulated genes was analyzed using MapMan (58). Enrichment of each category was tested with Fisher’s exact test (p-values ≤ 0.05).

### Motif searching using MEME

The 3-kb sequences upstream of the gene transcription start sites for all annotated soybean genes (Phytozome v9.1) were collected from the same strand as the gene (in the same orientation). The putative promoter regions for each of the clusters were written to individual files. Additionally, one file contained 279 randomly selected promoters, which were written to a separate file for use as a representative promoter-negative set. Discriminative Motif Discovery (http://meme-suite.org/tools/meme) was then performed using MEME V4.11. Position-Specific Priors (PSPs) were derived for each cluster using clusters I, II and III as the positive set and the 279 randomly selected promoters as the negative set, with psp-gen. Using this discriminative method, all motifs identified by MEME should be enriched in the clusters. MEME was then run using the -psp option, -mod zoops (zero or one motif per sequence), specifying the maximum width of the motif as 15 bp and the minimum width as 6 bp in both strands of sequences. In addition, motif sequences were used to search each soybean promoter sequence. Counts of occurrence for the forward and reverse complement of each motif were noted. Motif enrichment and significance levels were determined using 1000 random gene sets, which were constructed by sampling without replacement using all soybean genes. Thousand random samples of 279 genes were used to determine the mean occurrence of each motif in the promoters of all genes. Observed counts for each motif found in the 279-gene set were then compared to the mean. The probability that a motif is enriched was expressed as a p-value, and the p-value cutoff was 0.05.

### Motif conservation analysis

Motif phylogenetic conservation analysis was performed as previously described (25), with *Glycine max* (Phytozome 9.1) as the query species and *Arabidopsis thaliana* (TAIR10), *Carica papaya* (Hawaii Agriculture Research Center), *Theobroma cacao* (Phytozome 9.1), *Populus trichocarpa* (Phytozome 9.1), *Prunus persica* (Phytozome 9.1), *Cucumis melo* (Melonomics v3.5), *Eucalyptus glandiformis* (Phytozome 9.1), *Vitis vinifera* (GeneScope v1), *Solanum lycopersicum* (ITAG 2.3), *Beta vulgaris* (RefBeet 1.1), *Oryza sativa* (MSU RGAP 7) and *Amborella trichopoda* (Amborella V1.0) as the comparator species from the PLAZA 3.0 database. For the alignment-based approach, only the Sigma alignment tool (59) was used. The newly discovered motifs were used as inputs for the comparative motif mapping approach and overlapped with the alignment-based conserved non-coding sequences obtained for *G. max*. In the alignment-based approach, pairwise alignments are generated between the query promoter (soybean putative promoter region: 2 kb sequences upstream of gene transcription start) and the orthologous promoters. The algorithm checks the significance of pairwise regions against a background model of 1000 randomly generated orthologous groups for which the same process of alignment between the promoters is performed. A score is calculated based on both the length of the region and the number of species. The p-value is calculated based on how many times the algorithm observes conservation scores in the random groups that are as good as or better that the real conservation score from the original orthologous group. All conserved motifs had a false discovery rate <10%.

### Motif similarity tree

The newly discovered motifs were compared to motifs from the PLACE database using the STAMP tool (26). Motif similarity was determined with the Pearson correlation coefficient and aligned with the Ungapped Smith-Waterman algorithm, and a UPGMA tree was generated. Tree visualization was performed using the FigTree tool (http://tree.bio.ed.ac.uk/software/figtree/).

### GO enrichment analysis of the conserved targets of newly discovered motifs

GO annotations for *G. max* were obtained from the PLAZA 3.0 database (60). Of the 7954 target genes obtained from the multispecies phylogenetic footprinting analysis, each gene had at least one of the 49 previously identified conserved motifs. For each motif, the enrichment of conserved motif target genes toward GO slim annotations (hypergeometric distribution + Bonferroni correction) was determined. A heatmap was generated using Genesis (61), with a color gradient that shows the P-values of the different enriched gene sets.

### Phylogenetic analysis

To investigate their phylogenetic relationships, 20 BEE genes from 14 species were analyzed. Protein sequences were analyzed by the neighbor joining (NJ) method with the genetic distance calculated by MEGA v6.06 (62) (www.megasoftware.net/).

### Co-functional TF network construction

The functional network of selected TFs is based on a genome-scale co-functional network of soybean genes constructed using a machine learning approach with a Bayesian statistics framework (33). To train various data that support the functional association between soybean genes, a set of gold-standard gene pairs was generated by pairing two genes that share annotations using the KEGG, MapMan or Soycyc pathway database. Various biological data, including the co-citation of genes in research articles, co-expression, gene neighborhood, phylogenetic profiling and associalogs transferred from networks of the other species were also employed. The SoyNet (www.inetbio.org/soynet)-based method “find new members of a pathway” was used to explore functional connections among core TFs. Thirty-six TFs, including 33 rdcTFs and three core TFs candidates from 279 gene sets, were used as input genes. The SoyNet server was used to measure the retrieval rates of the 36 submitted TFs based on SoyNet connections among these TFs. The overall performance of the network for the retrieval of all core TFs was assessed based on receiver operating characteristic analysis (63), which can be summarized as the area under the ROC curve (64) score, indicating that core TFs for root growth regulation under water deficit are well connected to each other based on SoyNet analysis. Twenty-four core TFs were ultimately found to connect to each other in a co-functional TF–TF network, with the sum of LLS scores used to rank their interactions. Network statistics analysis was applied in Cytoscape (34), and the TFs were ranked based on their BC to current hub genes. The BC of a node reflects the amount of control that this node exerts over the interactions of other nodes in the network (Yoon et al. 2006), which helps target hub TFs that join communities (dense subnetworks) rather than TFs that lie inside a community. To integrate co-functional connections with TF-DNA interactions from Y1H, four candidate regulators (*ARG, PRG, NRG and HRG*) of BRG were added in 36 core TFs list and were explored the co-functional connections of these genes in SoyNet. Then TF-DNA interactions were combined with all co-functional connections to create a total interactions list of core TFs for statistically analysis using Cytoscape to visualize possible mutual regulation among core TFs involved in multiple signaling pathways.

### Construction of a Gateway-compatible library of soybean root-related water stress-responsive TFs

A representative TF library was constructed that included 214 root-related water stress-responsive TFs in 22 TF families. These drought-related TFs were selected based on a >2-fold change at least in one of five comparisons from soybean root elongation microarray data in response to water deficit. Soybean TF coding sequences were amplified by PCR and cloned into the pDONR221 Gateway donor vector (Life Technologies). All clones included in the collection (214 TFs) were sequence validated (Table S8). Each TF was transferred into a Y1H-compatible destination plasmid (pDEST22) carrying the GAL4-Activation Domain (GAL4_AD) located at the TF insertion site using One Shot™ Mach1™ T1 Phage-Resistant Chemically Competent *E. coli* (Invitrogen). TFs cloned in pDEST22 were thus expressed in yeast as C-terminal fusions to GAL4_AD. After destination plasmid validation by gene-specific colony PCR and sequencing, 214 clones were successfully cloned into pDEST22. The pDEST22-TF collection was arrayed in three 96-well plates.

### Bait construction and yeast one-hybrid assay

A total of 214 *E. coli* strains harboring different soybean TFs were arrayed in three 96-well plates, and plasmids were prepared using the Zymore plasmid purification DNA system according to the manufacturer’s recommendations. The oligonucleotides used to amplify motif repeats and promoter fragments (~ 500 bp) for mGbox, Gbox-I-gc, dASE, mdASE, PROM-BRG, PROM-*Glyma05g26930*, PROM-*Glyma04g04310*, PROM-*Glyma06g03450*, PROM-*Glyma02g15780* and PROM-*Glyma08g29100* are described in Supplemental Table S14. These bait DNAs were subcloned into pENTR/TOPO-D (Gateway; Invitrogen), followed by pGLAC recombined with the reporter gene *β-galactosidase* (Gal). The pGLAC recombinants were transformed into yeast strain Y4271 and screened against the root drought stress-related TF library using the Y1H protocol, as previously described (65). The DNA bait strains were tested for self-activation prior to screening by transforming them with prey vectors. pDEST22 empty vector was used as a negative control. The transformants were grown on SD -Trp plates (Clontech, 2% agar) for selection. Positive interactions were visually identified by the yellow color of the yeast cells caused by the cleavage of ortho-nitrophenyl from colorless ortho-nitrophenyl-β-D-galactoside by β-galactosidase. Each TF was tested in quadruplicate. The first screen was applied against the whole TF library with replicates. All interacting TFs were assembled into a small interaction library for the second screen. The second screen was applied with replicates in the same plate. Each positive TF clone was sequenced to reconfirm its identity.

### Dual-Luc assay-based quantification of the stress inducibility of conserved motifs/promoters

A vector system was created to generate a single vector with the CaMV 35S mini promoter (*35S*) fused to the firefly luciferase reporter gene and *35S* fused to the *Renilla* luciferase reporter gene. The constitutively expressed *Renilla* gene served as a control to normalize for transformation efficiency. This system included one destination vector, pLL-QMP, and one entry vector, pLL-P/M, with Gateway Technology (Invitrogen) used to clone the promoter fragment/motif repeats (Figure S3). The 3 kb pRTL2-*Renilla Hind*III digested fragment was inserted into *Sac*I-digested pFLASH to create pFLASH-FR with both firefly *LUC* and *Renilla* luciferase (*REN*) genes. *Hind*III-digested pFLASH-FR was ligated with the *Hind*III-digested PCR fragment *Mini35S* to yield pLL-QMP (Figure S4). Target motif repeats/promoter fragments were amplified from *G. max* genomic DNA using the appropriate primers with attB1 and attB2 sites. Each amplified fragment was cloned into a pDONR-P1-P2 vector by performing BP reactions to produce pLL-P/M. Finally, the fully functional expression vector pLL-QMP-PM was generated by Gateway LR cloning (Figure S4).

Transgenic soybean hairy roots were then generated (detailed below) using each final pLL-QMP-PM construct. *Agrobacterium rhizogenes* strain K599 was transformed with the pLL-QMP-PM construct by electroporation. The bacterial solution was drawn into a syringe and injected three times into the cotyledon node and the upper hypocotyls. Infected seedlings were transferred into pots. The plants were maintained in a growth chamber under a 12 h light/12 h dark cycle at 28°C for 2–3 weeks, at which time the hairy roots were approximately 5–10 cm long. Hairy root samples were harvested 6 h after treatment with 100 μM ABA, 150 mM NaCl2 or 16% PEG and assayed for luciferase activity using the Dual-Luciferase Reporter Assay System (Promega) according to the manufacturer’s instructions. Approximately 100 mg of tissue was frozen in liquid N and homogenized using a Retsch Mixer Mill MM400 for 1 min at 30 Hz. The ground tissue was thawed in lysis buffer (0.1 M HEPES, pH 7.8, 1% Triton X-100, 1 mM CaCl2 and 1 mM MgCl2) at 25°C for 15 min. Luciferase Assay Reagent II (50 μL) was added to 10 μL aliquots of the lysates to measure firefly luciferase activity using a Spectra Max M5/M5e plate reader to measure total light emission after a 1000 ms integration time. Firefly luciferase activity was quenched with 50 μL of Stop & Glo Reagent, which contains *Renilla* luciferin substrate, and total light emission was measured at a 100 ms integration time. An expression vector containing part of the coding sequence of the β-glucuronidase reporter gene rather than a TF gene was used to measure baseline firefly luciferase activity. To estimate the relative affinity of the TFs for each promoter fragment, five biological replicates of TF-expressing vectors were compared to the average results for the GUS expression vector. Firefly luciferase activity was divided by *Renilla* luciferase activity to normalize the transformation efficiency of each hairy root sample. The relative binding of the TFs to promoter bait sequences was determined relative to the control using two-way ANOVA + Tukey HSD.

### Generation of transgenic soybean hairy roots

Soybean line Williams 82 was used in this study. Sterile soybean seeds were germinated and grown in sterilized vermiculite for 7 days in a growth chamber at 25°C under fluorescent white light with a 16/8 h light/dark cycle. Transgenic soybean hairy roots were generated as previously described (66) with some modifications. *Agrobacterium rhizogenes* strain K599 was transformed with each individual construct by electroporation. Cotyledons from geminated soybean seedlings were inoculated with the bacterial culture to induce hairy roots, as previously described (66).

Fourteen days post-inoculation, transgenic hairy roots were identified based on growth on hygromycin. The roots were isolated and removed from the cotyledons 18–25 days postinoculation and cultured on ½ MS solid medium (3% sucrose, 0.6% agar) plus carbenicillin (500 mg/L) in the dark at 26°C. After 2–3 weeks, 3–4 cm long lateral roots or the primary root tip were cut and subcultured on ½ MS solid medium plus carbenicillin (500 mg/L). After subculturing 3–4 times on ½ MS solid medium containing carbenicillin, transgenic roots were used immediately, frozen or maintained on ½ MS solid medium (without carbenicillin) for treatment. The concentrations of the treatments were as follows: (10, 50, 100) μM ABA, 50 μM naphthaleneacetic acid (NAA), (5, 0.5, 0.05) μM BL and 10 μM PCZ. Stocks were made in 80% ethanol at identical dilutions except for NAA, for which the stock was dissolved in sodium hydroxide, with mock treatment performed using an equal amount of solvent alone. For the PEG, ABA, BL, PCZ and NAA treatments, transgenic hairy roots were incubated in the dark on PEG-infused plates (−1.7 MPa) (67, 68) or ½ MS solid medium containing 100 μM ABA, 50 μM naphthaleneacetic acid (NAA), (5, 0.5, 0.05) nM BL or 10 μM PCZ and measured 7 days later (n=12-15, three repeats). Statistical analysis for this study was carried out with R (version 3.2.4). For each analysis of effects of different genotypes on hormone/osmotic stress response, two-way ANOVA followed by Tukey HSD multiple comparison procedure were used to detect significant differences between mock control and dose of hormone/osmotic stress treatments associated with genotypes and primary root length. The linear model is y= intercept+ hormone/osmotic stress + genotype + interaction between stress and genotype +residual error. We first performed two-way ANOVA which not only assess the independent impact of the two factors but also can assess the interaction of the two factors. When the interaction term was found to be significant, we concluded that genotype affected hormone/osmotic stress response. *P*-value associated with particular levels of hormone/osmotic stress treatment were adjusted for multiple comparisons using Turkey HSD method.

### Quantitative RT-PCR analysis

Total RNA was extracted from root samples collected in the morning using an RNeasy kit (Qiagen). Genomic DNA was removed using an RNase-free DNase set (Cat# 79254, Qiagen). The cDNA was synthesized via treatment with reverse transcriptase and oligo(dT) primer (SuperScript III First-Strand Synthesis System; Invitrogen). Quantitative RT-PCR was performed on an ABI7900HT detection system (Life Technologies) using Maxima SYBR Green/ROX qPCR Master Mix (2X) (Cat# K0223, Thermo, USA) following the manufacturer’s protocol. The primer sets used for qRT-PCR are listed in Table S14. The comparative Ct method for quantification was used to quantify the relative expression levels of specific genes (69). Gene expression was measured in at least three biological replicates with three technical replicates. Significance relative to control was determined by two-tailed Welch’s *t*-test.

### Molecular cloning and constructs

The 2 kb *BRG* promoter of was subcloned in pENTR/TOPO-D and fused to *GFP* in the Gateway-compatible destination vector pMDC107 for soybean hairy root transformation. The *ARG*, *BRG*, *HRG*, *NRG* and *PRG* overexpression constructs were generated by subcloning in pDONR221, followed by cloning in pMDC32. For the ChIP construct, *ARG* was subcloned in pDONR221 and then fused to *GFP* in the pMDC83 vector. The *Arg-si* and *Brg-si* vectors were generated by subcloning synthetic fragments (Table S14) in pENTR/TOPO-D, followed by cloning in the destination vector pANDA (RNAi construct). The *ARG-cres* and *BRG-cres* vectors were subcloned in pDONR221 and fused to the CRES domain in pB7WG2-CRES. The primers used are listed in Table S14.

### Confocal laser-scanning microscopy

Confocal laser-scanning microscopy was carried out on a Leica TCP SP8. Excitation was performed using a 488-nm laser for GFP. Cell walls were stained with propidium iodide as previously described (69). Transgenic hairy root tips (PROM-BRG::GFP) used for scanning were placed onto PEG-infused plates (−1.7 MPa) (67, 68) or MS solid medium containing 10 μM, 50 μM or 100 μM ABA at 5 h or 48 h.

### Chromatin immunoprecipitation of ARG

Chromatin immunoprecipitation was conducted as previously described (65) Approximately 5 g (fresh weight) of root tissue (35S::GFP and 35S::ARG::GFP) on day 3 with/without ABA (100 μM) treatment was harvested and crosslinked for 15 min under a vacuum in crosslinking buffer (10 mMTris, pH8.0, 1 mM EDTA, 250 mM sucrose, 1 mM PMSF and 1% formaldehyde). Technical replicates containing approximately 1.5 mg DNA were resuspended in 800 μL SII buffer, incubated with 2 μg anti-GFP antibodies (ab290, Abcam) bound to Protein G Dynabeads (Invitrogen) for 1.5h at 4°C and washed five times with SII buffer. Chromatin was eluted from the beads twice at 65°C with Stop buffer (20 mM Tris-HCl, pH 8.0, 100 mM NaCl, 20 mM EDTA and 1% SDS). RNase- and DNase-free glycogen (2 μg) (Boehringer Mannheim) was added to the input and eluted chromatin, after which they were incubated with DNase- and RNase-free proteinase K (Invitrogen) at 65°C overnight and treated with 2 μg RNase A (Qiagen) for 1h at 37°C. DNA was purified using a Qiagen PCR Purification kit and resuspended in 100 μL H20. Quantitative PCR of the technical replicates was performed using a QuantiFast SYBR Green PCR kit (Qiagen) with the following cycling conditions: 2 min at 95°C, followed by 40 cycles of 15 s at 95°C, 15 s at 55°C and 20 s at 68°C. The primers used in this study are listed in Table S14. The results were normalized to the input DNA using the following equation: 100 × 2^(Ct input-Ct ChIP)^.

### Constructs and generation of transgenic Arabidopsis plants

Full-length cDNA of the selected genes *ARG*, *PRG*, *HRG*, *NRG* and *BRG* was amplified using the primers listed in Table S14. The amplified fragments were introduced into the pDONR221 vector (Invitrogen) and cloned into pMDC32 by LR reactions (Invitrogen). The construct was electroporated into *Agrobacterium tumefaciens* GV3101 and transformed into Col-0. Sterilized Arabidopsis seeds were grown on ½ MS medium with 1% sucrose solidified with 0.3% Phytagel under a 16 h/8 h light/dark cycle at 22°C. For drought stress treatment, 14-day-old T3 transgenic Arabidopsis plants were subjected to control (well-watered) and drought stress (withholding water) conditions for an additional 12 days. Two transformation events were used for the *ARG-ox* and *NRG-ox* plants, respectively. More than three pots were prepared for each transformation event, with 25 plants per pot, and all pots were rotated daily during drought stress treatment to minimize environmental effects. For the root elongation assays, the germinated seedlings were transferred to medium in square Petri dishes and grown vertically in long days for 7 days. For the PEG and ABA treatments, WT (Col-0), *ARG-ox* and *BRG-ox* homozygous Arabidopsis plants were grown in long days for 3 days, followed by 7 days of growth on ½ MS medium (1% sucrose, 0.3% Phytagel) supplemented with 10 μM ABA, 50 μM ABA or PEG (−1.7 Mpa) under continuous light or in the dark. At least 18–24 seedlings were measured for each genotype in each set of experiments. Total root length and root tip number were measured using Rhizo software (70). Primary root length was measured from scanned images using ImageJ software. Statistical analysis was performed in R (version 3.2.4) using two-way ANOVA + Tukey HSD for multiple comparisons as described in Generation of transgenic soybean hairy roots

## ABBREVIATIONS

ABA: abscisic acid
AUX: auxin
GA: gibberellic acid
ETH: ethylene
SA: salicylic acid
JA: jasmonic acid
CK: cytokinin
BR: brassinosteroid
*ARG*: **A**BRE binding factor-like **R**oot **G**rowth regulator
*BRG*: *BEE-like Root Growth regulator*
*PRG*: **P**hytochrome-interacting factor-like **R**oot Growth regulator
*NRG*: **N**F-YB-like **R**oot **G**rowth regulator
*HRG*: **H**AT-like **R**oot **G**rowth regulator
rdcTFs: root drought-related core TFs
*LUC*: firefly luciferase
*REN*: *Renilla* luciferase
HB: Homeodomain/HOMEOBOX
bZIP: basic leucine zipper
AP2/EREBP: APETALA2/ethylene-responsive element binding protein
C2H2: Cys2-His2 zinc finger
ARF: auxin response factor
VMS: very mild stress
MS: mild stress
SS: severe stress
GRN: Gene Regulatory Network
CREs: *cis*-regulatory elements

## DECLARATIONS

### Ethics approval and consent to participate

Not applicable

### Consent for publication

Not applicable

### Availability of data and materials

All data generated or analysed during this study are included in this published article [and its supplementary information files]. The microarray data sets used in this study were deposited in the Gene Expression Omnibus under the accession number of GSE102749 (http://www.ncbi.nlm.nih.gov/geo/query/acc.cgi?acc=GSE102749). The following secure token has been created to allow review of record GSE102749 while it remains in private status: gjgvekembzmdzcj.

### Authors’ contributions

L.L. designed the research. J.V.d.V., K.V., Y.A. and L.L. performed conserved motif analysis. N. N and L.L. contributed phenotype analysis. L.L., J.W., L.S., and W.C. performed the Dual-Luc assay. L.L., and G.P. performed the hairy root transformation. G.P. and L.L. contributed the *Arabidopsis* transformation. S.P., L.S., and M.M. performed the RNA-seq experiments and B.V. performed the microarray experiments. E.K., I.L. and L.L. contributed the co-functional TF-TF network. L.L. and G.P. constructed the TF library. L.L. performed Y1H assay and constructed TF-DNA networks. C.Z., G.K., T.J., L.L. and D.X. contributed bioinformatics analyses. L.L. wrote the article. H.N. and K.V. critically reviewed the manuscript.

## Acknowledgements

This work was funded by a United Soybean Board (project# 1204) grant to H.N. We thank Dr. Steve Kay for the pFLASH vector, Dr. Zhiyong Wang and Tao Wang for the comments and suggestions. K.V. acknowledges the Multidisciplinary Research Partnership “Bioinformatics: from nucleotides to networks” Project (no. 01MR0310W) of Ghent University. J.V.d.V is indebted to the Agency for Innovation by Science and Technology (IWT) in Flanders for a pre-doctoral fellowship.

