## Supplemental figures and tables for "Traceback of Core Transcription Factors for Soybean Root Growth Maintenance under Water Deficit"

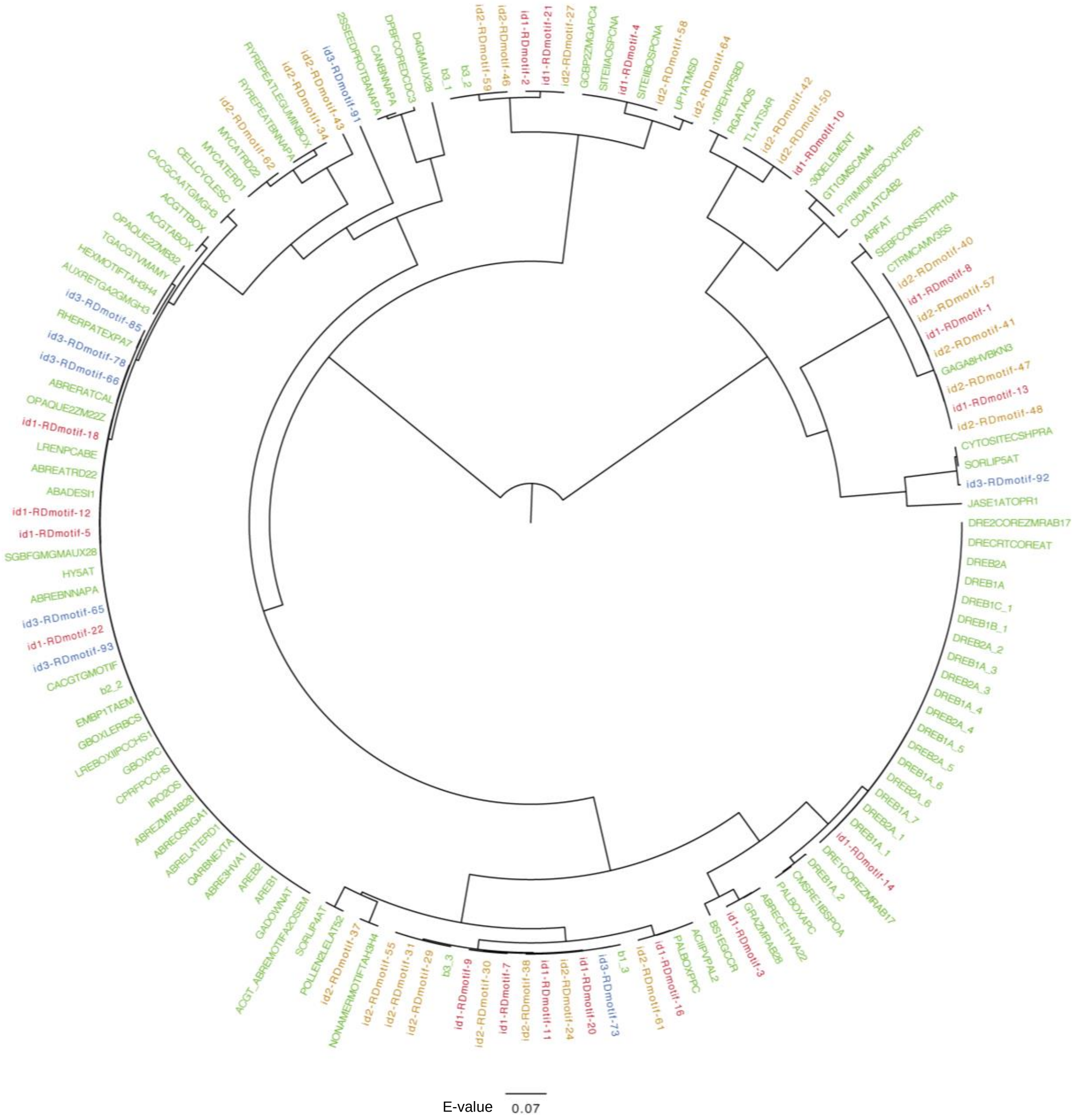


**Figure S1. Similarity tree of 49 conserved motifs with known motifs in the PLACE motif database.** The newly discovered motifs were compared to motifs from the PLACE database using the STAMP tool (Mahony and Benos, 2007). Motif similarity was determined with the Pearson correlation coefficient and aligned with the Ungapped Smith-Waterman algorithm. Green represents known motifs from the PLACE database http://www.dna.affrc.go.jp/PLACE/. Red represents the id1-motifs from cluster I. Yellow represents the id2-motifs from cluster II. Blue represents the id3-motifs from cluster III.

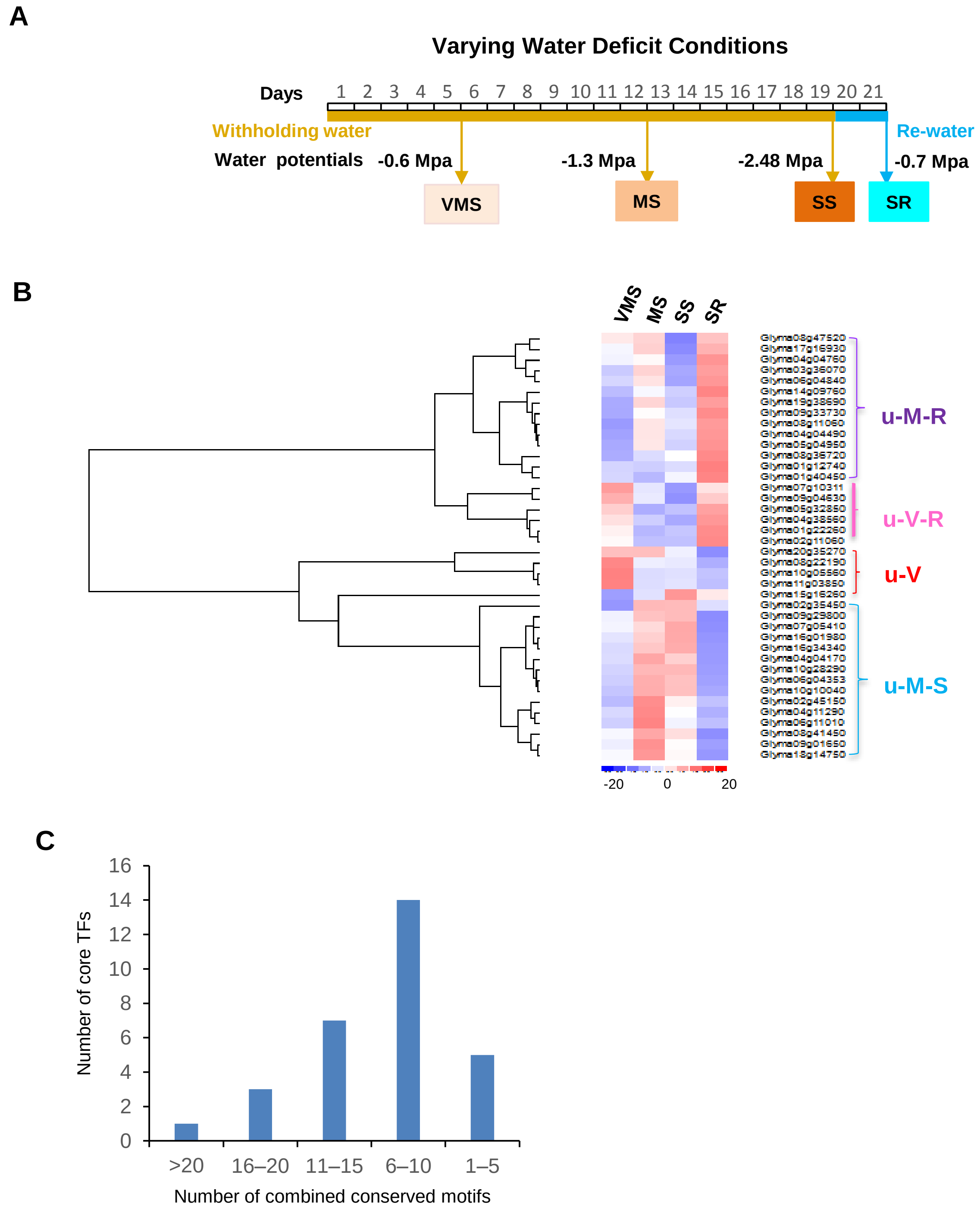


**Figure S2. Gene expression patterns of the core TFs under various water deficit conditions and the distribution of the 49 identified conserved motifs in the core TF genes.**

**(A)** Graphical description of various water deficit conditions in soybean primary roots. For very mild stress (VMS), drought stress was imposed by withholding water for 5 days. For mild stress (MS), water was withheld for 12 days. For severe stress (SS), water was withheld for 19 days. For water recovery after severe stress (SR), the plants were re-watered for 2 days after withholding water for 19 days. The water potential under each condition is indicated.

**(B)** Heatmap visualizing gene expression of the core TFs under different water-deficit conditions in soybean primary roots. u-V, genes upregulated under very mild stress (VMS). u-V-R, genes upregulated under VMS and water recovery after severe stress (SR). u-M-S, genes upregulated under mild stress (MS) and severe stress (SS). u-M-R, gene upregulated under MS and SR.

**(C)** Distribution of combined conserved motifs in the promoters of core TF genes. The 49 conserved motifs were identified in the 2-kb promoter regions of 35 core TF genes. The number of different motifs in the promoter of each core TF gene was counted. The number of core TFs was counted based on the range of combined motif number.

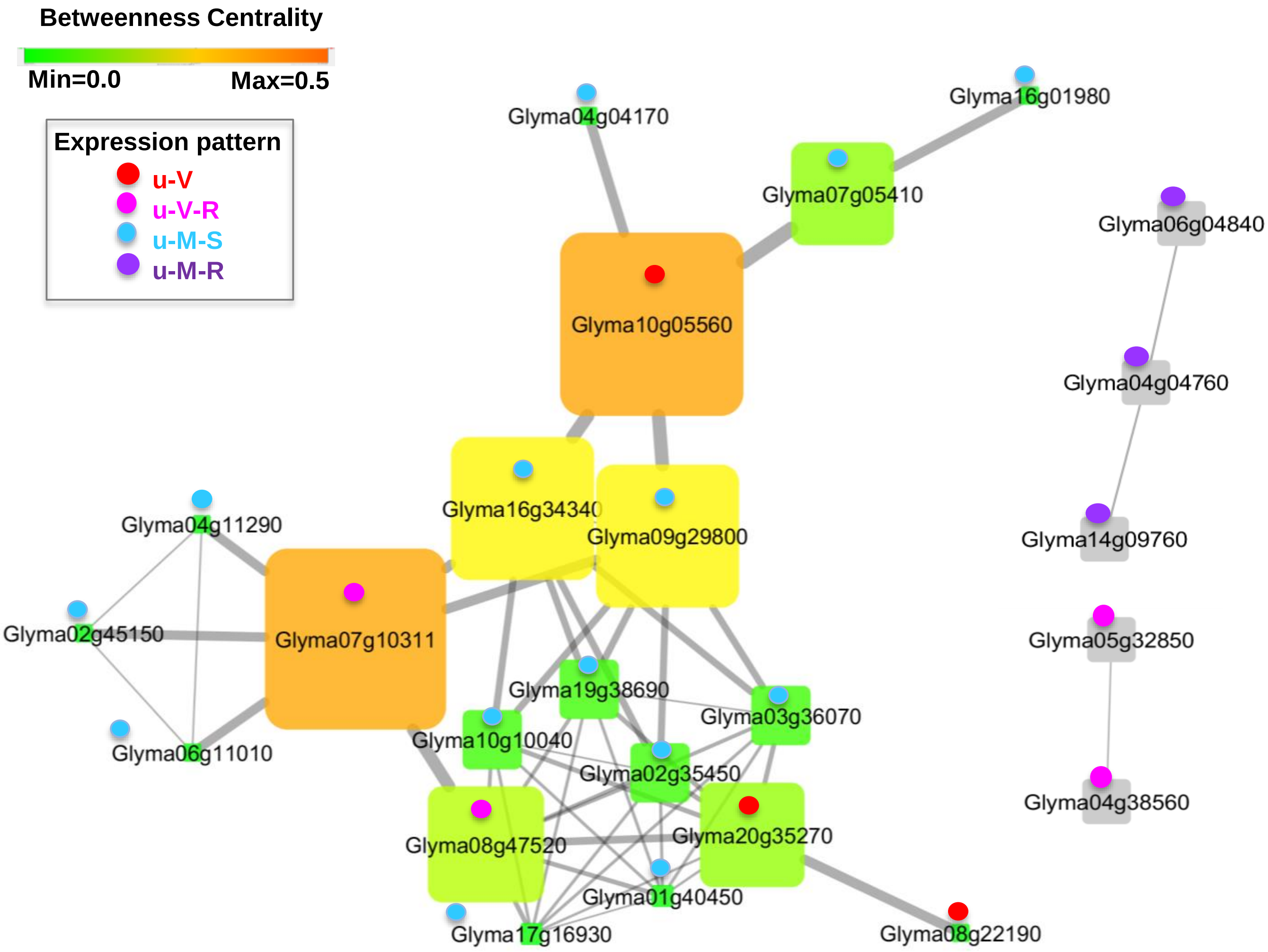


**Figure S3. Core TF co-functional network involved in the response to various water deficit conditions.**
A core TF co-functional network in soybean primary roots in response to water deficit. Each gene ID is shown in a colored box. The node size and the color intensity indicate the betweenness of centrality in the network to the hub genes. Edge betweenness is indicated by line thickness. The oval shape attached to each Gene ID represents the gene expression pattern under different water-deficit levels in soybean primary roots. An important hub gene, *Glyma01g10311*, was named *BEE-like Root Growth regulator* (*BRG)*. u-V (red), genes upregulated under VMS. u-V-R (pink), genes upregulated under VMS and SR. u-M-S (blue), genes upregulated under MS and SS. u-M-R (purple), gene upregulated under MS and SR. A heatmap of gene expression levels in roots under water deficit is shown in Figure S3A.

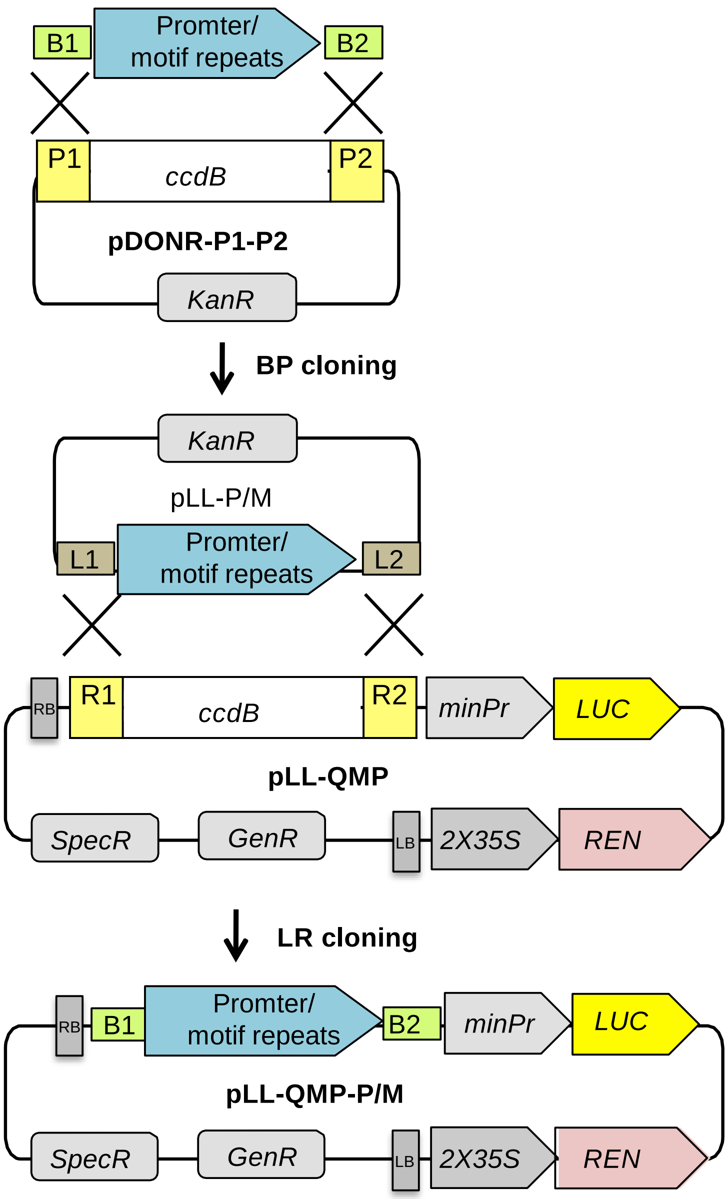


**Figure S4. Schematic diagram of dual-luciferase reporter vector construction.**
**(A)** A vector system was created to generate a single vector with the CaMV 35S mini promoter (*minPr*) fused to the firefly luciferase reporter gene (*LUC*) and the *35S* promoter fused to the *Renilla* luciferase reporter gene (*REN*). The constitutively expressed *Renilla* gene served as a control to normalize for transformation efficiency. Target motif repeats/promoter fragments were amplified from *G. max* genomic DNA using the appropriate primers with attB1 and attB2 sites. Each amplified fragment was cloned into the pDONR-P1-P2 vector by performing BP reactions to produce pLL-P/M. The fully functional expression vector pLL-QMP-PM was generated by Gateway LR cloning.
.

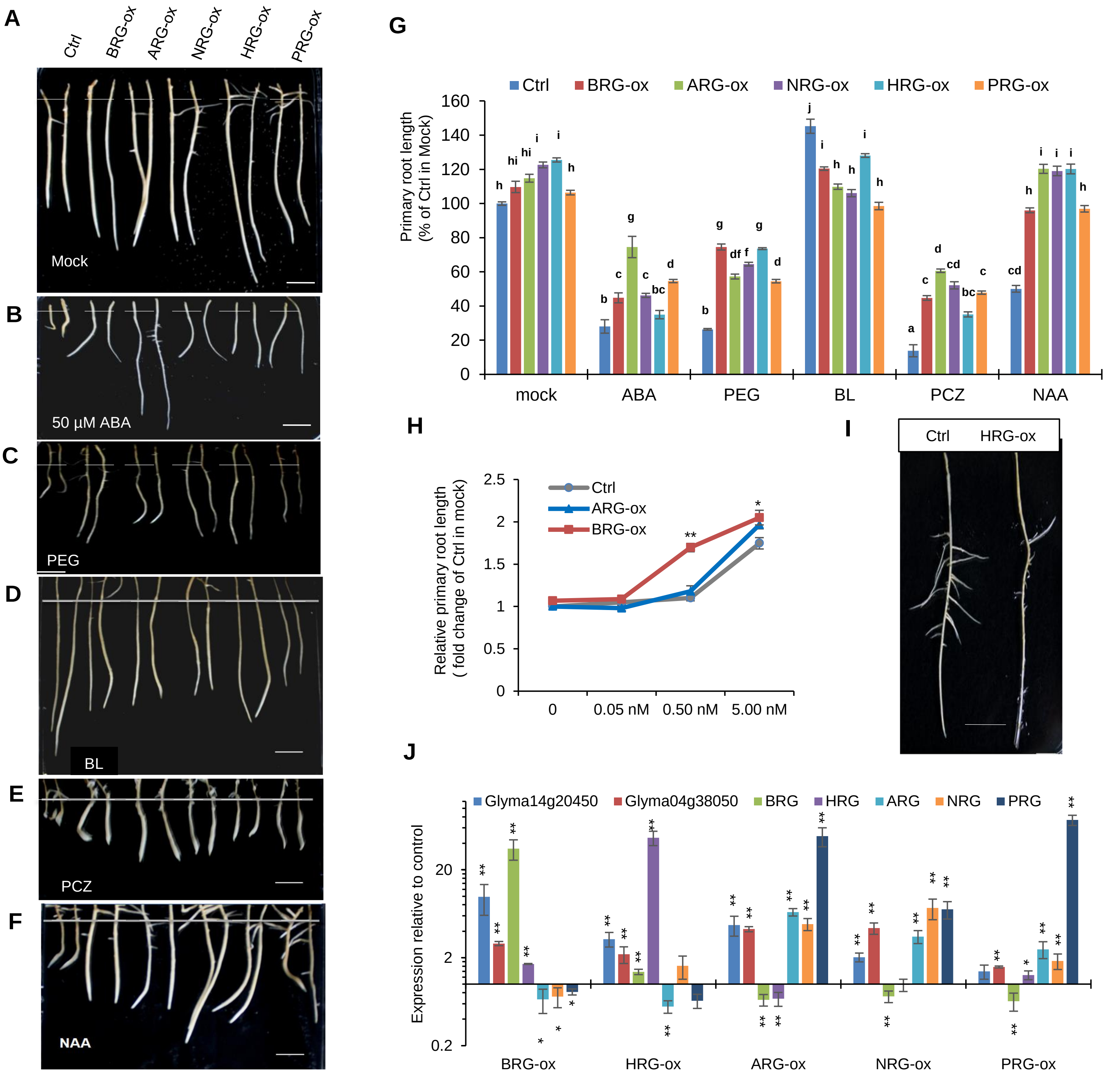

A
Ctrl
BRG-ox
ARG-ox
NRG-ox
HRG-ox
PRG-ox
Mock
B
50 µM ABA
C
PEG
D
BL
E
PCZ
F
NAA
G
Ctrl
BRG-ox
ARG-ox
NRG-ox
HRG-ox
PRG-ox
Primary root length (% of Ctrl in Mock)
mock
ABA
PEG
BL
PCZ
NAA
H
Relative primary root length ( fold change of Ctrl in mock)
Ctrl
ARG-ox
BRG-ox
0
0.05 nM
0.50 nM
5.00 nM
I
Ctrl
HRG-ox
J
Glyma14g20450
Glyma04g38050
BRG
HRG
ARG
NRG
PRG
Expression relative to control
BRG-ox
HRG-ox
ARG-ox
NRG-ox
PRG-ox

**Figure S5. Responses of transgenic soybean roots overexpressing *BRG*, *ARG*, *NRG, HRG* and *PRG* to different hormone or osmotic stress treatments**.

**(A)** Growth of roots from transgenic *BRG-ox, ARG-ox, NRG-ox, HRG-ox* and *PRG-ox* plants compared with the empty vector control (Ctrl) after 7 days on ½ MS solid medium (3% sucrose, 0.6% agar). Scale bar: 1 cm.

**(B)** The effect of 50 µM ABA on the length of transgenic roots.

**(C)** The effect of osmotic stress on the length of transgenic roots grown on PEG-infused plates (-1.2 Mpa).

**(D)** Effect of the hormone BR (0.5 nM BL) on the growth of transgenic roots. Note the hyposensitivity compared with the control.

**(E)** Effects of a BR inhibitor (10 µM PCZ) on the growth of transgenic roots. Note the lower sensitivity compared with the control.

**(F)** Effects of exogenous auxin on transgenic roots (50 µM NAA).

(B)–(F) Growth of transgenic roots compared with the empty vector control after 7 days on ½ MS solid medium supplemented with ABA, PEG, BL or PCZ. Scale bar: 1 cm.

**(G)** Quantification of the primary root growth of *BRG-ox, ARG-ox, NRG-ox, HRG-ox* and *PRG-ox* plants in response to 50 µM ABA, PEG (-1.2 Mpa), 0.5 nM BL, 10 µM PCZ, or 50 µM NAA treatment. Each sample from different root tips representing independent transformation events (mean ± SE; n=12). Comparisons are made between empty vector control plants and mutants prepared using the same growth conditions and the same treatments. Bars with the same letter are not significantly different (two-way ANOVA + Tukey HSD, $P < 0.01$, ANOVA table in Table S11 showing a significant genotype : hormone/PEG treatment interaction term) in response to ABA, PEG, BL, PCZ and NAA treatment.

**(H)** Transgenic *BRG-ox* roots show hypersensitivity to BR hormone (BL) at concentrations of 0.05 nM, 0.5 nM and 5 nM after three days of 10 µM PCZ treatments. Three biological samples, each from different root tips representing independent transformation events (mean ± SE; n=10). Significance relative to the empty vector control was determined by two-tailed Welch's *t*-test : single and double asterisks denote significance level at $P < 0.05$ and $P < 0.001$, respectively.

(I) Lateral root elongation in a wild-type plant (left) and a transgenic root overexpressing *HRG* at two weeks (right). Scale bar: 1 cm.

**(J)** RT-qPCR showing transcript abundance of *BRG, ARG, PRG, NRG* and *HRG* in *BRG-ox, ARG-ox, PRG-ox*, *NRG-ox* and *HRG-ox* relative to empty vector control. Root samples were collected after 7 days of culture on ½ MS solid medium. Expression was normalized to the *GmACT11* reference gene. *Glyma14g20450* and *Glyma04g38050* were used as positive drought-induced control genes (n=3 biological replicates with 3 technical replicates. Mean ± SE). Significance relative to the empty vector control was determined by two-tailed Welch's *t*-test : single and double asterisks denote significance level at $P < 0.05$ and $P < 0.001$, respectively.

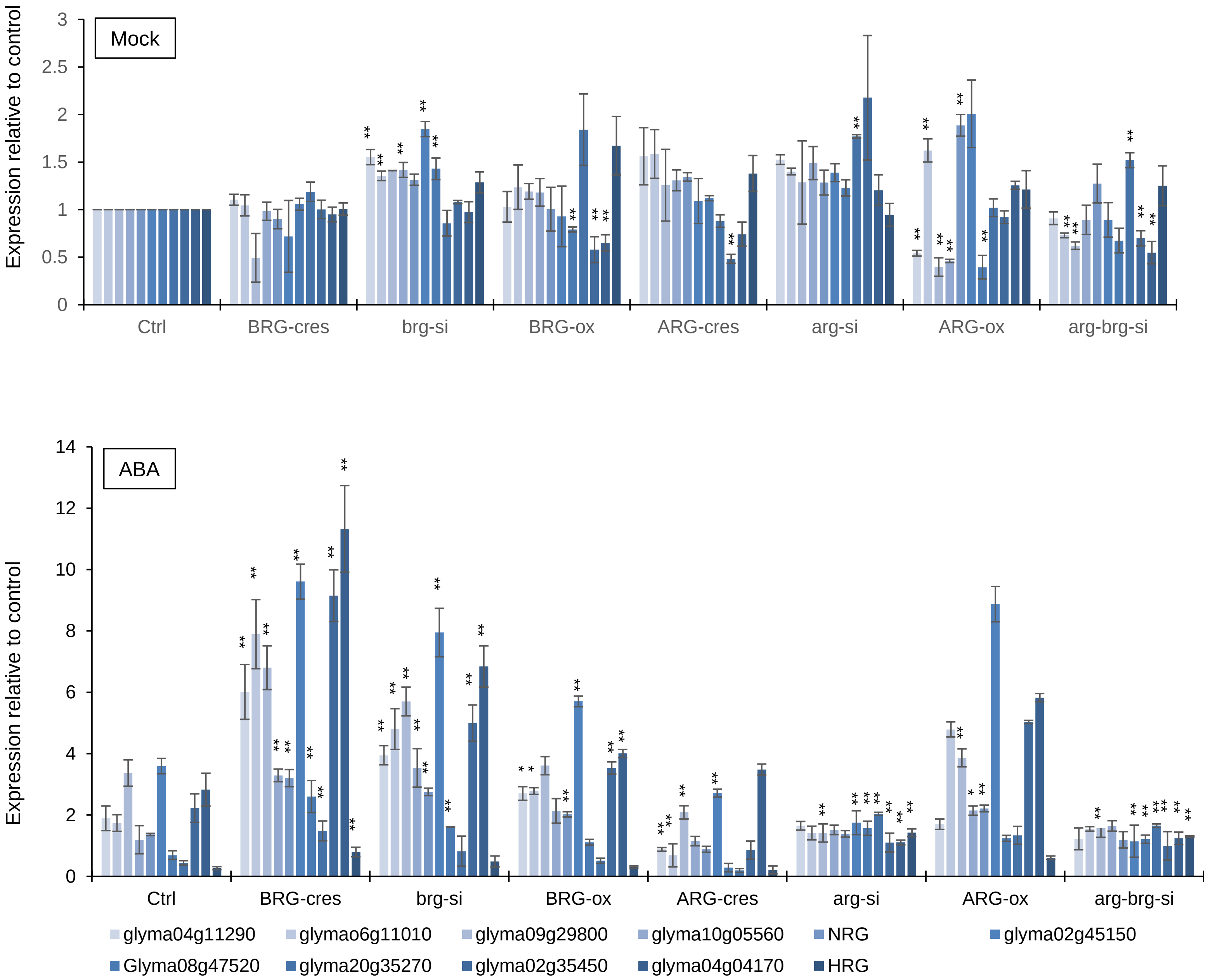


**Figure S6. RT-qPCR analysis showing fold differences in the expression of core TF genes in *BRG-ox*, *BRG-cres*, *brg-si*, *ARG-ox*, *ARG-cres* and *arg-si* relative to the empty vector control.**
Soybean root samples were collected after three days of culture on ½ MS solid medium with or without 100 µM ABA. *GmACT11* was used as a reference gene. All panels show data as n=3 biological replicates with 3 technical replicates (mean ± SE). Significance relative to the empty vector control under same condition was determined by two-tailed Welch's *t*-test: single and double asterisks denote significance level at P < 0.05 and P < 0.001, respectively.

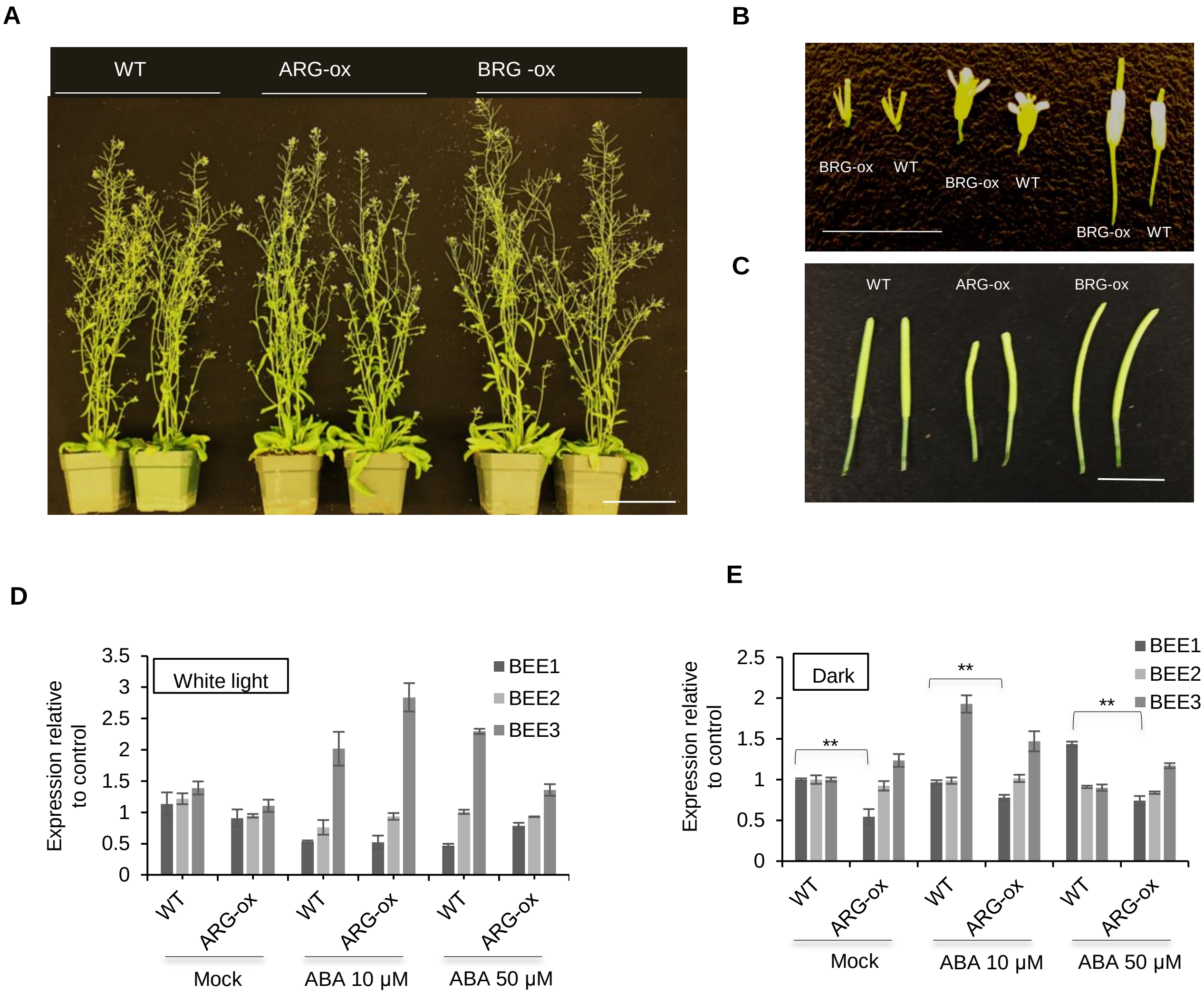


**Figure S7. Phenotypes of Arabidopsis plants overexpressing *BRG, ARG* and *NRG*.**
**(A)** Overall morphology of WT (Col-0), *BRG-ox* and *ARG-ox* Arabidopsis plants. Scale bar: 10 cm.
**(B)** The gynoecium and flower are larger in *BRG-ox* plants than in WT (Col-0). Scale bar: 1 cm.
**(C)** The siliques are larger in plants overexpressing *BRG* (*BRG-ox*) versus *ARG* (*ARG-ox*) than in WT (Col-0). Scale bar: 1 cm.
(D)- (E) RT-qPCR showing *BEE1*, *BEE2* and *BEE3* transcript levels in Arabidopsis root samples from WT and the *ARG-ox* mutant treated for 3 days with 50 µM ABA under continuous light or in the dark relative to mock-treated roots. Only *BEE1* was repressed in *ARG-ox* lines in the dark. Expression was normalized to the *At3g18780* (*ACT2*) reference gene. All panels show data as n=3 biological replicates with 3 technical replicates (mean ± SE). Significance relative to WT was determined by by two-tailed Welch's *t*-test: double asterisks denote significance level at $P < 0.001$.

**Table S1 Summary of Differentially Regulated Genes under water-deficit conditions across different root regions and time points.**

| Groups | Total | Up-regulated | Down-regulated |
|---|---|---|---|
| **D5hR1** | 998 | 477 | 521 |
| **D5hR2** | 2348 | 1282 | 1066 |
| **D48hR1** | 1400 | 1140 | 260 |
| **D48hR2** | 2307 | 2017 | 290 |
| **D48hR2vR3** | 4548 | 3409 | 1139 |

*P-value < 0.05 and genes with the regulation ratio log2 ≥ 2 or ≤ -2

**Table S2. List of 279 genes spatially-temporally regulated by water deficit conditions in soybean primary roots**

| Cluster | Gene ID |
|---|---|
| I | Glyma01g31750 |
| I | Glyma01g37630 |
| I | Glyma01g37810 |
| I | Glyma01g40320 |
| I | Glyma01g40450 |
| I | Glyma01g44530 |
| I | Glyma01g45000 |
| I | Glyma02g00820 |
| I | Glyma02g01520 |
| I | Glyma02g05930 |
| I | Glyma02g07070 |
| I | Glyma02g07370 |
| I | Glyma02g07530 |
| I | Glyma02g12470 |
| I | Glyma02g13800 |
| I | Glyma02g13850 |
| I | Glyma02g15520 |
| I | Glyma02g15780 |
| I | Glyma02g16170 |
| I | Glyma02g16520 |
| I | Glyma02g30800 |
| I | Glyma02g35630 |
| I | Glyma02g36200 |
| I | Glyma02g37710 |
| I | Glyma02g38250 |
| I | Glyma02g39750 |
| I | Glyma02g40040 |
| I | Glyma02g41070 |
| I | Glyma02g42290 |
| I | Glyma02g43860 |
| I | Glyma02g44690 |
| I | Glyma02g46030 |
| I | Glyma03g05050 |
| I | Glyma03g06360 |
| I | Glyma03g16600 |
| I | Glyma03g16620 |
| I | Glyma03g20630 |
| I | Glyma03g22320 |
| I | Glyma03g34110 |
| I | Glyma03g36070 |
| I | Glyma03g38810 |
| I | Glyma04g03150 |
| I | Glyma04g03510 |
| I | Glyma04g03750 |
| I | Glyma04g04730 |
| I | Glyma04g06460 |
| I | Glyma04g09310 |
| I | Glyma04g40530 |
| I | Glyma04g40580 |
| I | Glyma04g40590 |
| I | Glyma05g01330 |
| I | Glyma05g03650 |

| | |
|---|---|
| I | Glyma05g19920 |
| I | Glyma05g23150 |
| I | Glyma05g26070 |
| I | Glyma05g26930 |
| I | Glyma05g27030 |
| I | Glyma05g27640 |
| I | Glyma05g30690 |
| I | Glyma05g33080 |
| I | Glyma05g36110 |
| I | Glyma05g37420 |
| I | Glyma06g01290 |
| I | Glyma06g03200 |
| I | Glyma06g03450 |
| I | Glyma06g06500 |
| I | Glyma06g08990 |
| I | Glyma06g09470 |
| I | Glyma06g10970 |
| I | Glyma06g12270 |
| I | Glyma06g12580 |
| I | Glyma06g14160 |
| I | Glyma06g14220 |
| I | Glyma06g21170 |
| I | Glyma06g26370 |
| I | Glyma06g41520 |
| I | Glyma06g43940 |
| I | Glyma06g45090 |
| I | Glyma06g46680 |
| I | Glyma07g02540 |
| I | Glyma07g10311 |
| I | Glyma07g17000 |
| I | Glyma07g19120 |
| I | Glyma07g23260 |
| I | Glyma07g23480 |
| I | Glyma07g31200 |
| I | Glyma07g32980 |
| I | Glyma07g33460 |
| I | Glyma07g33950 |
| I | Glyma07g34830 |
| I | Glyma08g00480 |
| I | Glyma08g00640 |
| I | Glyma08g01620 |
| I | Glyma08g03540 |
| I | Glyma08g03850 |
| I | Glyma08g09910 |
| I | Glyma08g11620 |
| I | Glyma08g18470 |
| I | Glyma08g22670 |
| I | Glyma08g28980 |
| I | Glyma08g29100 |
| I | Glyma08g44630 |
| I | Glyma09g05170 |
| I | Glyma09g05850 |
| I | Glyma09g08270 |
| I | Glyma09g12040 |
| I | Glyma09g25590 |

| | |
|---|---|
| I | Glyma09g27320 |
| I | Glyma09g28800 |
| I | Glyma09g29510 |
| I | Glyma09g30190 |
| I | Glyma09g31220 |
| I | Glyma09g34980 |
| I | Glyma09g35050 |
| I | Glyma09g37120 |
| I | Glyma09g41190 |
| I | Glyma10g01560 |
| I | Glyma10g04690 |
| I | Glyma10g12130 |
| I | Glyma10g29280 |
| I | Glyma10g31350 |
| I | Glyma10g33860 |
| I | Glyma10g35550 |
| I | Glyma10g35650 |
| I | Glyma10g36570 |
| I | Glyma10g42820 |
| I | Glyma10g43420 |
| I | Glyma11g01010 |
| I | Glyma11g02960 |
| I | Glyma11g03500 |
| I | Glyma11g04840 |
| I | Glyma11g04970 |
| I | Glyma11g05800 |
| I | Glyma11g07670 |
| I | Glyma11g08320 |
| I | Glyma11g12720 |
| I | Glyma11g27480 |
| I | Glyma11g27720 |
| I | Glyma11g27900 |
| I | Glyma11g29340 |
| I | Glyma11g33300 |
| I | Glyma11g36000 |
| I | Glyma12g02540 |
| I | Glyma12g08520 |
| I | Glyma12g09430 |
| I | Glyma12g09810 |
| I | Glyma12g30990 |
| I | Glyma12g32270 |
| I | Glyma12g34310 |
| I | Glyma12g34550 |
| I | Glyma12g35070 |
| I | Glyma13g07900 |
| I | Glyma13g08070 |
| I | Glyma13g10070 |
| I | Glyma13g10140 |
| I | Glyma13g16440 |
| I | Glyma13g20510 |
| I | Glyma13g25280 |
| I | Glyma13g25890 |
| I | Glyma13g27110 |
| I | Glyma13g30290 |
| I | Glyma13g31450 |

| | |
|---|---|
| I | Glyma13g32310 |
| I | Glyma13g32320 |
| I | Glyma13g34520 |
| I | Glyma13g34550 |
| I | Glyma13g38150 |
| I | Glyma13g39320 |
| I | Glyma13g39330 |
| I | Glyma13g39710 |
| I | Glyma13g40450 |
| I | Glyma14g05960 |
| I | Glyma14g06600 |
| I | Glyma14g07990 |
| I | Glyma14g08030 |
| I | Glyma14g24260 |
| I | Glyma14g34640 |
| I | Glyma14g38170 |
| I | Glyma14g38210 |
| I | Glyma14g38390 |
| I | Glyma14g38580 |
| I | Glyma14g40680 |
| I | Glyma15g01100 |
| I | Glyma15g02210 |
| I | Glyma15g06010 |
| I | Glyma15g07890 |
| I | Glyma15g09430 |
| I | Glyma15g12480 |
| I | Glyma15g18640 |
| I | Glyma15g35070 |
| I | Glyma15g40510 |
| I | Glyma15g42980 |
| I | Glyma16g02960 |
| I | Glyma16g03780 |
| I | Glyma16g07830 |
| I | Glyma16g26060 |
| I | Glyma16g31280 |
| I | Glyma16g33950 |
| I | Glyma16g34100 |
| I | Glyma17g03360 |
| I | Glyma17g06220 |
| I | Glyma17g11110 |
| I | Glyma17g14180 |
| I | Glyma17g16930 |
| I | Glyma17g17210 |
| I | Glyma17g33450 |
| I | Glyma17g37020 |
| I | Glyma18g02980 |
| I | Glyma18g03530 |
| I | Glyma18g06740 |
| I | Glyma18g20610 |
| I | Glyma18g20990 |
| I | Glyma18g44650 |
| I | Glyma18g45260 |
| I | Glyma18g48230 |
| I | Glyma18g49560 |
| I | Glyma18g51880 |

| | |
|---|---|
| I | Glyma18g53970 |
| I | Glyma19g01100 |
| I | Glyma19g01120 |
| I | Glyma19g01450 |
| I | Glyma19g06540 |
| I | Glyma19g30460 |
| I | Glyma19g31460 |
| I | Glyma19g32700 |
| I | Glyma19g33330 |
| I | Glyma19g38690 |
| I | Glyma19g40470 |
| I | Glyma19g44610 |
| I | Glyma19g44980 |
| I | Glyma19g45120 |
| I | Glyma20g29000 |
| I | Glyma20g31030 |
| I | Glyma20g32140 |
| II | Glyma02g14910 |
| II | Glyma02g42990 |
| II | Glyma03g25640 |
| II | Glyma04g00920 |
| II | Glyma04g34080 |
| II | Glyma06g00950 |
| II | Glyma07g01960 |
| II | Glyma07g33570 |
| II | Glyma08g03260 |
| II | Glyma08g21630 |
| II | Glyma08g45130 |
| II | Glyma08g45140 |
| II | Glyma09g37090 |
| II | Glyma10g02130 |
| II | Glyma10g39760 |
| II | Glyma12g14130 |
| II | Glyma12g33600 |
| II | Glyma13g06130 |
| II | Glyma14g05300 |
| II | Glyma14g09710 |
| II | Glyma14g25590 |
| II | Glyma15g20700 |
| II | Glyma15g34770 |
| II | Glyma17g35460 |
| II | Glyma18g06610 |
| II | Glyma19g03570 |
| II | Glyma19g44410 |
| III | Glyma01g43780 |
| III | Glyma02g25930 |
| III | Glyma03g02980 |
| III | Glyma03g32520 |
| III | Glyma03g41580 |
| III | Glyma05g26390 |
| III | Glyma06g14850 |
| III | Glyma08g09300 |
| III | Glyma10g35280 |
| III | Glyma11g01730 |
| III | Glyma13g14190 |

| | |
|---|---|
| III | Glyma16g26070 |
| III | Glyma17g07010 |
| III | Glyma17g13550 |
| III | Glyma19g33310 |
| III | Glyma19g37860 |
| III | Glyma19g38390 |
| III | Glyma20g30460 |

*279 genes with more than two-fold changes in expression detected in both D48hR2 and D48hR2vR3. Hierarchical clustering of the 279 selected genes yielded three gene clusters: I, II and III.

### Table S3. List of the 49 identified conserved motifs

| Motif name | Motif regular expression | Fishers p value | Gene set matched conserved motif | Gene set matched motif | % of gene set conserved | Genomic matched conserved motif | Genomic matched motif | % of all genes conserved | species conserved |
|---|---|---|---|---|---|---|---|---|---|
| id1-RDmotif-1 | C[TC]CTC[TC]CT | 0.04595 | 42 | 97 | 43% | 5639 | 18144 | 31% | 6 |
| id1-RDmotif-2 | CC[CA][TGC]GCCC | 0.003977 | 1 | 23 | 4% | 106 | 2768 | 4% | 4 |
| id1-RDmotif-3 | G[AT]GGC[ATG]GC | 0.01259 | 3 | 30 | 10% | 238 | 4308 | 6% | 5 |
| id1-RDmotif-4 | GGG[GTA]CC[AC]A | 0.001392 | 8 | 42 | 19% | 476 | 5626 | 8% | 6 |
| id1-RDmotif-5 | CCACG[TA]G[TG] | 0.0001874 | 7 | 35 | 20% | 722 | 3944 | 18% | 6 |
| id1-RDmotif-7 | CCACCA[AC] | 0.01258 | 5 | 99 | 5% | 1068 | 17313 | 6% | 6 |
| id1-RDmotif-8 | [AG]GAGAGAGAG | 0.01042 | 17 | 29 | 59% | 2897 | 4064 | 71% | 6 |
| id1-RDmotif-9 | G[GA]GG[TC]GG[GT]GG | 0.01764 | 2 | 10 | 20% | 82 | 1030 | 8% | 3 |
| id1-RDmotif-10 | G[AG][AG]G[ATG][AGT]G[AG]A[GA] | 0.04466 | 11 | 146 | 8% | 1942 | 28068 | 7% | 6 |
| id1-RDmotif-11 | CC[CA][AC][CA]CAC[CA][CA] | 0.003533 | 11 | 50 | 22% | 427 | 7366 | 6% | 5 |
| id1-RDmotif-12 | CC[AC]C[GA][TA]G[TG]C | 0.0000617 | 5 | 35 | 14% | 491 | 3707 | 13% | 6 |
| id1-RDmotif-13 | [CT][TC]C[TA][CT][TC][CT][TC][CT][TC]C[TC][CT][TC]C | 0.007869 | 21 | 70 | 30% | 2499 | 11440 | 22% | 8 |
| id1-RDmotif-14 | GCCGACAA[AT]GC[AG] | 6.57E-09 | 3 | 5 | 60% | 4 | 11 | 36% | 4 |
| id1-RDmotif-16 | [CG][CA][CA]AC[CA][AC][AC]C[AC][CA][CA][CA][CA][CA] | 0.005136 | 7 | 31 | 23% | 186 | 4176 | 4% | 5 |
| id1-RDmotif-18 | CGTGGC | 0.00503 | 9 | 49 | 18% | 1215 | 7340 | 17% | 7 |
| id1-RDmotif-20 | [CG][CA]C[AC][CA]CAC | 0.02001 | 11 | 111 | 10% | 1021 | 20082 | 5% | 7 |
| id1-RDmotif-21 | CCCT[TG]CCC | 0.02244 | 1 | 20 | 5% | 133 | 2739 | 5% | 3 |
| id1-RDmotif-22 | CCACG[TG]G | 0.00704 | 6 | 38 | 16% | 985 | 5511 | 18% | 6 |
| id2-RDmotif-24 | [CA]CC[CA]C[CA]C | 0.01081 | 4 | 24 | 17% | 1816 | 24122 | 8% | 8 |
| id2-RDmotif-27 | GGGCAGGC | 0.01132 | 1 | 2 | 50% | 11 | 301 | 4% | 3 |
| id2-RDmotif-29 | TGGGG[TG]C | 0.0108 | 5 | 8 | 63% | 350 | 5331 | 7% | 5 |
| id2-RDmotif-30 | CGTGGTGG | 0.01561 | 2 | 3 | 67% | 50 | 971 | 5% | 4 |
| id2-RDmotif-31 | CAGTGGGG | 0.0436 | 2 | 2 | 100% | 22 | 628 | 4% | 3 |
| id2-RDmotif-34 | CATG[CA]A[CA]C | 0.0000356 | 2 | 18 | 11% | 247 | 9426 | 3% | 5 |
| id2-RDmotif-37 | G[AT][TC]CC[CA]AC | 0.001833 | 3 | 14 | 21% | 401 | 9553 | 4% | 5 |
| id2-RDmotif-38 | CCC[AC][CA]C[CA]C[TCA][TC] | 0.0002052 | 5 | 12 | 42% | 398 | 5767 | 7% | 4 |
| id2-RDmotif-40 | TCTCTCTC[TA] | 0.01537 | 7 | 13 | 54% | 4452 | 11452 | 39% | 7 |
| id2-RDmotif-41 | A[GA][TA]G[AT]G[AG]GTG | 0.03463 | 2 | 6 | 33% | 542 | 4410 | 12% | 4 |
| id2-RDmotif-42 | [TC][CT]CTTCTTC | 0.03161 | 2 | 13 | 15% | 1041 | 12840 | 8% | 3 |
| id2-RDmotif-43 | G[GA][GA][GT]CC[TAG]TG | 0.001926 | 2 | 10 | 20% | 159 | 5707 | 3% | 5 |
| id2-RDmotif-46 | [AG][CG][AG][AT]GG[GC]C[AG][ACG] | 0.0000552 | 2 | 14 | 14% | 189 | 6499 | 3% | 4 |
| id2-RDmotif-47 | C[TC][CT]T[CT]TCT[CT]T[CT]TCT[TC] | 0.003995 | 2 | 8 | 25% | 1343 | 4440 | 30% | 6 |
| id2-RDmotif-48 | [GA][GAT][GA][AG]A[AG]GA[ACG]A[AG]A[AG][AT]G | 0.009739 | 1 | 8 | 13% | 661 | 5227 | 13% | 6 |
| id2-RDmotif-50 | [TC][CT]CT[TC]CTTC | 0.04611 | 2 | 15 | 13% | 1316 | 16388 | 8% | 5 |
| id2-RDmotif-55 | CCCC[CA]C | 0.0253 | 7 | 18 | 39% | 1833 | 18768 | 10% | 8 |
| id2-RDmotif-57 | [TC]CTCTCTC | 0.02423 | 7 | 16 | 44% | 5622 | 16086 | 35% | 6 |
| id2-RDmotif-58 | G[TG]TCCCAC | 0.0000232 | 2 | 8 | 25% | 153 | 1968 | 8% | 3 |
| id2-RDmotif-59 | [AG][CG]AAGGG[CT] | 0.001572 | 2 | 14 | 14% | 241 | 9378 | 3% | 5 |
| id2-RDmotif-61 | [TG]GT[TC]G[GT]TG | 0.04447 | 1 | 17 | 6% | 792 | 19003 | 4% | 7 |
| id2-RDmotif-62 | [CT][CA]C[TA]TGG[CAG] | 0.02517 | 5 | 23 | 22% | 631 | 25300 | 2% | 5 |
| id2-RDmotif-64 | GAGGC[TC]C | 0.0009602 | 1 | 9 | 11% | 155 | 4304 | 4% | 5 |
| id3-RDmotif-65 | CACG[TG]G | 0.02913 | 3 | 14 | 21% | 1718 | 19889 | 9% | 8 |
| id3-RDmotif-66 | CCAC[CG][TC][GC]T | 0.0002251 | 2 | 11 | 18% | 724 | 7056 | 10% | 6 |
| id3-RDmotif-73 | C[ACG]C[CA]C[AC]CC | 0.02452 | 2 | 9 | 22% | 836 | 10678 | 8% | 6 |
| id3-RDmotif-78 | CCAC[GAC][CT]GT[GC][AC] | 7.93E-08 | 1 | 9 | 11% | 243 | 1742 | 14% | 4 |
| id3-RDmotif-85 | [CT][CA]AC[AC]CGTG[AC][CA] | 0.002132 | 1 | 3 | 33% | 28 | 677 | 4% | 3 |
| id3-RDmotif-91 | [GT][GA]A[AC][CGA]C[AGCT]T[GT][GC][CAT]AG[CT] | 0.00000682 | 2 | 6 | 33% | 18 | 1118 | 2% | 3 |
| id3-RDmotif-92 | TCC[CA]TC[AC]GT[CT][GAT][CT] | 0.000000441 | 3 | 5 | 60% | 10 | 368 | 3% | 3 |
| id3-RDmotif-93 | CACGTG | 0.04832 | 3 | 11 | 27% | 1932 | 16013 | 12% | 7 |

*Identificaiton of 49 conserved motifs in soybean via a comparison of soybean with 11 dicot and one monocot genome using multi-species phylogenetic footprinting. *p* -value is calculated based on how many times the algorithm observes conservation scores in the random groups that are as good as or better that the real conservation score from the original orthologous group.

**Table S4. List of the core set of drought-related TFs in soybean roots**

| Glyma 1.1 ID | Top TAIR10 Blastp Hit (4) | E-value (E<10-6 | Coverage | Gene Ontology Biological Process IDs (5) | Gene Ontology Biological Process Descriptions | PFAM_Descriptions | RM-TF | RR-TF |
|---|---|---|---|---|---|---|---|---|
| Glyma01g12740 | AT1G32640.1 \| Symbols: ATMYC2, RD22BP1, JAI1, JIN1, MYC2, ZBF1 \| Basic helix-loop-helix (bHLH) DNA-binding family protein \| chr1:11799042-11800913 REVERSE LENGTH=623 | 0 | 99.22 | GO:0006568 GO:0006612 GO:0009269 GO:0009611 GO:0009620 GO:0009694 GO:0009695 GO:0009737 GO:0009738 GO:0009753 GO:0009759 GO:0009863 GO:0009867 GO:0009963 GO:0010200 GO:0010363 GO:0043069 GO:0043619 GO:0045893 GO:0048364 GO:0051090 GO:2000068 | tryptophan metabolic process; "protein targeting to membrane"; "response to desiccation"; "response to wounding"; "response to fungus"; "jasmonic acid metabolic process"; "jasmonic acid biosynthetic process"; "response to abscisic acid stimulus"; "abscisic acid mediated signaling pathway"; "response to jasmonic acid stimulus"; "indole glucosinolate biosynthetic process"; "salicylic acid mediated signaling pathway"; "jasmonic acid mediated signaling pathway"; "positive regulation of flavonoid biosynthetic process"; "response to chitin"; "regulation of plant-type hypersensitive response"; "negative regulation of programmed cell death"; "regulation of transcription from RNA polymerase II promoter in response to oxidative stress"; "positive regulation of transcription, DNA-dependent"; "root development"; "regulation of sequence-specific DNA binding transcription factor activity"; "regulation of defense response to insect" | Helix-loop-helix DNA-binding domain | N | Y |
| Glyma01g22260 | AT1G25560.1 \| Symbols: TEM1, EDF1 \| AP2/B3 transcription factor family protein \| chr1:8981891-8982976 REVERSE LENGTH=361 | 2.00E-150 | 98.7 | GO:0006355 GO:0009873 GO:0030003 GO:0048573 GO:0070838 | regulation of transcription, DNA-dependent; "ethylene mediated signaling pathway"; "cellular cation homeostasis"; "photoperiodism, flowering"; "divalent metal ion transport" | AP2 domain; "B3 DNA binding domai | Y | Y |
| Glyma02g03020 | AT1G70000.2 \| Symbols: \| myb-like transcription factor family protein \| chr1:26363674-26364635 REVERSE LENGTH=261 | 4.00E-77 | 99.67 | GO:0006355 GO:0009651 GO:0009723 GO:0009733 GO:0009737 GO:0009739 GO:0009751 GO:0009753 GO:0046686 GO:0080167 | regulation of transcription, DNA-dependent; "response to salt stress"; "response to ethylene stimulus"; "response to auxin stimulus"; "response to abscisic acid stimulus"; "response to gibberellin stimulus"; "response to salicylic acid stimulus"; "response to jasmonic acid stimulus"; "response to cadmium ion"; "response to karrikin" | Myb-like DNA-binding domain | Y | N |
| Glyma02g11060 | AT1G25560.1 \| Symbols: TEM1, EDF1 \| AP2/B3 transcription factor family protein \| chr1:8981891-8982976 REVERSE LENGTH=361 | 2.00E-144 | 98.76 | GO:0006355 GO:0009873 GO:0030003 GO:0048573 GO:0070838 | regulation of transcription, DNA-dependent; "ethylene mediated signaling pathway"; "cellular cation homeostasis"; "photoperiodism, flowering"; "divalent metal ion transport" | B3 DNA binding domain; "AP2 domai | Y | Y |
| Glyma02g35450 | AT2G35940.3 \| Symbols: BLH1 \| BEL1-like homeodomain 1 \| chr2:15089171-15091699 REVERSE LENGTH=680 | 0 | 99.85 | GO:0000303 GO:0006355 GO:0006612 GO:0009610 GO:0009651 GO:0009733 GO:0009737 GO:0009743 GO:0009862 GO:0009873 GO:0009963 GO:0010197 GO:0010201 GO:0010363 GO:0019344 GO:0048513 | response to superoxide; "regulation of transcription, DNA-dependent"; "protein targeting to membrane"; "response to symbiotic fungus"; "response to salt stress"; "response to auxin stimulus"; "response to abscisic acid stimulus"; "response to carbohydrate stimulus"; "systemic acquired resistance, salicylic acid mediated signaling pathway"; "ethylene mediated signaling pathway"; "positive regulation of flavonoid biosynthetic process"; "polar nucleus fusion"; "response to continuous far red light stimulus by the high-irradiance response system"; "regulation of plant-type hypersensitive response"; "cysteine biosynthetic process"; "organ development" | Homeobox domain; "Associated with | Y | Y |
| Glyma02g45150 | AT2G43010.2 \| Symbols: PIF4, SRL2, AtPIF4 \| phytochrome interacting factor 4 \| chr2:17887003-17888823 FORWARD LENGTH=428 | 6.00E-41 | 55.95 | GO:0000165 GO:0006355 GO:0006612 GO:0007623 GO:0009617 GO:0009630 GO:0009704 GO:0009862 GO:0009867 GO:0010017 GO:0010161 GO:0010310 GO:0010363 GO:0010600 GO:0010928 GO:0030003 GO:0031348 GO:0035304 | MAPK cascade; "regulation of transcription, DNA-dependent"; "protein targeting to membrane"; "circadian rhythm"; "response to bacterium"; "gravitropism"; "de-etiolation"; "systemic acquired resistance, salicylic acid mediated signaling pathway"; "jasmonic acid mediated signaling pathway"; "red or far-red light signaling pathway"; "red light signaling pathway"; "regulation of hydrogen peroxide metabolic process"; "regulation of plant-type hypersensitive response"; "regulation of auxin biosynthetic process"; "regulation of auxin mediated signaling pathway"; "cellular cation homeostasis"; "negative regulation of defense response"; "regulation of protein dephosphorylation" | Helix-loop-helix DNA-binding domain | N | Y |

| | | | | | | | | |
|---|---|---|---|---|---|---|---|---|
| Glyma03g36070 | AT2G35940.3 \| Symbols: BLH1 \| BEL1-like homeodomain 1 \| chr2:15089171-15091699 REVERSE LENGTH=680 | 8.00E-172 | 99.71 | GO:0000303 GO:0006355 GO:0006612 GO:0009610 GO:0009651 GO:0009733 GO:0009737 GO:0009743 GO:0009862 GO:0009873 GO:0009963 GO:0010197 GO:0010201 GO:0010363 GO:0019344 GO:0048513 | response to superoxide; "regulation of transcription, DNA-dependent"; "protein targeting to membrane"; "response to symbiotic fungus"; "response to salt stress"; "response to auxin stimulus"; "response to abscisic acid stimulus"; "response to carbohydrate stimulus"; "systemic acquired resistance, salicylic acid mediated signaling pathway"; "ethylene mediated signaling pathway"; "positive regulation of flavonoid biosynthetic process"; "polar nucleus fusion"; "response to continuous far red light stimulus by the high-irradiance response system"; "regulation of plant-type hypersensitive response"; "cysteine biosynthetic process"; "organ development" | Associated with HOX; "Homeobox do | Y | Y |
| Glyma04g04170 | AT1G45249.1 \| Symbols: ABF2, AREB1, ATAREB1 \| abscisic acid responsive elements-binding factor 2 \| chr1:17165420-17167415 REVERSE LENGTH=416 | 6.00E-122 | 94.5 | GO:0006355 GO:0009414 GO:0009651 GO:0009737 GO:0009738 GO:0010255 GO:0045893 | regulation of transcription, DNA-dependent; "response to water deprivation"; "response to salt stress"; "response to abscisic acid stimulus"; "abscisic acid mediated signaling pathway"; "glucose mediated signaling pathway"; "positive regulation of transcription, DNA-dependent" | bZIP transcription factor | N | Y |
| Glyma04g04490 | AT4G37260.1 \| Symbols: MYB73, ATMYB73 \| myb domain protein 73 \| chr4:17540602-17541564 FORWARD LENGTH=320 | 6.00E-80 | 99.62 | GO:0006355 GO:0009723 GO:0009737 GO:0009751 GO:0009753 GO:0010200 GO:0046686 | regulation of transcription, DNA-dependent; "response to ethylene stimulus"; "response to abscisic acid stimulus"; "response to salicylic acid stimulus"; "response to jasmonic acid stimulus"; "response to chitin"; "response to cadmium ion" | Myb-like DNA-binding domain | Y | Y |
| Glyma04g04760 | AT1G27730.1 \| Symbols: STZ, ZAT10 \| salt tolerance zinc finger \| chr1:9648302-9648985 REVERSE LENGTH=227 | 6.00E-67 | 96.15 | GO:0002679 GO:0006979 GO:0007165 GO:0009409 GO:0009414 GO:0009611 GO:0009612 GO:0009620 GO:0009644 GO:0009651 GO:0009695 GO:0009723 GO:0009733 GO:0009737 GO:0009738 GO:0009753 GO:0009873 GO:0010117 GO:0010200 GO:0015979 GO:0035264 GO:0035556 GO:0042538 GO:0045892 | respiratory burst involved in defense response; "response to oxidative stress"; "signal transduction"; "response to cold"; "response to water deprivation"; "response to wounding"; "response to mechanical stimulus"; "response to fungus"; "response to high light intensity"; "response to salt stress"; "jasmonic acid biosynthetic process"; "response to ethylene stimulus"; "response to auxin stimulus"; "response to abscisic acid stimulus"; "abscisic acid mediated signaling pathway"; "response to jasmonic acid stimulus"; "ethylene mediated signaling pathway"; "photoprotection"; "response to chitin"; "photosynthesis"; "multicellular organism growth"; "intracellular signal transduction"; "hyperosmotic salinity response"; "negative regulation of transcription, DNA-dependent" | C2H2-like | Y | Y |
| Glyma04g11290 | AT1G78080.1 \| Symbols: RAP2.4 \| related to AP2 4 \| chr1:29364790-29365794 FORWARD LENGTH=334 | 1.00E-96 | 99.68 | GO:0006355 GO:0006970 GO:0009409 GO:0009414 GO:0009416 GO:0009611 GO:0009651 GO:0009736 GO:0009873 GO:0010017 GO:0045595 GO:0071472 | regulation of transcription, DNA-dependent; "response to osmotic stress"; "response to cold"; "response to water deprivation"; "response to light stimulus"; "response to wounding"; "response to salt stress"; "cytokinin mediated signaling pathway"; "ethylene mediated signaling pathway"; "red or far-red light signaling pathway"; "regulation of cell differentiation"; "cellular response to salt stress" | AP2 domain | Y | Y |
| Glyma04g38560 | AT1G01720.1 \| Symbols: ATAF1, ANAC002 \| NAC (No Apical Meristem) domain transcriptional regulator superfamily protein \| chr1:268471-269514 FORWARD LENGTH=289 | 1.00E-121 | 99.32 | GO:0006355 GO:0007275 GO:0009414 GO:0009611 GO:0009620 GO:0009695 GO:0009737 GO:0009753 GO:0009788 GO:0010200 GO:0042538 | regulation of transcription, DNA-dependent; "multicellular organismal development"; "response to water deprivation"; "response to wounding"; "response to fungus"; "jasmonic acid biosynthetic process"; "response to abscisic acid stimulus"; "response to jasmonic acid stimulus"; "negative regulation of abscisic acid mediated signaling pathway"; "response to chitin"; "hyperosmotic salinity response" | No apical meristem (NAM) protein | Y | Y |

| | | | | | | | | |
|---|---|---|---|---|---|---|---|---|
| Glyma05g04950 | AT5G47390.1 \| Symbols: \| myb-like transcription factor family protein \| chr5:19227001-19228546 FORWARD LENGTH=365 | 2.00E-142 | 99.7 | GO:0006355 GO:0009651 GO:0009723 GO:0009737 GO:0009739 GO:0009751 GO:0009753 GO:0046686 | regulation of transcription, DNA-dependent; "response to salt stress"; "response to ethylene stimulus"; "response to abscisic acid stimulus"; "response to gibberellin stimulus"; "response to salicylic acid stimulus"; "response to jasmonic acid stimulus"; "response to cadmium ion" | Myb-like DNA-binding domain | Y | Y |
| Glyma05g32850 | AT1G01720.1 \| Symbols: ATAF1, ANAC002 \| NAC (No Apical Meristem) domain transcriptional regulator superfamily protein \| chr1:268471-269514 FORWARD LENGTH=289 | 5.00E-121 | 98.33 | GO:0006355 GO:0007275 GO:0009414 GO:0009611 GO:0009620 GO:0009695 GO:0009737 GO:0009753 GO:0009788 GO:0010200 GO:0042538 | regulation of transcription, DNA-dependent; "multicellular organismal development"; "response to water deprivation"; "response to wounding"; "response to fungus"; "jasmonic acid biosynthetic process"; "response to abscisic acid stimulus"; "response to jasmonic acid stimulus"; "negative regulation of abscisic acid mediated signaling pathway"; "response to chitin"; "hyperosmotic salinity response" | No apical meristem (NAM) protein | Y | Y |
| Glyma06g04840 | AT1G27730.1 \| Symbols: STZ, ZAT10 \| salt tolerance zinc finger \| chr1:9648302-9648985 REVERSE LENGTH=227 | 1.00E-66 | 96.15 | GO:0002679 GO:0006979 GO:0007165 GO:0009409 GO:0009414 GO:0009611 GO:0009612 GO:0009620 GO:0009644 GO:0009651 GO:0009695 GO:0009723 GO:0009733 GO:0009737 GO:0009738 GO:0009753 GO:0009873 GO:0010117 GO:0010200 GO:0015979 GO:0035264 GO:0035556 GO:0042538 GO:0045892 | respiratory burst involved in defense response; "response to oxidative stress"; "signal transduction"; "response to cold"; "response to water deprivation"; "response to wounding"; "response to mechanical stimulus"; "response to fungus"; "response to high light intensity"; "response to salt stress"; "jasmonic acid biosynthetic process"; "response to ethylene stimulus"; "response to auxin stimulus"; "response to abscisic acid stimulus"; "abscisic acid mediated signaling pathway"; "response to jasmonic acid stimulus"; "ethylene mediated signaling pathway"; "photoprotection"; "response to chitin"; "photosynthesis"; "multicellular organism growth"; "intracellular signal transduction"; "hyperosmotic salinity response"; "negative regulation of transcription, DNA-dependent" | C2H2-like | N | Y |
| Glyma06g11010 | AT1G78080.1 \| Symbols: RAP2.4 \| related to AP2 4 \| chr1:29364790-29365794 FORWARD LENGTH=334 | 7.00E-100 | 99.67 | GO:0006355 GO:0006970 GO:0009409 GO:0009414 GO:0009416 GO:0009611 GO:0009651 GO:0009736 GO:0009873 GO:0010017 GO:0045595 GO:0071472 | regulation of transcription, DNA-dependent; "response to osmotic stress"; "response to cold"; "response to water deprivation"; "response to light stimulus"; "response to wounding"; "response to salt stress"; "cytokinin mediated signaling pathway"; "ethylene mediated signaling pathway"; "red or far-red light signaling pathway"; "regulation of cell differentiation"; "cellular response to salt stress" | AP2 domain | Y | Y |
| Glyma07g05410 | AT1G01060.3 \| Symbols: LHY, LHY1 \| Homeodomain-like superfamily protein \| chr1:33992-37061 REVERSE LENGTH=645 | 1.00E-145 | 99.87 | GO:0006355 GO:0007623 GO:0009409 GO:0009639 GO:0009651 GO:0009723 GO:0009733 GO:0009737 GO:0009739 GO:0009751 GO:0009753 GO:0042752 GO:0042754 GO:0043433 GO:0046686 GO:0048574 | regulation of transcription, DNA-dependent; "circadian rhythm"; "response to cold"; "response to red or far red light"; "response to salt stress"; "response to ethylene stimulus"; "response to auxin stimulus"; "response to abscisic acid stimulus"; "response to gibberellin stimulus"; "response to salicylic acid stimulus"; "response to jasmonic acid stimulus"; "regulation of circadian rhythm"; "negative regulation of circadian rhythm"; "negative regulation of sequence-specific DNA binding transcription factor activity"; "response to cadmium ion"; "long-day photoperiodism, flowering" | Myb-like DNA-binding domain | Y | Y |

| | | | | | | | | |
|---|---|---|---|---|---|---|---|---|
| Glyma08g10140 | AT1G14920.1 \| Symbols: GAI, RGA2 \| GRAS family transcription factor family protein \| chr1:5149414-5151015 FORWARD LENGTH=533 | 0 | 94.21 | GO:0006808 GO:0009651 GO:0009723 GO:0009737 GO:0009739 GO:0009740 GO:0009845 GO:0009863 GO:0009867 GO:0009938 GO:0010029 GO:0010162 GO:0010187 GO:0010218 GO:0010233 GO:0010325 GO:0042538 GO:0048444 GO:2000033 GO:2000377 | regulation of nitrogen utilization; "response to salt stress"; "response to ethylene stimulus"; "response to abscisic acid stimulus"; "response to gibberellin stimulus"; "gibberellic acid mediated signaling pathway"; "seed germination"; "salicylic acid mediated signaling pathway"; "jasmonic acid mediated signaling pathway"; "negative regulation of gibberellic acid mediated signaling pathway"; "regulation of seed germination"; "seed dormancy process"; "negative regulation of seed germination"; "response to far red light"; "phloem transport"; "raffinose family oligosaccharide biosynthetic process"; "hyperosmotic salinity response"; "floral organ morphogenesis"; "regulation of seed dormancy process"; "regulation of reactive oxygen species metabolic process" | GRAS family transcription factor; "Tra | Y | N |
| Glyma08g22190 | AT3G15540.1 \| Symbols: IAA19, MSG2 \| indole-3-acetic acid inducible 19 \| chr3:5264100-5265378 FORWARD LENGTH=197 | 7.00E-75 | 93.88 | GO:0006355 GO:0009630 GO:0009638 GO:0009733 GO:0009741 GO:0048527 GO:0080086 | regulation of transcription, DNA-dependent; "gravitropism"; "phototropism"; "response to auxin stimulus"; "response to brassinosteroid stimulus"; "lateral root development"; "stamen filament development" | AUX/IAA family | Y | Y |
| Glyma08g36720 | AT1G32640.1 \| Symbols: ATMYC2, RD22BP1, JAI1, JIN1, MYC2, ZBF1 \| Basic helix-loop-helix (bHLH) DNA-binding family protein \| chr1:11799042-11800913 REVERSE LENGTH=623 | 0 | 99.24 | GO:0006568 GO:0006612 GO:0009269 GO:0009611 GO:0009620 GO:0009694 GO:0009695 GO:0009737 GO:0009738 GO:0009753 GO:0009759 GO:0009863 GO:0009867 GO:0009963 GO:0010200 GO:0010363 GO:0043069 GO:0043619 GO:0045893 GO:0048364 GO:0051090 GO:2000068 | tryptophan metabolic process; "protein targeting to membrane"; "response to desiccation"; "response to wounding"; "response to fungus"; "jasmonic acid metabolic process"; "jasmonic acid biosynthetic process"; "response to abscisic acid stimulus"; "abscisic acid mediated signaling pathway"; "response to jasmonic acid stimulus"; "indole glucosinolate biosynthetic process"; "salicylic acid mediated signaling pathway"; "jasmonic acid mediated signaling pathway"; "positive regulation of flavonoid biosynthetic process"; "response to chitin"; "regulation of plant-type hypersensitive response"; "negative regulation of programmed cell death"; "regulation of transcription from RNA polymerase II promoter in response to oxidative stress"; "positive regulation of transcription, DNA-dependent"; "root development"; "regulation of sequence-specific DNA binding transcription factor activity"; "regulation of defense response to insect" | Helix-loop-helix DNA-binding domain | N | Y |
| Glyma08g47520 | AT5G13180.1 \| Symbols: ANAC083, VNI2, NAC083 \| NAC domain containing protein 83 \| chr5:4196643-4197577 FORWARD LENGTH=252 | 1.00E-93 | 96 | GO:0006355 GO:0006612 GO:0007275 GO:0009651 GO:0009737 GO:0009741 GO:0009963 GO:0010089 GO:0010150 GO:0010363 GO:0045892 | regulation of transcription, DNA-dependent; "protein targeting to membrane"; "multicellular organismal development"; "response to salt stress"; "response to abscisic acid stimulus"; "response to brassinosteroid stimulus"; "positive regulation of flavonoid biosynthetic process"; "xylem development"; "leaf senescence"; "regulation of plant-type hypersensitive response"; "negative regulation of transcription, DNA-dependent" | No apical meristem (NAM) protein | Y | Y |
| Glyma09g04630 | AT3G16770.1 \| Symbols: RAP2.3, ATEBP, ERF72, EBP \| ethylene-responsive element binding protein \| chr3:5705784-5706768 FORWARD LENGTH=248 | 1.00E-47 | 96.22 | GO:0006355 GO:0008219 GO:0009723 GO:0009735 GO:0009753 GO:0009873 GO:0010286 GO:0045893 GO:0051707 | regulation of transcription, DNA-dependent; "cell death"; "response to ethylene stimulus"; "response to cytokinin stimulus"; "response to jasmonic acid stimulus"; "ethylene mediated signaling pathway"; "heat acclimation"; "positive regulation of transcription, DNA-dependent"; "response to other organism" | AP2 domain | Y | Y |
| Glyma09g29800 | AT1G18330.1 \| Symbols: EPR1, RVE7 \| Homeodomain-like superfamily protein \| chr1:6306196-6307640 REVERSE LENGTH=346 | 4.00E-61 | 60.81 | GO:0006355 GO:0007623 GO:0009651 GO:0009723 GO:0046686 | regulation of transcription, DNA-dependent; "circadian rhythm"; "response to salt stress"; "response to ethylene stimulus"; "response to cadmium ion" | Myb-like DNA-binding domain | Y | Y |

| | | | | | | | | |
|---|---|---|---|---|---|---|---|---|
| Glyma09g33730 | AT1G32640.1 \| Symbols: ATMYC2, RD22BP1, JAI1, JIN1, MYC2, ZBF1 \| Basic helix-loop-helix (bHLH) DNA-binding family protein \| chr1:11799042-11800913 REVERSE LENGTH=623 | 0 | 90.44 | GO:0006568<br>GO:0006612<br>GO:0009269<br>GO:0009611<br>GO:0009620<br>GO:0009694<br>GO:0009695<br>GO:0009737<br>GO:0009738<br>GO:0009753<br>GO:0009759<br>GO:0009863<br>GO:0009867<br>GO:0009963<br>GO:0010200<br>GO:0010363<br>GO:0043069<br>GO:0043619<br>GO:0045893<br>GO:0048364<br>GO:0051090<br>GO:2000068 | tryptophan metabolic process; "protein targeting to membrane"; "response to desiccation"; "response to wounding"; "response to fungus"; "jasmonic acid metabolic process"; "jasmonic acid biosynthetic process"; "response to abscisic acid stimulus"; "abscisic acid mediated signaling pathway"; "response to jasmonic acid stimulus"; "indole glucosinolate biosynthetic process"; "salicylic acid mediated signaling pathway"; "jasmonic acid mediated signaling pathway"; "positive regulation of flavonoid biosynthetic process"; "response to chitin"; "regulation of plant-type hypersensitive response"; "negative regulation of programmed cell death"; "regulation of transcription from RNA polymerase II promoter in response to oxidative stress"; "positive regulation of transcription, DNA-dependent"; "root development"; "regulation of sequence-specific DNA binding transcription factor activity"; "regulation of defense response to insect" | Helix-loop-helix DNA-binding domain | Y | Y |
| Glyma10g05560 | AT3G09600.1 \| Symbols: \| Homeodomain-like superfamily protein \| chr3:2946459-2948270 FORWARD LENGTH=298 | 5.00E-134 | 87.46 | GO:0006355<br>GO:0007623<br>GO:0009651<br>GO:0009723<br>GO:0009733<br>GO:0009737<br>GO:0009739<br>GO:0009751<br>GO:0009753<br>GO:0032922<br>GO:0043966<br>GO:0046686<br>GO:0048573<br>GO:0048574 | regulation of transcription, DNA-dependent; "circadian rhythm"; "response to salt stress"; "response to ethylene stimulus"; "response to auxin stimulus"; "response to abscisic acid stimulus"; "response to gibberellin stimulus"; "response to salicylic acid stimulus"; "response to jasmonic acid stimulus"; "circadian regulation of gene expression"; "histone H3 acetylation"; "response to cadmium ion"; "photoperiodism, flowering"; "long-day photoperiodism, flowering" | Myb-like DNA-binding domain | Y | Y |
| Glyma10g10040 | AT2G35940.3 \| Symbols: BLH1 \| BEL1-like homeodomain 1 \| chr2:15089171-15091699 REVERSE LENGTH=680 | 1.00E-176 | 99.85 | GO:0000303<br>GO:0006355<br>GO:0006612<br>GO:0009610<br>GO:0009651<br>GO:0009733<br>GO:0009737<br>GO:0009743<br>GO:0009862<br>GO:0009873<br>GO:0009963<br>GO:0010197<br>GO:0010201<br>GO:0010363<br>GO:0019344<br>GO:0048513 | response to superoxide; "regulation of transcription, DNA-dependent"; "protein targeting to membrane"; "response to symbiotic fungus"; "response to salt stress"; "response to auxin stimulus"; "response to abscisic acid stimulus"; "response to carbohydrate stimulus"; "systemic acquired resistance, salicylic acid mediated signaling pathway"; "ethylene mediated signaling pathway"; "positive regulation of flavonoid biosynthetic process"; "polar nucleus fusion"; "response to continuous far red light stimulus by the high-irradiance response system"; "regulation of plant-type hypersensitive response"; "cysteine biosynthetic process"; "organ development" | Associated with HOX; "Homeobox do | Y | Y |
| Glyma11g33720 | AT3G03450.1 \| Symbols: RGL2 \| RGA-like 2 \| chr3:819636-821279 REVERSE LENGTH=547 | 0 | 88.76 | GO:0009062<br>GO:0009651<br>GO:0009686<br>GO:0009723<br>GO:0009737<br>GO:0009739<br>GO:0009740<br>GO:0009793<br>GO:0009845<br>GO:0009863<br>GO:0009867<br>GO:0009938<br>GO:0010029<br>GO:0010162<br>GO:0010187<br>GO:0010325<br>GO:0042538<br>GO:0048444<br>GO:0048608<br>GO:2000033<br>GO:2000377 | fatty acid catabolic process; "response to salt stress"; "gibberellin biosynthetic process"; "response to ethylene stimulus"; "response to abscisic acid stimulus"; "response to gibberellin stimulus"; "gibberellic acid mediated signaling pathway"; "embryo development ending in seed dormancy"; "seed germination"; "salicylic acid mediated signaling pathway"; "jasmonic acid mediated signaling pathway"; "negative regulation of gibberellic acid mediated signaling pathway"; "regulation of seed germination"; "seed dormancy process"; "negative regulation of seed germination"; "raffinose family oligosaccharide biosynthetic process"; "hyperosmotic salinity response"; "floral organ morphogenesis"; "reproductive structure development"; "regulation of seed dormancy process"; "regulation of reactive oxygen species metabolic process" | GRAS family transcription factor; "Tra | Y | N |
| Glyma12g06680 | AT1G07540.1 \| Symbols: TRFL2 \| TRF-like 2 \| chr1:2318433-2321048 REVERSE LENGTH=630 | 9.00E-115 | 68.68 | NA | NA | Myb-like DNA-binding domain | N | N |

| | | | | | | | | |
|---|---|---|---|---|---|---|---|---|
| Glyma14g09760 | AT1G27730.1 \| Symbols: STZ, ZAT10 \| salt tolerance zinc finger \| chr1:9648302-9648985 REVERSE LENGTH=227 | 4.00E-62 | 94.02 | GO:0002679 GO:0006979 GO:0007165 GO:0009409 GO:0009414 GO:0009611 GO:0009612 GO:0009620 GO:0009644 GO:0009651 GO:0009695 GO:0009723 GO:0009733 GO:0009737 GO:0009738 GO:0009753 GO:0009873 GO:0010117 GO:0010200 GO:0015979 GO:0035264 GO:0035556 GO:0042538 GO:0045892 | respiratory burst involved in defense response; "response to oxidative stress"; "signal transduction"; "response to cold"; "response to water deprivation"; "response to wounding"; "response to mechanical stimulus"; "response to fungus"; "response to high light intensity"; "response to salt stress"; "jasmonic acid biosynthetic process"; "response to ethylene stimulus"; "response to auxin stimulus"; "response to abscisic acid stimulus"; "abscisic acid mediated signaling pathway"; "response to jasmonic acid stimulus"; "ethylene mediated signaling pathway"; "photoprotection"; "response to chitin"; "photosynthesis"; "multicellular organism growth"; "intracellular signal transduction"; "hyperosmotic salinity response"; "negative regulation of transcription, DNA-dependent" | C2H2-like | Y | Y |
| Glyma15g16260 | AT3G16770.1 \| Symbols: RAP2.3, ATEBP, ERF72, EBP \| ethylene-responsive element binding protein \| chr3:5705784-5706768 FORWARD LENGTH=248 | 3.00E-52 | 95.98 | GO:0006355 GO:0008219 GO:0009723 GO:0009735 GO:0009753 GO:0009873 GO:0010286 GO:0045893 GO:0051707 | regulation of transcription, DNA-dependent; "cell death"; "response to ethylene stimulus"; "response to cytokinin stimulus"; "response to jasmonic acid stimulus"; "ethylene mediated signaling pathway"; "heat acclimation"; "positive regulation of transcription, DNA-dependent"; "response to other organism" | AP2 domain | Y | Y |
| Glyma16g01980 | AT1G01060.3 \| Symbols: LHY, LHY1 \| Homeodomain-like superfamily protein \| chr1:33992-37061 REVERSE LENGTH=645 | 7.00E-143 | 99.87 | GO:0006355 GO:0007623 GO:0009409 GO:0009639 GO:0009651 GO:0009723 GO:0009733 GO:0009737 GO:0009739 GO:0009751 GO:0009753 GO:0042752 GO:0042754 GO:0043433 GO:0046686 GO:0048574 | regulation of transcription, DNA-dependent; "circadian rhythm"; "response to cold"; "response to red or far red light"; "response to salt stress"; "response to ethylene stimulus"; "response to auxin stimulus"; "response to abscisic acid stimulus"; "response to gibberellin stimulus"; "response to salicylic acid stimulus"; "response to jasmonic acid stimulus"; "regulation of circadian rhythm"; "negative regulation of circadian rhythm"; "negative regulation of sequence-specific DNA binding transcription factor activity"; "response to cadmium ion"; "long-day photoperiodism, flowering" | Myb-like DNA-binding domain | Y | Y |
| Glyma16g34340 | AT1G18330.1 \| Symbols: EPR1, RVE7 \| Homeodomain-like superfamily protein \| chr1:6306196-6307640 REVERSE LENGTH=346 | 6.00E-58 | 59 | GO:0006355 GO:0007623 GO:0009651 GO:0009723 GO:0046686 | regulation of transcription, DNA-dependent; "circadian rhythm"; "response to salt stress"; "response to ethylene stimulus"; "response to cadmium ion" | Myb-like DNA-binding domain | Y | Y |
| Glyma17g06290 | AT4G26150.1 \| Symbols: CGA1, GATA22, GNL \| cytokinin-responsive gata factor 1 \| chr4:13253210-13254659 FORWARD LENGTH=352 | 3.00E-37 | 63.47 | GO:0006355 GO:0007623 GO:0009416 GO:0009735 GO:0009740 GO:0009910 GO:0010187 GO:0010380 GO:0010468 | regulation of transcription, DNA-dependent; "circadian rhythm"; "response to light stimulus"; "response to cytokinin stimulus"; "gibberellic acid mediated signaling pathway"; "negative regulation of flower development"; "negative regulation of seed germination"; "regulation of chlorophyll biosynthetic process"; "regulation of gene expression" | GATA zinc finger | N | N |
| Glyma19g38690 | AT2G35940.3 \| Symbols: BLH1 \| BEL1-like homeodomain 1 \| chr2:15089171-15091699 REVERSE LENGTH=680 | 1.00E-172 | 99.71 | GO:0000303 GO:0006355 GO:0006612 GO:0009610 GO:0009651 GO:0009733 GO:0009737 GO:0009743 GO:0009862 GO:0009873 GO:0009963 GO:0010197 GO:0010201 GO:0010363 GO:0019344 GO:0048513 | response to superoxide; "regulation of transcription, DNA-dependent"; "protein targeting to membrane"; "response to symbiotic fungus"; "response to salt stress"; "response to auxin stimulus"; "response to abscisic acid stimulus"; "response to carbohydrate stimulus"; "systemic acquired resistance, salicylic acid mediated signaling pathway"; "ethylene mediated signaling pathway"; "positive regulation of flavonoid biosynthetic process"; "polar nucleus fusion"; "response to continuous far red light stimulus by the high-irradiance response system"; "regulation of plant-type hypersensitive response"; "cysteine biosynthetic process"; "organ development" | Associated with HOX; "Homeobox do | Y | Y |

| | | | | | | | | |
|---|---|---|---|---|---|---|---|---|
| Glyma20g35270 | AT4G14550.1 \| Symbols: IAA14, SLR \| indole-3-acetic acid inducible 14 \| chr4:8348521-8349923 REVERSE LENGTH=228 | 3.00E-118 | 95.56 | GO:0006355 GO:0009733 GO:0009741 GO:0010102 GO:0010583 GO:0045892 GO:0048527 | regulation of transcription, DNA-dependent; "response to auxin stimulus"; "response to brassinosteroid stimulus"; "lateral root morphogenesis"; "response to cyclopentenone"; "negative regulation of transcription, DNA-dependent"; "lateral root development" | AUX/IAA family | Y | Y |

* 33 of the 35 core TFs belong to one of the soybean root-related transcriptome profiles, with 25 TFs in common.

RM-TF: TFs in soybean root elongation region-specific Microarray gene expression profile under water deficits

RR-TF: TFs in soybean primary root RNA-seq transcriptome profiling during varying water deficit conditions

Y represents being in gene list

N represents not being in gene list

### Table S5. Core TFs involved in multiple signaling pathways

| | | Response to abiotic stress stumili | | | | | | | | | | Response to hormones stumili or being mediated hormon signaling | | | | | | | | Others | | | | In co-function network |
|---|---|---|---|---|---|---|---|---|---|---|---|---|---|---|---|---|---|---|---|---|---|---|---|---|
| No. | Glyma 1.1 ID | water deprivation | desiccation | salt | hyperosmotic salinity | cold | osmotic | superoxide | peroxide | oxidative | heat | ABA | Auxin | BR | CK | ETH | GA | JA | SA | circadian rhythms | light | gravitropism | flavonoid biosynthetic | |
| 1 | Glyma01g12740 | | Y | | | | | | | Y | | Y | | | | | | Y | Y | | | | | |
| 2 | Glyma01g22260 | | | | | | | | | | | | | | | Y | | | | | | | | |
| 3 | Glyma01g40450 | | | | | | | | | | | | | | Y | | Y | | | | | | | Y |
| 4 | Glyma02g03020 | | | Y | | | | | | | | Y | Y | | | | | Y | Y | | | | | |
| 5 | Glyma02g11060 | | | | | | | | | | | | | | | Y | | | | | | | | |
| 6 | Glyma02g35450 | | | Y | | | | Y | Y | | | Y | Y | | | Y | | | Y | | Y | | Y | Y |
| 7 | Glyma02g45150 | | | | | | | | Y | | | | Y | | | | | | Y | Y | Y | Y | | Y |
| 8 | Glyma03g36070 | | | Y | | | | Y | Y | | | Y | Y | | | Y | | | Y | | Y | | Y | Y |
| 9 | Glyma04g04170 | Y | | Y | | | | | | | | Y | | | | | | | | | | | | Y |
| 10 | Glyma04g04490 | | | | | | | | | | | Y | | | | Y | | Y | Y | | | | | |
| 11 | Glyma04g04760 | Y | | Y | Y | Y | Y | | | Y | | Y | Y | | | Y | | Y | | | Y | | | Y |
| 12 | Glyma04g11290 | Y | | Y | | Y | Y | | | | | | | | Y | Y | | | | | Y | | | Y |
| 13 | Glyma04g38560 | Y | | | Y | | Y | | | | | Y | | | | | | Y | | | | | | Y |
| 14 | Glyma05g04950 | | | | | | | | | | | Y | | | | Y | Y | Y | Y | | | | | |
| 15 | Glyma05g32850 | Y | | | Y | | Y | | | | | Y | | | | | | Y | | | | | | Y |
| 16 | Glyma06g04840 | Y | | Y | Y | Y | Y | | | Y | | Y | Y | | | Y | | Y | | | Y | | | Y |
| 17 | Glyma06g11010 | Y | | Y | | Y | Y | | | | | | | | Y | Y | | | | | Y | | | Y |
| 18 | Glyma07g05410 | | | Y | | Y | | | | | | Y | Y | | | Y | Y | Y | Y | Y | Y | | | Y |
| 19 | Glyma07g10311 | | | | | | | | | | | | | Y | | | | | | | | | | Y |
| 20 | Glyma08g10140 | | | Y | Y | | Y | | | | | Y | | | | Y | Y | Y | Y | | Y | | | |
| 21 | Glyma08g22190 | | | | | | | | | | | | Y | Y | | | | | | | | Y | | Y |
| 22 | Glyma08g36720 | | Y | | | | | | | Y | | Y | | | | | | Y | Y | | | | Y | |
| 23 | Glyma08g47520 | | | Y | | | | | | | | Y | | Y | | | | | | | | | Y | Y |
| 24 | Glyma09g04630 | | | | | | | | | | Y | | | | Y | Y | | Y | | | | | | |
| 25 | Glyma09g29800 | | | Y | | | | | | | | | | | | Y | | | | Y | | | | Y |
| 26 | Glyma09g33730 | | Y | | | | | | | Y | | Y | | | | | | Y | Y | | | | Y | |
| 27 | Glyma10g05560 | | | Y | | | | | | | | Y | Y | | | Y | Y | Y | Y | Y | | | | Y |
| 28 | Glyma10g10040 | | | Y | | | | Y | Y | | | Y | Y | | | Y | | | Y | | Y | | Y | Y |
| 29 | Glyma11g33720 | | | Y | Y | | Y | | | | | Y | | | | Y | Y | Y | Y | | | | | |
| 30 | Glyma14g09760 | Y | | Y | Y | Y | Y | | | Y | | Y | Y | | | Y | | Y | | | Y | | | Y |
| 31 | Glyma15g16260 | | | | | | | | | | Y | | | | Y | Y | | Y | | | | | | |
| 32 | Glyma16g01980 | | | Y | | Y | | | | | | Y | Y | | | Y | Y | Y | Y | Y | Y | | | Y |
| 33 | Glyma16g34340 | | | Y | | | | | | | | | | | | Y | | | | Y | | | | Y |
| 34 | Glyma17g16930 | | | | | | | | | | | | | | Y | | | | | | | | | Y |
| 35 | Glyma19g38690 | | | Y | | | | Y | Y | | | Y | Y | | | Y | | | Y | | Y | | Y | Y |
| 36 | Glyma20g35270 | | | | | | | | | | | | Y | Y | | | | | | | | | | Y |
| | | 8 | 3 | 20 | 7 | 7 | 9 | 4 | 5 | 6 | 2 | 22 | 14 | 4 | 6 | 23 | 7 | 19 | 16 | 6 | 13 | 2 | 8 | |
| | Core TFs involved | 27 | | | | | | | | | | 36 | | | | | | | | 20 | | | | 24 |

*Core TFs are involved in multiple hormonal and other signaling pathways based on SoyBase genome annotation . Y represents the core TF involved in the signaling pathway pointed

**Table S6. Core TFs in the co-functional TF–TF network**

| Gene ID | TF Family | Score | Evidences | Betweenness Centrality | Closeness Centrality | Neighborhood Connectivity | Number Of Directed Edges | Radiality | Topological Coefficient |
|---|---|---|---|---|---|---|---|---|---|
| Glyma01g40450 | Homeodomain/HOMEOBOX | 28.14 | AT-CC:0.62 HS-LC:0.38 | 0.007 | 0.474 | 6.714 | 7 | 0.778 | 0.610 |
| Glyma02g35450 | Homeodomain/HOMEOBOX | 25.37 | AT-CC:0.65 HS-LC:0.20 GM-CX:0.15 | 0.047 | 0.514 | 7.500 | 6 | 0.811 | 0.577 |
| Glyma02g45150 | bHLH | 11.65 | AT-CC:0.71 AT-LC:0.29 | 0.000 | 0.375 | 4.000 | 3 | 0.667 | 0.667 |
| Glyma03g36070 | Homeodomain/HOMEOBOX | 22.33 | AT-CC:0.70 HS-LC:0.23 GM-CX:0.07 | 0.047 | 0.514 | 7.500 | 6 | 0.811 | 0.577 |
| Glyma04g04170 | BZIP | 3.51 | AT-CC:1.00 | 0.000 | 0.310 | 4.000 | 1 | 0.556 | 0.000 |
| Glyma04g04760 | C2H2 (Zn) | 4.98 | GM-CX:1.00 | 0.000 | 1.000 | 1.000 | 1 | 1.000 | 0.000 |
| Glyma04g11290 | AP2-EREBP | 9.23 | AT-CC:0.68 GM-CX:0.32 | 0.000 | 0.375 | 4.000 | 3 | 0.667 | 0.667 |
| Glyma04g38560 | NAC | 3.23 | GM-CX:1.00 | 0.000 | 1.000 | 1.000 | 1 | 1.000 | 0.000 |
| Glyma05g32850 | NAC | 3.23 | GM-CX:1.00 | 0.000 | 1.000 | 1.000 | 1 | 1.000 | 0.000 |
| Glyma06g04840 | C2H2 (Zn) | 2.39 | GM-CX:1.00 | 0.000 | 1.000 | 1.000 | 1 | 1.000 | 0.000 |
| Glyma06g11010 | AP2-EREBP | 8.9 | AT-CC:0.67 GM-CX:0.33 | 0.000 | 0.375 | 4.000 | 3 | 0.667 | 0.667 |
| Glyma07g05410 | MYB/HD-like | 5.35 | GM-CX:0.56 AT-CX:0.44 | 0.111 | 0.310 | 4.000 | 1 | 0.556 | 0.000 |
| Glyma07g10311 | bHLH | 19.28 | AT-CC:1.00 | 0.302 | 0.545 | 5.333 | 6 | 0.833 | 0.369 |
| Glyma08g22190 | AUX-IAA-ARF | 3.19 | AT-HT:1.00 | 0.000 | 0.333 | 8.000 | 1 | 0.600 | 0.000 |
| Glyma08g47520 | NAC | 25.1 | AT-CC:1.00 | 0.146 | 0.529 | 6.500 | 8 | 0.822 | 0.490 |
| Glyma09g29800 | MYB/HD-like | 25.03 | AT-CC:0.87 GM-CX:0.13 | 0.209 | 0.621 | 5.250 | 8 | 0.878 | 0.320 |
| Glyma10g05560 | MYB/HD-like | 13.45 | AT-CC:0.71 AT-CX:0.16 GM-CX:0.12 | 0.307 | 0.439 | 4.250 | 4 | 0.744 | 0.469 |
| Glyma10g10040 | Homeodomain/HOMEOBOX | 24.46 | AT-CC:0.70 HS-LC:0.21 GM-CX:0.09 | 0.047 | 0.514 | 7.500 | 6 | 0.811 | 0.577 |
| Glyma14g09760 | C2H2 (Zn) | 2.59 | GM-CX:1.00 | 0.000 | 1.000 | 1.000 | 1 | 1.000 | 0.000 |
| Glyma16g01980 | MYB/HD-like | 3.13 | GM-CX:1.00 | 0.000 | 0.391 | 8.000 | 1 | 0.689 | 0.000 |
| Glyma16g34340 | MYB/HD-like | 24.26 | AT-CC:0.93 GM-CX:0.07 | 0.209 | 0.600 | 6.000 | 7 | 0.867 | 0.353 |
| Glyma17g16930 | Homeodomain/HOMEOBOX | 25.96 | AT-CC:0.60 HS-LC:0.40 | 0.004 | 0.474 | 6.714 | 7 | 0.778 | 0.610 |
| Glyma19g38690 | Homeodomain/HOMEOBOX | 22.35 | AT-CC:0.70 HS-LC:0.23 GM-CX:0.07 | 0.047 | 0.514 | 7.500 | 6 | 0.811 | 0.577 |
| Glyma20g35270 | AUX-IAA-ARF | 22.24 | AT-CC:0.79 AT-HT:0.13 GM-CX:0.08 | 0.115 | 0.486 | 5.875 | 8 | 0.789 | 0.575 |

*Each core TF was scored as the sum of LLS by summing network edge weights connecting two TFs with network evidence, such as co-expression, literature mining, physical protein interactions and other data from orthologs

AT-CX = mRNA co-expression between Arabidopsis genes

AT-LC = Literature-curated Arabidopsis protein interactions

AT-CC = By co-citation of *Arabidopsis thaliana* orthologs in Pubmed articles

AT-HT = By high-throughput *Arabidopsis thaliana* orthologous PPI

GM-CX = By co-expression of *Glycine max* (soybean) genes

HS-LC = By literature curated Homo sapiens (human) orthologous PPIs

### Table S7. Consensus sequences of selected motifs from Group I

| Motif name | Sequence | Conserved motif related |
|---|---|---|
| dACAC | ACACACACAC | RDmotif 11, 16, 20, 38 |
| dASE | GAGGAAG | RDmotif 10, 48 |
| dGA | GGGAGG | RDmotif 10, 46, 48 |
| dGAGA | GAGAGAGAGA | RDmotif 8 |
| dGT | TGGTGG | RDmotif 30, 61 |
| dTG | TGGGGT | RDmotif 29, 31 |
| mdASE | acagaag-gagacca | dASE mutant |
| mdGAGT | aaatggg-gggtaaa | dGA and dGT mutant |
| mdTG | caaggt-tggaac | dTG mutant |

### Table S8. List of 214 drought-related TF genes

| Gene ID |
|---|
| Glyma01g02350.1 |
| Glyma01g02830.1 |
| Glyma01g06170.1 |
| Glyma01g11390.1 |
| Glyma01g17850.1 |
| Glyma01g28820.1 |
| Glyma01g30610.1 |
| Glyma01g39360.1 |
| Glyma01g44130.1 |
| Glyma02g00820.1 |
| Glyma02g01960.1 |
| Glyma02g04550.1 |
| Glyma02g04710.3 |
| Glyma02g05070.1 |
| Glyma02g08110.1 |
| Glyma02g10530.1 |
| Glyma02g10940.1 |
| Glyma02g14880.1 |
| Glyma02g15520.1 |
| Glyma02g15520.2 |
| Glyma02g15780.1 |
| Glyma02g26480.1 |
| Glyma02g41180.1 |
| Glyma02g44260.1 |
| Glyma02g46970.1 |
| Glyma02g47380.1 |
| Glyma03g04000.1 |
| Glyma03g08270.X |
| Glyma03g25280.1 |
| Glyma03g25280.2 |
| Glyma03g29750.1 |
| Glyma03g35010.X |
| Glyma03g36070.1 |
| Glyma03g38040.1 |
| Glyma03g39990.2 |
| Glyma03g40610.1 |
| Glyma03g40730.1 |
| Glyma04g04190.X |
| Glyma04g04310.1 |
| Glyma04g04490.1 |
| Glyma04g05290.1 |
| Glyma04g05500.X |
| Glyma04g08990.1 |
| Glyma04g34080.1 |

| |
|---|
| Glyma04g34720.1 |
| Glyma04g36110.1 |
| Glyma04g40960.1 |
| Glyma04g43180.1 |
| Glyma04g43640.1 |
| Glyma05g03650.1 |
| Glyma05g28130.X |
| Glyma05g29590.1 |
| Glyma05g31400.1 |
| Glyma05g35060.1 |
| Glyma05g35650.1 |
| Glyma05g36970.1 |
| Glyma05g38290.1 |
| Glyma06g04350.X |
| Glyma06g04353.1 |
| Glyma06g05170.X |
| Glyma06g06780.1 |
| Glyma06g08990.1 |
| Glyma06g12070.1 |
| Glyma06g14720.1 |
| Glyma06g15820.1 |
| Glyma06g20230.1 |
| Glyma06g20800.1 |
| Glyma06g21020.1 |
| Glyma06g38410.1 |
| Glyma07g02180.1 |
| Glyma07g02320.1 |
| Glyma07g05740.X |
| Glyma07g06090.1 |
| Glyma07g09520.1 |
| Glyma07g10180.1 |
| Glyma07g15850.X |
| Glyma07g30860.1 |
| Glyma07g36670.1 |
| Glyma08g02020.1 |
| Glyma08g03540.1 |
| Glyma08g06440.1 |
| Glyma08g09190.1 |
| Glyma08G14600.1 |
| Glyma08g17140.1 |
| Glyma08g18470.1 |
| Glyma08g20170.1 |
| Glyma08g24420.1 |
| Glyma08g38190.1 |
| Glyma08g38470.1 |
| Glyma08g42300.1 |
| Glyma08g47520.1 |

| |
|---|
| Glyma08g48230.1 |
| Glyma09g01650.1 |
| Glyma09g04630.1 |
| Glyma09g07960.1 |
| Glyma09g25590.1 |
| Glyma09g33241.1 |
| Glyma09g33630.3 |
| Glyma09g37050.1 |
| Glyma09g37780.1 |
| Glyma09g41050.1 |
| Glyma10g03530.1 |
| Glyma10g04210.1 |
| Glyma10g04350.1 |
| Glyma10g23440.1 |
| Glyma10g28290.1 |
| Glyma10g30650.1 |
| Glyma10g33060.1 |
| Glyma10g34760.X |
| Glyma10g34780.1 |
| Glyma11g01100.1 |
| Glyma11g02960.1 |
| Glyma11g03850.1 |
| Glyma11g04840.1 |
| Glyma11g05810.X |
| Glyma11g11640.1 |
| Glyma11g13960.X |
| Glyma12g01960.1 |
| Glyma12g02540.1 |
| Glyma12g07110.1 |
| Glyma12g22880.1 |
| Glyma12g29010.1 |
| Glyma12g30990.1 |
| Glyma12g33320.1 |
| Glyma12g33600.1 |
| glyma12g33990.1 |
| Glyma12g35000.1 |
| Glyma12g36540.1 |
| Glyma13g01200.1 |
| Glyma13g01290.1 |
| Glyma13g04630.1 |
| Glyma13g05380.1 |
| Glyma13g11940.1 |
| Glyma13g12200.1 |
| Glyma13g16770.1 |
| Glyma13g16770.X |
| Glyma13g18410.1 |
| Glyma13g30720.1 |

| |
|---|
| Glyma13g31010.1 |
| Glyma13g32090.1 |
| Glyma13g32650.1 |
| Glyma13g33290.1 |
| Glyma13g35550.1 |
| Glyma13g35810.1 |
| Glyma13g36540.1 |
| Glyma13g38080.1 |
| Glyma14g03300.1 |
| Glyma14g03510.1 |
| Glyma14g24220.1 |
| Glyma14g36010.1 |
| Glyma14g37050.1 |
| Glyma14g39530.1 |
| Glyma14g40450.1 |
| Glyma15g00660.1 |
| Glyma15g01550.1 |
| Glyma15g01550.4 |
| Glyma15g01550.5 |
| Glyma15g01550.6 |
| Glyma15g04620.1 |
| Glyma15g07010.1 |
| Glyma15g08360.1 |
| Glyma15g19460.1 |
| Glyma16g02390.1 |
| Glyma16g02570.1 |
| Glyma16g02960.1 |
| Glyma16g05480.1 |
| Glyma17g03570.X |
| Glyma17g06450.1 |
| Glyma17g07330.1 |
| Glyma17g07860.1 |
| Glyma17g07860.X |
| Glyma17g10490.1 |
| Glyma17g15480.1 |
| Glyma17g16360.1 |
| Glyma17g16930.1 |
| Glyma17g16930.X |
| Glyma17g18110.1 |
| Glyma17g19830.1 |
| Glyma17g20070.1 |
| Glyma17g34010.1 |
| Glyma17g35430.1 |
| Glyma17g36370.1 |
| Glyma17g37430.1 |
| Glyma17g37710.1 |
| Glyma18g02430.1 |

| |
|---|
| Glyma18g04580.1 |
| Glyma18g05050.1 |
| Glyma18g05050.X |
| Glyma18g20990.1 |
| Glyma18g44560.1 |
| Glyma18g48730.1 |
| Glyma18g49360.1 |
| Glyma18g51250.1 |
| Glyma19g27240.X |
| Glyma19g35180.1 |
| Glyma19g35180.2 |
| Glyma19g35180.3 |
| Glyma19g36570.1 |
| Glyma19g38800.1 |
| Glyma19g40470.1 |
| Glyma19g40980.1 |
| Glyma19g40980.X |
| Glyma19g41250.1 |
| Glyma19g41610.1 |
| Glyma19g41610.2 |
| Glyma19g43580.1 |
| Glyma19g44190.1 |
| Glyma19g44610.1 |
| Glyma20g00240.1 |
| Glyma20g01690.1 |
| Glyma20g16910.1 |
| Glyma20g24690.1 |
| Glyma20g34570.1 |
| Glyma20g39220.1 |

X represents new splicing varient

**Table S9 List of 13 TF hits in the TF regulatory network**

| Gene ID | TF family | Top TAIR10 Blastp Hit | Gene Ontology Biological Process IDs | Gene Ontology Biological Process Descriptions |
|---|---|---|---|---|
| Glyma02g01960 | AP2 | AT3G16770.1 \| Symbols: RAP2.3, ATEBP, ERF72, EBP \| ethylene-responsive element binding protein \| chr3:5705784-5706768 FORWARD LENGTH=248 | GO:0006355 GO:0008219 GO:0009723 GO:0009735 GO:0009753 GO:0009873 GO:0010286 GO:0045893 GO:0051707 | regulation of transcription, DNA-dependent; "cell death"; "response to ethylene stimulus"; "response to cytokinin stimulus"; "response to jasmonic acid stimulus"; "ethylene mediated signaling pathway"; "heat acclimation"; "positive regulation of transcription, DNA-dependent"; "response to other organism" |
| Glyma06g04353 | bZIP | AT1G45249.1 \| Symbols: ABF2, AREB1, ATAREB1 \| abscisic acid responsive elements-binding factor 2 \| chr1:17165420-17167415 REVERSE LENGTH=416 | GO:0006355 GO:0009414 GO:0009651 GO:0009737 GO:0009738 GO:0010255 GO:0045893 | regulation of transcription, DNA-dependent; "response to water deprivation"; "response to salt stress"; "response to abscisic acid stimulus"; "abscisic acid mediated signaling pathway"; "glucose mediated signaling pathway"; "positive regulation of transcription, DNA-dependent" |
| Glyma06g21020 | NAC | AT5G61430.1 \| Symbols: ANAC100, ATNAC5, NAC100 \| NAC domain containing protein 100 \| chr5:24701328-24702553 REVERSE LENGTH=336 | GO:0006355 GO:0007275 | regulation of transcription, DNA-dependent; "multicellular organismal development" |
| Glyma08g18470 | NAC | AT1G56010.2 \| Symbols: NAC1, ANAC022 \| NAC domain containing protein 1 \| chr1:20946852-20949144 REVERSE LENGTH=324 | GO:0006355 GO:0007275 GO:0009734 GO:0010072 GO:0048527 | regulation of transcription, DNA-dependent; "multicellular organismal development"; "auxin mediated signaling pathway"; "primary shoot apical meristem specification"; "lateral root development" |
| Glyma09g01650 | CBF/NF-Y | AT4G14540.1 \| Symbols: NF-YB3 \| nuclear factor Y, subunit B3 \| chr4:8344663-8345148 FORWARD LENGTH=161 | GO:0006355 | regulation of transcription, DNA-dependent |
| Glyma09g33241 | AP2 | AT5G10510.2 \| Symbols: AIL6 \| AINTEGUMENTA-like 6 \| chr5:3315991-3320008 FORWARD LENGTH=581 | GO:0006355 GO:0009855 GO:0009887 GO:0009908 GO:0009944 GO:0010014 GO:0010075 GO:0010080 GO:0010089 GO:0010492 GO:0035265 GO:0044036 GO:0048364 GO:0060771 | regulation of transcription, DNA-dependent; "determination of bilateral symmetry"; "organ morphogenesis"; "flower development"; "polarity specification of adaxial/abaxial axis"; "meristem initiation"; "regulation of meristem growth"; "regulation of floral meristem growth"; "xylem development"; "maintenance of shoot apical meristem identity"; "organ growth"; "cell wall macromolecule metabolic process"; "root development"; "phyllotactic patterning" |
| Glyma09g37050 | NAC | AT5G61430.1 \| Symbols: ANAC100, ATNAC5, NAC100 \| NAC domain containing protein 100 \| chr5:24701328-24702553 REVERSE LENGTH=336 | GO:0006355 GO:0007275 | regulation of transcription, DNA-dependent; "multicellular organismal development" |
| Glyma10g28290 | bHLH | AT1G09530.2 \| Symbols: PIF3, POC1, PAP3 \| phytochrome interacting factor 3 \| chr1:3077216-3079367 FORWARD LENGTH=524 | GO:0006355 GO:0007165 GO:0007623 GO:0009630 GO:0009639 GO:0009704 GO:0009740 GO:0010017 GO:0031539 | regulation of transcription, DNA-dependent; "signal transduction"; "circadian rhythm"; "gravitropism"; "response to red or far red light"; "de-etiolation"; "gibberellic acid mediated signaling pathway"; "red or far-red light signaling pathway"; "positive regulation of anthocyanin metabolic process" |
| Glyma11g03850 | MYB-HD-like | AT5G47370.1 \| Symbols: HAT2 \| Homeobox-leucine zipper protein 4 (HB-4) / HD-ZIP protein \| chr5:19216482-19217647 REVERSE LENGTH=283 | GO:0006351 GO:0006355 GO:0009641 GO:0009733 GO:0009734 GO:0009826 GO:0045892 | transcription, DNA-dependent; "regulation of transcription, DNA-dependent"; "shade avoidance"; "response to auxin stimulus"; "auxin mediated signaling pathway"; "unidimensional cell growth"; "negative regulation of transcription, DNA-dependent" |
| Glyma13g18410 | AP2 | AT3G23240.1 \| Symbols: ERF1, ATERF1 \| ethylene response factor 1 \| chr3:8295705-8296361 FORWARD LENGTH=218 | GO:0006355 GO:0006952 GO:0009867 GO:0009873 | regulation of transcription, DNA-dependent; "defense response"; "jasmonic acid mediated signaling pathway"; "ethylene mediated signaling pathway" |
| Glyma17g07860 | AP2 | AT1G16060.1 \| Symbols: ADAP \| ARIA-interacting double AP2 domain protein \| chr1:5508563-5511609 FORWARD LENGTH=345 | GO:0006355 GO:0009414 GO:0009651 GO:0009737 GO:0009887 GO:0010187 GO:0040008 | regulation of transcription, DNA-dependent; "response to water deprivation"; "response to salt stress"; "response to abscisic acid stimulus"; "organ morphogenesis"; "negative regulation of seed germination"; "regulation of growth" |

| | | | | |
|---|---|---|---|---|
| Glyma20g34570 | AP2 | AT3G23240.1 \| Symbols: ERF1, ATERF1 \| ethylene response factor 1 \| chr3:8295705-8296361 FORWARD LENGTH=218 | GO:0006355 GO:0006952 GO:0009867 GO:0009873 | regulation of transcription, DNA-dependent; "defense response"; "jasmonic acid mediated signaling pathway"; "ethylene mediated signaling pathway" |

*13 different TFs were identified for the motif Gbox-I-gc group using Y1H screen.

**Table S10. Core TFs in core sensory TF network**

| Gene ID | TF Family | Score | Evidences | BetweennessCentrality | ClosenessCentrality | Degree | NeighborhoodConnectivity | Radiality | TopologicalCoefficient |
|---|---|---|---|---|---|---|---|---|---|
| ARG | BZIP | 9.26 | AT-CC:0.64 GM-CX:0.36; Y1H | 0.05110849 | 0.44 | 4 | 5.25 | 0.78787879 | 0.40384615 |
| BRG | bHLH | 19.28 | AT-CC:1.00; Y1H | 0.45029516 | 0.57894737 | 10 | 4.4 | 0.87878788 | 0.24705882 |
| Glyma01g40450 | Homeodomain/HOMEOBOX | 30.86 | AT-CC:0.58 HS-LC:0.36 AT-HT:0.07 | 0.01731602 | 0.44 | 8 | 6.875 | 0.78787879 | 0.57291667 |
| Glyma02g35450 | Homeodomain/HOMEOBOX | 25.37 | AT-CC:0.65 HS-LC:0.20 GM-CX:0.15 | 0.0289125 | 0.46808511 | 7 | 7.57142857 | 0.81060606 | 0.58241758 |
| Glyma02g45150 | bHLH | 11.67 | AT-CC:0.70 AT-LC:0.30 | 0.004329 | 0.39285714 | 4 | 4.5 | 0.74242424 | 0.45 |
| Glyma03g36070 | Homeodomain/HOMEOBOX | 22.33 | AT-CC:0.70 HS-LC:0.23 GM-CX:0.07 | 0.0289125 | 0.46808511 | 7 | 7.57142857 | 0.81060606 | 0.58241758 |
| Glyma04g04170 | BZIP | 10.1 | AT-CC:0.66 GM-CX:0.34 | 0.00649351 | 0.36065574 | 3 | 4 | 0.70454545 | 0.57142857 |
| Glyma04g11290 | AP2-EREBP | 9.23 | AT-CC:0.68 GM-CX:0.32 | 0 | 0.38596491 | 3 | 5.66666667 | 0.73484848 | 0.56666667 |
| Glyma06g11010 | AP2-EREBP | 8.9 | AT-CC:0.67 GM-CX:0.33 | 0 | 0.38596491 | 3 | 5.66666667 | 0.73484848 | 0.56666667 |
| Glyma07g05410 | MYB/HD-like | 5.35 | GM-CX:0.56 AT-CX:0.44 | 0.09090909 | 0.31884058 | 2 | 3 | 0.64393939 | 0.5 |
| Glyma08g22190 | AUX-IAA-ARF | 3.19 | AT-HT:1.00 | 0 | 0.30985915 | 1 | 8 | 0.62878788 | 0 |
| Glyma08g47520 | NAC | 25.1 | AT-CC:1.00 | 0.13506494 | 0.51162791 | 8 | 7.75 | 0.84090909 | 0.44852941 |
| Glyma09g29800 | MYB/HD-like | 25.03 | AT-CC:0.87 GM-CX:0.13 | 0.14327693 | 0.56410256 | 7 | 7.14285714 | 0.87121212 | 0.35714286 |
| Glyma10g05560 | MYB/HD-like | 16.51 | AT-CC:0.76 AT-CX:0.14 GM-CX:0.10 | 0.22308802 | 0.44 | 5 | 4.6 | 0.78787879 | 0.4 |
| Glyma10g10040 | Homeodomain/HOMEOBOX | 24.46 | AT-CC:0.70 HS-LC:0.21 GM-CX:0.09 | 0.0289125 | 0.46808511 | 7 | 7.57142857 | 0.81060606 | 0.58241758 |
| Glyma16g01980 | MYB/HD-like | 3.13 | GM-CX:1.00 | 0 | 0.24444444 | 1 | 2 | 0.48484848 | 0 |
| Glyma16g34340 | MYB/HD-like | 24.26 | AT-CC:0.93 GM-CX:0.07 | 0.14327693 | 0.56410256 | 7 | 7.14285714 | 0.87121212 | 0.35714286 |
| Glyma17g16930 | Homeodomain/HOMEOBOX | 28.3 | AT-CC:0.56 HS-LC:0.38 AT-HT:0.06 | 0.01731602 | 0.44 | 8 | 6.875 | 0.78787879 | 0.57291667 |
| Glyma19g38690 | Homeodomain/HOMEOBOX | 22.35 | AT-CC:0.70 HS-LC:0.23 GM-CX:0.07 | 0.0289125 | 0.46808511 | 7 | 7.57142857 | 0.81060606 | 0.58241758 |
| Glyma20g35270 | AUX-IAA-ARF | 22.24 | AT-CC:0.79 AT-HT:0.13 GM-CX:0.08 | 0.09379509 | 0.44 | 8 | 6.625 | 0.78787879 | 0.59090909 |
| HRG | Homeodomain/HOMEOBOX | 5.07 | AT-HT:1.00; Y1H | 0.03174603 | 0.44897959 | 3 | 8.66666667 | 0.79545455 | 0.52083333 |
| NRG | NF-YB | 6 | AT-CC:1.00; Y1H | 0.01746032 | 0.41509434 | 3 | 5.66666667 | 0.76515152 | 0.47222222 |
| PRG | bHLH | 3.56 | AT-LC:1.00; Y1H | 0 | 0.37931034 | 2 | 7 | 0.72727273 | 0.7 |

*List of core TFs in a core TF sensory network with possible mutual regulation among core TFs by integrating co-functional network evidence and TF-DNA interactions.

AT-CX = mRNA co-expression between Arabidopsis genes

AT-LC = Literature-curated Arabidopsis protein interactions

AT-CC = By co-citation of *Arabidopsis thaliana* orthologs in Pubmed articles

AT-HT = By high-throughput *Arabidopsis thaliana* orthologous PPI

GM-CX = By co-expression of *Glycine max* (soybean) genes

HS-LC = By literature curated Homo sapiens (human) orthologous PPIs

Y1H = TF-DNA interaction by Y1H screen

**Table S11. ANOVA for soybean primary root growth of *BRG-ox, ARG-ox, NRG-ox, HRG-ox* and *PRG-ox* plants in response to 50 µM ABA, PEG (-1.2 Mpa), 0.5 nM BL, 10 µM PCZ, or 50 µM NAA treatment (also see Figure S5G).**

| | Df | Sum Sq | Mean Sq | F value | Pr(>F) |
|---|---|---|---|---|---|
| Genotype | 5 | 36623 | 7325 | 374.6 | <2e-16 |
| Treatment | 5 | 428636 | 85727 | 4384 | <2e-16 |
| Genotype X Treatment | 25 | 81355 | 3254 | 166.4 | <2e-16 |
| Residuals | 396 | 7743 | 20 | | |

**Table S12. ANOVA for soybean primary root growth of *BRG-ox*, *brg-si*, *BRG-cres*, *ARG-ox*, *arg-si*, *ARG-cres* and *arg-brg-si* in response to 10 µM, 50 µM, 100 µM ABA, PEG (-1.7 Mpa) or PCZ-BL (also see Figure 9H).**

| | Df | Sum Sq | Mean Sq | F value | Pr(>F) |
|---|---|---|---|---|---|
| Genotype | 7 | 80427 | 11490 | 528.93 | <2e-16 |
| Treatment | 5 | 514951 | 102990 | 4741.2 | <2e-16 |
| Genotype X Treatment | 35 | 32175 | 919 | 42.32 | <2e-16 |
| Residuals | 672 | 14597 | 22 | | |

**Table S13. ANOVA for Arabidopsis WT, *ARG-ox* and *BRG-ox* primary root length in response to 10 µM ABA, 50 µM ABA or PEG (-1.7 Mpa) under continuous light and in the dark (also see Figure 10E).**

Under continuous light

| | Df | Sum Sq | Mean Sq | F value | Pr(>F) |
|---|---|---|---|---|---|
| Genotype | 2 | 732 | 366 | 40.78 | 1.26E-15 |
| Treatment | 3 | 294825 | 98275 | 10942.51 | <2e-16 |
| Genotype X Treatment | 6 | 2883 | 480 | 53.5 | <2e-16 |
| Residuals | 204 | 1832 | 9 | | |

In the dark

| | Df | Sum Sq | Mean Sq | F value | Pr(>F) |
|---|---|---|---|---|---|
| Genotype | 2 | 2777 | 1389 | 82.8 | <2e-16 |
| Treatment | 3 | 124767 | 41589 | 2479.89 | <2e-16 |
| Genotype X Treatment | 6 | 3991 | 665 | 39.67 | <2e-16 |
| Residuals | 204 | 3421 | 17 | | |

**Table S14. List of primers used in this study**

| Primer Name | Sequence |
|---|---|
| **Conserved motifs clone** | |
| mGbox-fw | CACCTAATCAATTGATACTAATCAATTGATACTAATCAATTGATACTAATCAATTGATAA |
| mGbox-rv | TTATCAATTGATTAGTATCAATTGATTAGTATCAATTGATTAGTATCAATTGATTAGGTG |
| I-Gbox-gc-fw | CACCTAGCCACGTGGCACTAGCCACGTGGCACTAGCCACGTGGCACTAGCCACGTGGCAA |
| I-Gbox-gc-rv | TTGCCACGTGGCTAGTGCCACGTGGCTAGTGCCACGTGGCTAGTGCCACGTGGCTAGGTG |
| II-Gbox-fw | CACCTAAGCACGTGCAACTAAGCACGTGCAACTAAGCACGTGCAACTAAGCACGTGCAAA |
| II-Gbox-rv | TTTGCACGTGCTTAGTTGCACGTGCTTAGTTGCACGTGCTTAGTTGCACGTGCTTAGGTG |
| hB-Gbox-fw | CACCTAGGCACGTGGTACTAGGCACGTGGTACTAGGCACGTGGTACTAGGCACGTGGTAA |
| hB-Gbox-rv | TTACCACGTGCCTAGTACCACGTGCCTAGTACCACGTGCCTAGTACCACGTGCCTAGGTG |
| I-Gbox-tc-fw | CACCTATCCACGTGGCACTATCCACGTGGCACTATCCACGTGGCACTATCCACGTGGCAA |
| I-Gbox-tc-rv | TTGCCACGTGGATAGTGCCACGTGGATAGTGCCACGTGGATAGTGCCACGTGGATAGGTG |
| I-Gbox-ac-fw | CACCTAACCACGTGGCACTAACCACGTGGCACTAACCACGTGGCACTAACCACGTGGCAA |
| I-Gbox-ac-rv | TTGCCACGTGGTTAGTGCCACGTGGTTAGTGCCACGTGGTTAGTGCCACGTGGTTAGGTG |
| dGAGA-fw | CACCGAGAGAGAGAGAGAGAGAGAGAGAGAGAGAGAGAGAGAGAGAGAGAGAGAGAGAGAGA |
| dGAGA-rv | TCTCTCTCTCTCTCTCTCTCTCTCTCTCTCTCTCTCTCTCTCTCTCTCTCTCTCTCGGTG |
| dACAC-fw | CACCACACACACACACACACACACACACACACACACACACACACACACACACACACACACA |
| dACAC-rv | TGTGTGTGTGTGTGTGTGTGTGTGTGTGTGTGTGTGTGTGTGTGTGTGTGTGTGTGGTG |
| dGA BOX-fw | CACCTGGGAGGAGTGGGAGGTGTGGGAGGTGAGGGAGGTATGGGAGGAT |
| dGA BOX-rv | ATCCTCCCATACCTCCCTCACCTCCCACACCTCCCACTCCTCCCAGGTG |
| dGT BOX-fw | CACCTGGTGGGAGTGGTGGGTGTGGTGGGTGAGGTGGGATAGGTGGGAT |
| dGT BOX-rv | ATCCCACCTATCCCACCTCACCCACCACACCCACCACTCCCACCAGGTG |
| mGAGT fw | CACCTAAATGGGAGTAAATGGGTATGGGTAAATGAGGGTAAAATAGGGTAAAAT |
| mGAGT rv | ATTTTACCCTATTTTACCCTCATTTACCCATACCCATTTACTCCCATTTAGGTG |
| dASE-box fw | CACCTGAGGAAGAGTGAGGAAGTATGAGGAAGTGAGAGGAAGATAGAGGAAGAT |
| dASE-box rv | ATCTTCCTCTATCTTCCTCTCACTTCCTCATACTTCCTCACTCTTCCTCAGGTG |
| mASEfw | CACCTACAGAAGAGTACAGAAGTATGAGACCATGAGAGACCAATAGAGACCAAT |
| mASE rv | ATTGGTCTCTATTGGTCTCTCATGGTCTCATACTTCTGTACTCTTCTGTAGGTG |
| dTG-fw | CACCATGGGGTTGATGGGGTACTTGGGGTCTATGGGGTTCATGGGGTAA |
| dTG-rv | TTACCCCATGAACCCCATAGACCCCAAGTACCCCATCAACCCCATGGTG |
| mTG-fw | CACCACAAGGTTGACAAGGTACTTGGAACCTATGGAACTCATGGAACAA |
| mTG-rv | TTGTTCCATGAGTTCCATAGGTTCCAAGTACCTTGTCAACCTTGTGGTG |
| **Dual-Luc vector construction** | |
| Ren-sacI-Rv | AATTCAGAGCTCCCGATCTAGTAACATAGATGACACC |
| Ren-sacI-Fw | TATTCTGAGCTCGTGCCTGCAGGTCAACATGGTGGAGC |
| Mini35S-HindIII-Rv | TTCCAGGAAGCTTTCCTCTCCAAATGAAATGAACTTCCTTATATAGAGGAAGGGTCTTGCGAAGGA TAGTGGGGGAATTATCGAACCACTTTGTACAAG |
| Mini35S-HindIII-fw | AAGCTTTCCTCTCCAAATGAAATGAACTTC |
| **ChIP-PCR** | |
| Chip-BRG-R1fw | CAAAAACTACAGAAGCTTGAGGTG |
| Chip-BRG-R1rv | TTGGTGAATAGGACCATTCTGAGG |
| Chip-BRGpr5-fw | ATTGAGAATTAGACGCATTTGCG |
| Chip-BRGpr5-rv | ATACTTGTACTTCTTGTTACAAAG |
| Chip-BRGcs-fw | TTGCAGACCATGGGTATGGCAG |

| | |
|---|---|
| Chip-BRGcs-rv | TCTACTTGATGCTGCAAGGACTGC |
| Chip-BRG-R2fw | AAACTACAGAAGCTTGAGGTGCC |
| Chip-BRG-R2rv | CTATTTATACTGTGGAATTATGGC |
| **CDS Clone primers** | |
| Glyma07g10311_attB1 | GGGGACAAGTTTGTACAAAAAAGCAGGCTTCATGGCTGAATTCACAGAAAATATGC |
| Glyma07g10311_attB2 | GGGGACCACTTTGTACAAGAAAGCTGGGTATCAAAGGGGCCATGCTGGTTGGAAG |
| Glyma06g04353-attB1 | GGGGACAAGTTTGTACAAAAAAGCAGGCTTCATGAGGTTTGGAAGGACATTTTCG |
| Glyma06g04353-attB2ws | GGGGACCACTTTGTACAAGAAAGCTGGGTACTACCATGGACCAGTTTGTGTTCG |
| Glyma06g04353-attB2wos | GGGGACCACTTTGTACAAGAAAGCTGGGTACCATGGACCAGTTTGTGTTCGTCTTAG |
| Glyma11g03850-attB1 | GGGGACAAGTTTGTACAAAAAAGCAGGCTTCATGACGGTTGAAAAGGAAGATTTGGG |
| Glyma11g03850-attB2 | GGGGACCACTTTGTACAAGAAAGCTGGGTATCAAGATCTGCGACGGAGGGCATCGAAG |
| Glyma09g01650-attB1 | GGGGACAAGTTTGTACAAAAAAGCAGGCTTCATGGCTGAGTCCGACAACGAGTCAG |
| Glyma09g01650-attB2 | GGGGACCACTTTGTACAAGAAAGCTGGGTACTATCTAGTTCTTGCAGAAGATGCTG |
| Glyma10g28290-attB1 | GGGGACAAGTTTGTACAAAAAAGCAGGCTTCATGTCGGTGCCATCCAGTGAAGTGG |
| Glyma10g28290-attB2 | GGGGACCACTTTGTACAAGAAAGCTGGGTATTATCTGCCAGGTCCAGGATTAACAG |
| **Q-PCR primers** | |
| GmACT11-F (Glyma18g52780-F) | ATCTTGACTGAGCGTGGTTATTCC |
| GmACT11-R (Glyma18g52780-R) | GCTGGTCCTGGCTGTCTCC |
| Glyma09g01650-Fw | TGTGGCATATGTGGCATACACAG |
| Glyma09g01650-Rv | AGCCATAGTCCTGCCCTTACAC |
| Glyma10g28290-Fw | AGATGCCTACCCAGTGTGAAGC |
| Glyma10g28290-Rv | GCCAGGTCCAGGATTAACAGATCC |
| Glyma10g33060-Fw | ACTCACCCTCTTCCGGTTTCTG |
| Glyma10g33060-Rv | GGCGATCATGCCGTATAGAAGC |
| Glyma11g03850-Fw | GAACGACCCTTTCACTTCTTCAGC |
| Glyma11g03850-Rv | ACCTGCTTCCTCTTCGCAATCG |
| Glyma20g34570-Fw | ACTCACCCTCTTCCGATTTCTCC |
| Glyma20g34570-Rv | AAGGTACCCTTCCCACGAGAAC |
| Glyma02g35450-fw | AGAGAGGATTGCCTGAGAGATCGG |
| Glyma02g35450-rv | TTGTCCGAGTCCTTGGGATAAGGG |
| Glyma02g45150-fw | TGGTTCCCAAGCAGTGCAACAC |
| Glyma02g45150-rv | AAGGTGGAAGAACCCGCTTTGC |
| Glyma04g04170-fw | TAATGGCAGTCTGCGAGGAAGG |
| Glyma04g04170-rv | TCCATGGTATAAGCCTGTTTGCG |
| Glyma06g04350-fw | GACGAGTTGCTGAAGAACATTTGG |
| Glyma06g04350-rv | AAACCTCATCCACGGTCTTCCC |
| Glyma06g11010-fw | CCGCTTTCGGATCTGACGTTTG |
| Glyma06g11010-rv | ATCCGAATCACCTTCCCACTGAG |
| SC-Glyma04g38050-F | CGTCCAGAATTGCTCAACAG |
| SC-Glyma04g38050-R | TGGGGTTATAGCCTTGTTGG |
| SC-Glyma14g20450-F | AATGCCGAAAGCCATTACAG |
| SC-Glyma14g20450-R | GCTTTGTTTTCCCTGCGTTA |
| AT1G78080-fw | TCTTCCTCCGACGCATCACAAC |
| AT1G78080-rv | TCAGTAACGGCTTTGGGCTGAG |

| | |
|---|---|
| AT1G18330-fw | TCGGTGGACGGATAACTTGCAG |
| AT1G18330-rv | GCTTAGTTCCTCACTTCGATCCTC |
| AT1G45249-fw | CGAGTTACAACGAAAGCAGGCAAG |
| AT1G45249-rv | AGAAGATTCCTCATCTCCGTCTCC |
| AT2G35940-fw | AGTTCTCCGTGCTTGGCTCTTC |
| AT2G35940-rv | GCTTGTCCGAATCCTTAGGGTATG |
| AT3G09600-fw | ACTCTTCGTGGAGCAGAAGCTG |
| AT3G09600-rv | TGAAGCACTGGAGGCTGTTTAGC |
| AT3G15540-fw | TGGTTCGAGCCAAGGCTATGATG |
| AT3G15540-rv | CATCTTTCAAGGCCACACCGATGC |
| AT3G19290-fw | TGGTGGAAATGCAGAAGAATGAGC |
| AT3G19290-rv | TCACCATGGTCCGGTTAATGTCC |
| AT3G23240-fw | TCCCTTCAACGAGAACGACTCAG |
| AT3G23240-rv | AGGTTTGTTGCGTGGACTGCTC |
| AT4G14550-fw | CCAAGCTACGAGGACAAAGATGG |
| AT4G14550-rv | TGCATGACTCGACAAACATCGG |
| AT4G34000-fw | GTTCTCAACCTGCAACACAGTGC |
| AT4G34000-rv | TCCAGGAGATACTGCTGCAACC |
| AT5G13180-fw | CAAAGGCAAACCACCTCATGGC |
| AT5G13180-rv | TCTGAGTGGGACCCATAGAACTCG |
| AT1G18400-fw | TGGTGCCCGGATGTTATAAGGC |
| AT1G18400-rv | GCTGCAGTGAGTTTCATCGAGAGG |
| AT1G73830-fw | TGCTGTGGAATCCATGCAGAAG |
| AT1G73830-rv | ATGGAAGACAGAACTCCCATCCC |
| AT4G36540-fw | TCCAATCTCTGCAACAACAAGTCG |
| AT4G36540-rv | ACCTGTGAAGTAAGCCTGAAACTG |
| **Promoter clone primer** | |
| 35S-attB1 | GGGGACAAGTTTGTACAAAAAAGCAGGCTAGTGCCAAGCTTGGCGTGCCTGCAGG |
| 35S-attB2 | GGGGACCACTTTGTACAAGAAAGCTGGGTATCCTCTCCAAATGAAATGAACTTCC |
| ABRC3-attB1 | GGGGACAAGTTTGTACAAAAAAGCAGGCTCGGCCATGTACGAGCACCGCC |
| ABRC3-attB2 | GGGGACCACTTTGTACAAGAAAGCTGGGTAGCATACCTGTATTATACAGATCC |
| PROM-Glyma02g15780 -FW | CACCTATCATCATCAAATGGCGTGACAC |
| PROM-Glyma02g15780 -RV | ACAATTTAGTCCCTGAGATTGTACC |
| PROM-Glyma04g04310-FW | CACCCAATAAATAGTGACGTACACTGATC |
| PROM-Glyma04g04310-RV | ATTCATAATCATATTCACAGGAGTG |
| PROM-Glyma06g03450-FW | CACCATTTGTGCTTTGCTTAGACTTAGAG |
| PROM-Glyma06g03450-RV | ATATGGAACTATCCCCATGTTTGG |
| PROM-Glyma07g10311-FW | CACCTGCATATTTTCTGTGAATTCAGCC |
| PROM-Glyma07g10311 -RV | TTCCAATTCCCACTAGCTAGAGC |
| PROM-Glyma16g26070 -FW | CACCCAACATTGTCTTGCCCTCAAGGATG |
| PROM-Glyma16g26070 -RV | AACACCAAACACTGAAATACTCCAGG |
| PROM-Glyma05g26930-FW | CACCACTCACACAAATTCAAAGACACAG |
| PROM-Glyma05g26930-RV | GGATAGTTACATTCCTGCCTTAAACG |
| PROM-Glyma08g29100-FW | CACCACTAGCATCACTTGTGTACTAAGGC |

| | |
|---|---|
| PROM-Glyma08g29100-RV | ATCCACCACAATATAACCAGCAGTC |
| PROM-Glyma11g27480-FW | CACCCTTGATTGATTCTAATAACCTCTTC |
| PROM-Glyma11g27480-RV | TGATCCACATAACAACGTACACG |
| PROM-Glyma12g33600-FW | CACCGTTGATGTTGAATTTACCCTAACAC |
| PROM-Glyma12g33600-RV | TCTCTGGGGTATCTGTATCTATTTG |
| **RNA silence fragment** | |
| BRG-siRNA | CACCCTTCAGTTTTGGATGTTTTAACTCAGCAGCTATACAAGACATGAGATCACACTTCATCTGCTCTTTTACTTTAATTTTTCAAAGTTTAAAAGATCAATTTGTTTCCACTATGTGCTACCAGCACATGCAAGTGGACATTCTTGCTTGTACAATCCTTCAGACGTGTAGAAATGAAATGGAAGTTAATAAAAAAAAGGCTATATTTCA |
| ARG-siRNA | CACCAGTAGCTGTGTTGAAAAATGAAAAGCGAAAACGAAAAGGGATTGTTAGTGTGAGCAGAAATTAAGACACTTGGCGCATCCACCACGCCCTTAGAAACCGAGGACGGAGGAACGCAGCCAACCAACTCTCAACTCCCATCATACCAAATCAGGAATACCTAAGTAACATCAAATAAATACCAAACCTTCCTAAG |
